# Better estimation of SNP heritability from summary statistics provides a new understanding of the genetic architecture of complex traits

**DOI:** 10.1101/284976

**Authors:** Doug Speed, David J Balding

## Abstract

LD Score Regression (LDSC) has been widely applied to the results of genome-wide association studies. However, its estimates of SNP heritability are derived from an unrealistic model in which each SNP is expected to contribute equal heritability. As a consequence, LDSC tends to over-estimate confounding bias, under-estimate the total phenotypic variation explained by SNPs, and provide misleading estimates of the heritability enrichment of SNP categories. Therefore, we present SumHer, software for estimating SNP heritability from summary statistics using more realistic heritability models. After demonstrating its superiority over LDSC, we apply SumHer to the results of 24 large-scale association studies (average sample size 121 000). First we show that these studies have tended to substantially over-correct for confounding, and as a result the number of genome-wide significant loci has under-reported by about 20%. Next we estimate enrichment for 24 categories of SNPs defined by functional annotations. A previous study using LDSC reported that conserved regions were 13-fold enriched, and found a further twelve categories with above 2-fold enrichment. By contrast, our analysis using SumHer finds that conserved regions are only 1.6-fold (SD 0.06) enriched, and that no category has enrichment above 1.7-fold. SumHer provides an improved understanding of the genetic architecture of complex traits, which enables more efficient analysis of future genetic data.

---

LD Score Regression (LDSC) has been frequently used to analyze summary statistics from genome-wide association studies (GWAS).^1–4^ It has four main uses: to estimate the average bias due to confounding, to estimate the “SNP heritability” of a trait (the proportion of phenotypic variance explained by all SNPs), to estimate the heritability enrichments of SNP categories, and to estimate the genetic correlations between pairs of traits. LDSC estimates are derived from a specific heritability model in which each SNP in the genome is expected to contribute equally.^1^ Although this model is widely used in statistical genetics, we recently showed that across a range of human traits, it poorly reflects reality.^5^ In particular, it fails to appreciate that in regions of high linkage disequilibrium (LD), the average heritability of each SNP tends to be lower due to multiple tagging of causal variation.^6^ As a result of this model misspecification, LDSC tends to over-estimate confounding bias, under-estimate SNP heritability and produce exaggerated estimates of enrichment.

We propose SumHer, software for estimating SNP heritability from summary statistics that allows the user to specify the heritability model. We apply SumHer to publicly-available GWAS results for 24 disease and quantitative traits,^7^ using a heritability model that we have previously shown to perform well.^5, 6^ We first show that these GWAS tended to over-correct for confounding; when we adjust their results using SumHer, the total number of genome-wide significant loci increases from 1 760 to 2 190. A previous study by Finucane *et al*. used LDSC to estimate enrichments for 24 categories of SNPs defined by functional annotations.^3^ The authors concluded that heritability is highly concentrated in specific functional categories; most notably, they estimated that across 17 diseases, conserved regions contribute 35% of SNP heritability, 13-fold higher than their expected contribution. When we repeat this analysis using SumHer and our 24 traits, the estimated enrichments are more modest: for example, conserved regions are estimated to contribute only 5.8% of SNP heritability and the highest enrichment is only 1.7-fold (transcription start sites), consistent with an omnigenic model of genetic architecture.^8^ We finish by providing an example of how results from SumHer can enable more efficient analysis of genetic data. We show how for body mass index, height, HDL & LDL cholesterol and triglyceride, we are able to significantly improve the predictive performance of polygenic risk scores by incorporating our preferred heritability model and estimates of enrichments. We make SumHer freely available within our software package LDAK (www.ldak.org)^6^

## Results

### SumHer

SumHer has the same four aims as LDSC;^1–3^ we outline them here, with methodological details provided in Online Methods. Suppose we are provided with summary statistics from a GWAS where each of *m* SNPs has been tested individually for association with a particular trait. Suppose also that we have access to a well-matched reference panel, from which we can reliably estimate 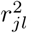, the squared correlation between SNPs *j* and *l*. Let *S*_1_, *S*_2_, …, *S*_*m*_ denote the *χ*^2^(1) test statistics from single-SNP analysis; the first aim is to estimate the average inflation of these test statistics due to confounding. Let 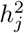 denote the heritability directly contributed by SNP *j*; the second aim is to estimate 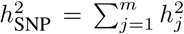, the SNP heritability of the trait.^9^ Let ℂ index a category of SNPs; the third aim is to estimate 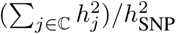, the proportion of SNP heritability contributed by SNPs in ℂ (we can then estimate the enrichment of the category by dividing its estimated proportion of SNP heritability by its expected proportion). Finally, if we are also provided with summary statistics from a second trait, the fourth aim is to estimate the correlation between SNP effect sizes for the two traits.^10^

In order to achieve these four aims, we must specify a heritability model, which describes how 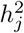 is expected to vary across the genome. Suppose this heritability model takes the form 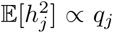 The main difference between LDSC and SumHer is that SumHer allows for any heritability model (i.e., the user can specify arbitrary *q*_*j*_), whereas LDSC assumes all *q*_*j*_ are the same. We recommend using SumHer with the “LDAK Model”: *q*_*j*_ = [*f*_*j*_(1 − *f*_*j*_)]^0.75^ *w*_*j*_, where *f*_*j*_ is the minor allele frequency (MAF) of SNP *j* and *w*_*j*_ is a weighting based on local levels of LD.^5, 6^ In this model, a SNP with high MAF is expected to contribute more heritability than one with low MAF, while a SNP in a region of low LD is expected to contribute more than one in a region of high LD. By contrast, LDSC estimates are obtained by setting *q*_*j*_ = 1, which corresponds to the assumption that all SNPs are expected to contribute equally.^1^ We refer to this as the “GCTA Model” as this is a core assumption of the software GCTA.^5, 9^

A second difference between LDSC and SumHer is how they estimate confounding bias. In a GWAS with no confounding, 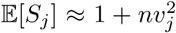, where *n* is the sample size and 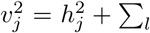 near 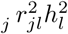 is the heritability tagged by SNP *j* (a working definition of “near” is within 1 Mb^1^). Both LDSC and SumHer estimate the deviation of test statistics from their expected values assuming no confounding. LDSC uses the model 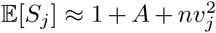, where *A* indicates the average amount each test statistic is inflated additively due to confounding (LDSC then reports 1 + *A*, which it refers to as “the intercept”). By contrast, we recommend using the model 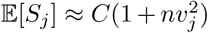, where *C* reflects how much each test statistic is inflated multiplicatively. There are two reasons why we prefer our approach. Firstly, it is standard practice to correct test statistics by scaling (i.e., divide each by *C*);^10^ in theory, one could instead shift test statistics (i.e., subtract *A* from each), but this would result in some negative values. Secondly, the SumHer model for estimating bias accommodates test statistics that have been subjected to genomic control.^11^ Although genomic control is intended to reduce bias, we find that it is often the biggest source of bias in GWAS results and a major hindrance when using summary statistics to interrogate genetic architecture (see below).

In total we use six versions of SumHer, which differ according to their assumed heritability model and allowance for confounding bias. **LDSC-Zero** assumes the GCTA Model and that there is no bias (*A* = 0, *C* = 1); this is equivalent to using the LDSC software^1^ with the option `--intercept-h2 1`. **LDSC** assumes the GCTA Model and allows for additive bias (*A* free to vary, *C* = 1); this is equivalent to using the LDSC software^1^ with default options. **SumHer-Zero** assumes the LDAK Model and that there is no bias (*A* = 0, *C* = 1); this is our recommended version when estimating 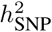 or enrichments and confident that confounding is negligible. **SumHer-GC** assumes the LDAK Model and allows for multiplicative bias (*A* = 0, *C* free to vary); this is our recommended version when estimating confounding or genetic correlations, or when estimating 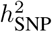 or enrichments and it is likely that test statistics are biased due to population structure or relatedness, or were obtained using genomic control or mixed-model association analysis (see below). **Hybrid-Zero** assumes the heritability model

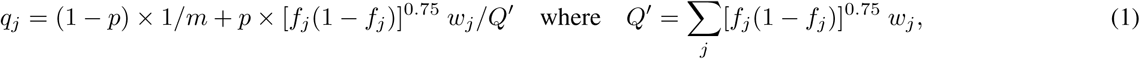

and that there is no bias (*A* = 0, *C* = 1), while **Hybrid-GC** assumes the same heritability model but allows for multiplicative bias (*A* = 0, *C* free to vary). Model (1) is a linear combination of the GCTA and LDAK models, where *p* indicates the weight assigned to the LDAK model; using this heritability model allows us to compare the fit of the GCTA and LDAK models on real data (see below).

For all analyses, our reference panel is 8 850 unrelated Caucasian individuals from the Health and Retirement Study^12^ (HRS). When estimating enrichments of SNP categories, we use the 24 functional annotations used by Finucane *et al*.,^3^ which include coding, conserved, enhancer and promoter regions (see Supplementary Table 1 for a full list). When analyzing real data, we exclude SNPs within the major histocompatibility complex (Chromosome 6: 25-34 Mb), as well as SNPs which individually explain *>*1% of phenotypic variation, and SNPs in LD with these (within 1 cM and 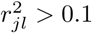).

**Table 1:** Estimates of confounding bias for the 24 summary GWAS. Columns 2 & 3 report the average sample size (*n*) and genomic inflation factor (GIF) for each trait. Columns 4-7 report estimates of 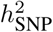 and confounding from both LDSC and SumHer-GC (LDSC measures confounding via the intercept 1 + *A*, while SumHer-GC uses the scaling factor *C*). Columns 8-11 report the number of significant loci based on the published test statistics, then after correction via genomic control, LDSC and SumHer-GC (dividing test statistics by the GIF, 1 + *A* and *C*, respectively). When estimating 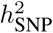, LDSC requires uncorrected test statistics from classical regression. However the test statistics from all 24 GWAS were calculated either using genomic control or mixed-model association analysis; ^*†*^ indicates the 11 traits for which classical regression was used and the impact of genomic control was lowest.

| Trait | $n$ | GIF | LDSC | | SumHer-GC | | Number of significant loci<br>after dividing test statistics by | | | |
| --- | --- | --- | --- | --- | --- | --- | --- | --- | --- | --- |
| | | | $h^2_{\text{SNP}}$ (SD) | $1 + A$ (SD) | $h^2_{\text{SNP}}$ (SD) | $C$ (SD) | 1 | GIF | $1 + A$ | $C$ |
| Alzheimer's Diseases<sup>26</sup> | 54 000 | 1.07 | 0.02 (0.01)<sup>†</sup> | 1.07 (0.01) | 0.09 (0.02) | 1.03 (0.01) | 19 | 17 | 17 | 18 |
| Coronary Artery<sup>27</sup> | 80 000 | 1.07 | 0.03 (0.01) | 1.07 (0.01) | 0.11 (0.01) | 0.99 (0.01) | 11 | 7 | 7 | 11 |
| Crohn's Disease<sup>28</sup> | 21 000 | 1.08 | 0.18 (0.05)<sup>†</sup> | 1.10 (0.02) | 0.67 (0.08) | 0.97 (0.02) | 55 | 51 | 50 | 56 |
| Depression<sup>29</sup> | 161 000 | 1.12 | 0.01 (0.00) | 1.05 (0.01) | 0.06 (0.01) | 0.97 (0.01) | 1 | 0 | 1 | 1 |
| Ever Smoked?<sup>30</sup> | 74 000 | 1.05 | 0.04 (0.00)<sup>†</sup> | 1.03 (0.01) | 0.12 (0.01) | 0.96 (0.01) | 0 | 0 | 0 | 0 |
| Inflammatory Bowel<sup>28</sup> | 35 000 | 1.13 | 0.12 (0.02)<sup>†</sup> | 1.14 (0.02) | 0.49 (0.05) | 0.98 (0.02) | 60 | 51 | 51 | 62 |
| Neuroticism<sup>29</sup> | 171 000 | 1.24 | 0.04 (0.00) | 1.09 (0.01) | 0.16 (0.02) | 0.88 (0.02) | 14 | 5 | 12 | 24 |
| Rheumatoid Arthritis<sup>31</sup> | 58 000 | 1.00 | 0.06 (0.01) | 1.00 (0.01) | 0.26 (0.04) | 0.89 (0.01) | 71 | 71 | 71 | 81 |
| Schizophrenia<sup>32</sup> | 81 000 | 1.35 | 0.22 (0.01)<sup>†</sup> | 1.24 (0.01) | 0.65 (0.03) | 0.90 (0.01) | 82 | 35 | 44 | 108 |
| Subjective-Wellbeing<sup>29</sup> | 298 000 | 1.12 | 0.01 (0.00) | 1.09 (0.01) | 0.05 (0.01) | 0.93 (0.02) | 1 | 0 | 0 | 2 |
| Type 2 Diabetes<sup>33</sup> | 157 000 | 1.08 | 0.03 (0.00) | 1.08 (0.01) | 0.11 (0.01) | 0.95 (0.01) | 31 | 27 | 26 | 33 |
| Ulcerative Colitis<sup>28</sup> | 27 000 | 1.09 | 0.07 (0.02)<sup>†</sup> | 1.11 (0.01) | 0.38 (0.05) | 1.00 (0.01) | 30 | 28 | 27 | 30 |
| Bone Mineral Density<sup>34</sup> | 33 000 | 1.08 | 0.08 (0.01)<sup>†</sup> | 1.07 (0.01) | 0.27 (0.03) | 1.00 (0.01) | 16 | 16 | 16 | 16 |
| Body Mass Index<sup>35</sup> | 229 000 | 0.92 | 0.07 (0.01) | 0.82 (0.01) | 0.28 (0.02) | 0.56 (0.02) | 60 | 70 | 92 | 239 |
| Fasting Glucose<sup>36</sup> | 58 000 | 1.04 | 0.04 (0.01) | 1.05 (0.01) | 0.11 (0.02) | 0.99 (0.01) | 20 | 18 | 18 | 21 |
| Glycated Hemoglobin<sup>37</sup> | 46 000 | 1.02 | 0.02 (0.01)<sup>†</sup> | 1.03 (0.01) | 0.07 (0.02) | 1.00 (0.01) | 8 | 8 | 8 | 8 |
| HDL Cholesterol<sup>38</sup> | 95 000 | 0.95 | 0.08 (0.02) | 1.00 (0.04) | 0.35 (0.04) | 0.74 (0.02) | 99 | 106 | 99 | 142 |
| Height<sup>39</sup> | 245 000 | 1.69 | 0.16 (0.01)<sup>†</sup> | 1.73 (0.05) | 0.38 (0.03) | 1.02 (0.03) | 580 | 242 | 234 | 563 |
| LDL Cholesterol<sup>38</sup> | 90 000 | 0.96 | 0.07 (0.03) | 1.00 (0.04) | 0.36 (0.08) | 0.74 (0.04) | 83 | 89 | 83 | 117 |
| Menarche Age<sup>40</sup> | 253 000 | 1.36 | 0.12 (0.01)<sup>†</sup> | 1.25 (0.02) | 0.28 (0.02) | 0.91 (0.02) | 244 | 134 | 160 | 293 |
| Menopause Age<sup>41</sup> | 69 000 | 1.05 | 0.05 (0.01) | 1.07 (0.02) | 0.21 (0.03) | 0.93 (0.02) | 36 | 31 | 30 | 42 |
| Triglyceride Levels<sup>38</sup> | 92 000 | 0.96 | 0.11 (0.03) | 0.94 (0.03) | 0.34 (0.07) | 0.73 (0.03) | 67 | 71 | 72 | 109 |
| Waist-Hip Ratio<sup>42</sup> | 141 000 | 0.95 | 0.05 (0.00) | 0.93 (0.01) | 0.17 (0.01) | 0.77 (0.01) | 21 | 25 | 28 | 48 |
| Years Education<sup>43</sup> | 329 000 | 1.18 | 0.06 (0.00)<sup>†</sup> | 1.14 (0.01) | 0.17 (0.01) | 0.84 (0.01) | 151 | 88 | 88 | 166 |
| **Average** | **121 000** | **1.11** |  | **1.06 (0.00)** |  | **0.93 (0.00)** | **73** | **50** | **51** | **91** |
| **Total** |  |  |  |  |  |  | **1760** | **1190** | **1234** | **2190** |

### Simulated phenotypes

We first demonstrate the importance of choosing an appropriate heritability model via simulations. For this we use 7 548 unrelated Caucasian individuals from the three control cohorts of the Wellcome Trust Case Control Consortium (WTCCC), recorded for 3 280 768 common SNPs (MAF*>*0.01). We first generate 1 000 phenotypes each with 2 000 causal SNPs and 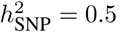; for half the phenotypes, we sample causal SNP effect sizes according to the GCTA Model, for the other half, according to the LDAK Model. We then analyze each phenotype using LDSC-Zero, LDSC, SumHer-Zero and SumHer-GC.

Figure 1a shows that, as expected, accurate estimates of 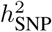 are returned when phenotypes are analyzed assuming the matching heritability model (i.e., when GCTA phenotypes are analyzed using LDSC-Zero or when LDAK phenotypes are analyzed using SumHer-Zero), but that using a different heritability model can result in very poor estimates; SumHer-Zero tends to over-estimate 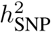 by about 20% when applied to GCTA phenotypes, while LDSC-Zero tends to under-estimate 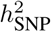 by about 50% when applied to LDAK phenotypes. Supplementary Figure 1a shows LDSC correctly infers that there is no confounding when used on GCTA phenotypes (average intercept 1.000, SD 0.0003), and therefore its estimates of 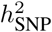 closely match those from LDSC-Zero. However, when used on LDAK phenotypes, LDSC wrongly infers that much of the causal signal is in fact confounding (average intercept 1.033, SD 0.0003), and as a result, its estimates of 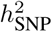 are on average about half those from LDSC-Zero and about 75% lower than the true value.

**Figure 1:**
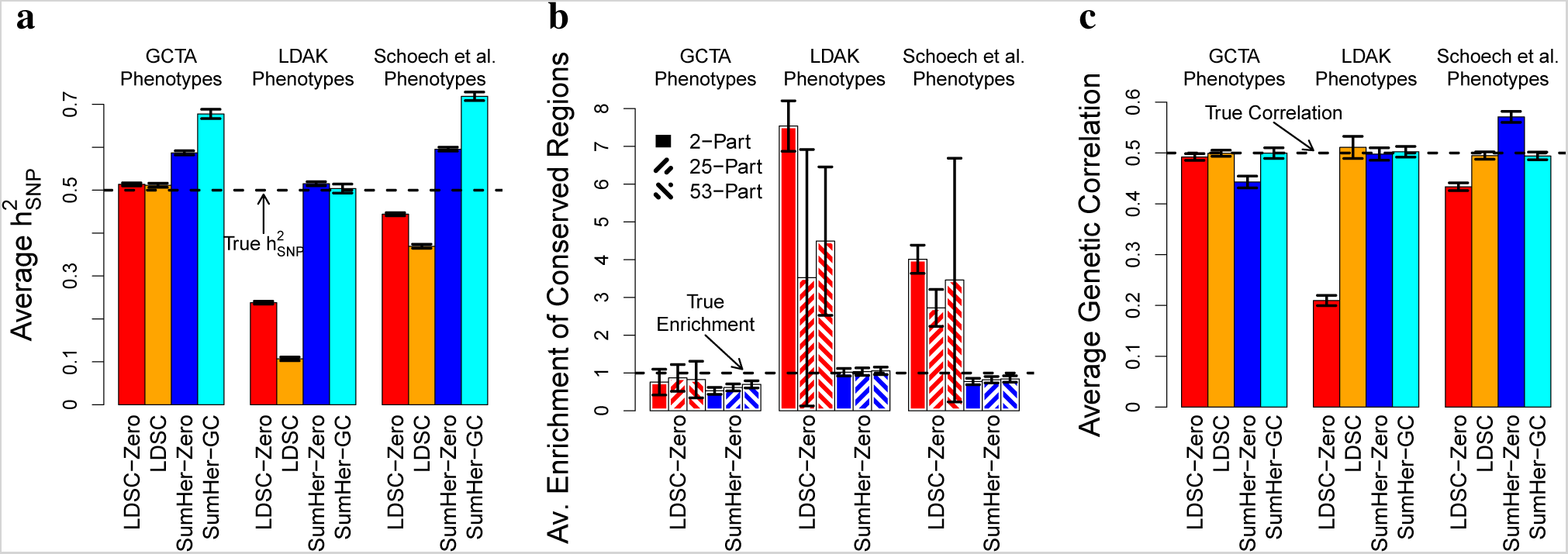
Estimates can depend sensitively on the heritability model. **(a)** We generate 500 phenotypes with 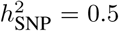 for each of three heritability models, GCTA, LDAK and Schoech *et al*.^13^ (see main text for details of each model). Bars report average estimates of 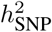 from LDSC-Zero, LDSC, SumHer-Zero and SumHer-GC. **(b)** For the same phenotypes, bars now report average estimates of the enrichment of heritability in conserved regions from LDSC-Zero and SumHer-Zero (true enrichment is 1). For each method, the three bars correspond to estimating enrichment using a 2-part, 25-part or 53-part model (see main text). **(c)** For each of the three heritability models, we now generate 500 additional pairs of phenotypes with genetic correlation 0.5. Bars report average estimates of genetic correlation from LDSC-Zero, LDSC, SumHer-Zero and SumHer-GC. In all plots, vertical line segments mark 95% confidence intervals for the average estimates.

Figure 1b reports the estimated enrichment of SNPs in conserved regions. As causal SNPs were picked at random from across the genome, the true enrichment is 1. Again, we see that assuming the correct heritability model produces reliable estimates, but assuming the wrong model can lead to misleading conclusions. In particular, we find that when LDSC-Zero is used to analyze LDAK phenotypes, it infers that conserved regions are at least 3-fold enriched for heritability. We chose to focus on conserved regions as this was the category that, by applying LDSC to real data, Finucane *et al*.^3^ found to be most enriched. We have previously shown that the LDAK Model better reflects real data than the GCTA Model^5^ (and provide further evidence below), and thus our simulations suggest that a substantial portion of the enrichment observed by Finucane *et al*. is an artifact of misspecifying the heritability model.

Figure 1b also examines how estimates of enrichment are affected by the way the genome is divided into SNP categories (Supplementary Fig. 1c provides a zoomed-in version). The simplest approach is to use a 2-part model, in this case partitioning SNPs into those inside and those outside conserved regions. Next we use a 25-part model, dividing the genome into the 24 functional categories (of which conserved regions are one), plus a category containing all SNPs. Finally, we use a 53-part model, constructed by adding 28 “buffer regions”, the approach recommended by Finucane *et al*. We find that when the correct heritability model is assumed, results appear insensitive to the approach used. However, when the wrong model is assumed, the three approaches can give substantially different estimates.

For Figure 1c, we generate 500 additional pairs of phenotypes, each with genetic correlation 0.5 (again, half the phenotypes are generated under the GCTA Model, half under the LDAK Model). As expected, using LDSC-Zero to analyze GCTA phenotypes or SumHer-Zero to analyze LDAK phenotypes produces accurate estimates of genetic correlation, whereas using LDSC-Zero on LDAK phenotypes or SumHer-Zero on GCTA phenotypes results in biased estimates. Surprisingly, it appears that biases can be minimized by allowing for confounding bias in the analysis (Supplementary Fig. 1f shows why this is the case). However, even if both LDSC and SumHer-GC produce unbiased estimates of genetic correlation, it remains that highest precision is achieved when the correct heritability model is assumed; the SD of LDSC estimates is about half that of SumHer-GC estimates when analyzing GCTA phenotypes, but about twice as high when analyzing LDAK phenotypes.

As well as the GCTA and LDAK phenotypes, for each analysis we additionally generate phenotypes according to a heritability model recently proposed by Schoech *et al*.^13^ We formally define this model in the Discussion, however, loosely speaking, it is intermediate between the GCTA and LDAK models. This is reflected by the estimates of 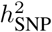 in Figure 1a: on average those from LDSC-Zero are about 5% too low, while those from SumHer-Zero are about 10% too high. We also consider changing the reference panel. While we prefer using the HRS dataset, as its larger sample size enables more accurate estimation of 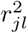, Supplementary Figure 2 shows that estimates are similar if we instead use the 404 non-Finnish Europeans from the 1000 Genomes Project.^14^

**Figure 2:**
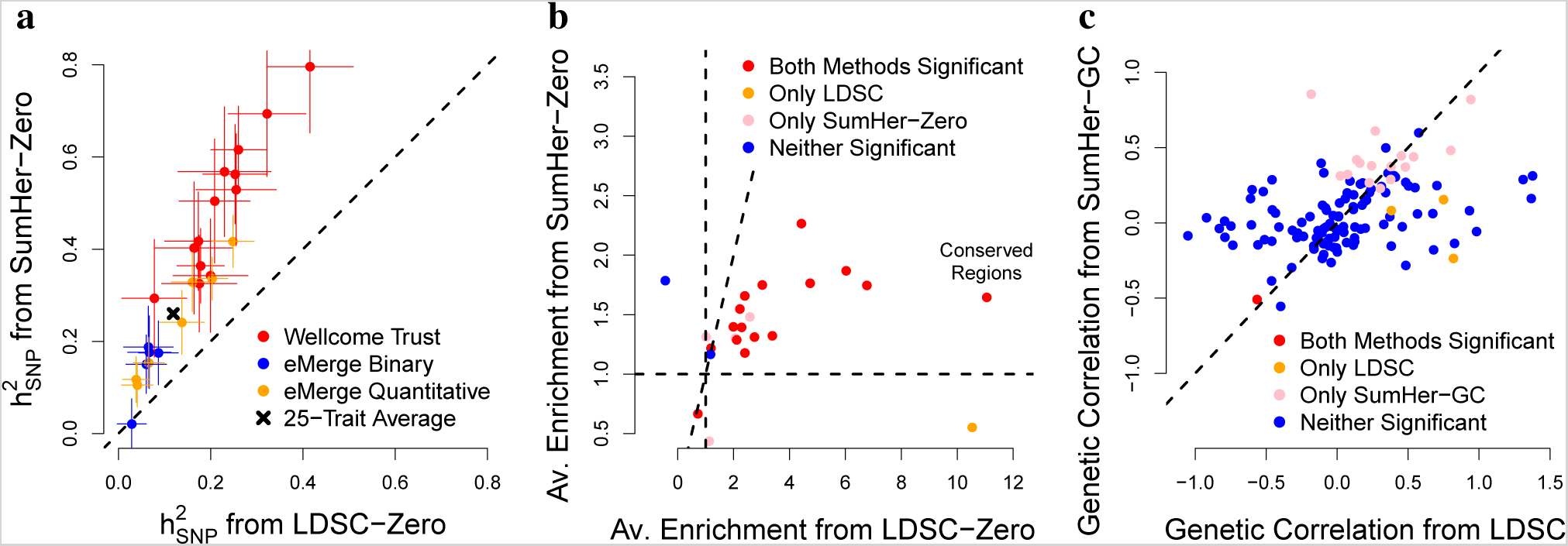
Importance of the heritability model for the 25 raw GWAS. **(a)** Estimates of 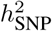 from LDSC-Zero (*x* axis) and SumHer-Zero (*y* axis). Colors distinguish between the 13 WTCCC, the 5 binary eMERGE and 7 quantitative eMERGE traits (black denotes the 25-trait average). Horizontal and vertical line segments mark 95% confidence intervals. Estimates for binary traits are on the observed scale. **(b)** Average estimates of enrichment for the 24 functional categories from LDSC-Zero (using a 53-part model) and SumHer-Zero (using a 25-part model). Colors indicate significant enrichments (*P <* 0.05) from one or both methods. **(c)** Estimates of genetic correlations between pairs of eMERGE traits using LDSC (*x* axis) and SumHer-GC (*y* axis). Colors indicate significant correlations (*P <* 0.05) from one or both methods.

### Real phenotypes

We now show that the choice of heritability model is also important when analyzing real data. We use “25 raw GWAS” (18 binary traits, 7 quantitative, average sample size 9 700; see Supplementary Table 2), for which we have individual-level genotype and phenotype data from either the WTCCC^15^ or the eMERGE Network.^16^ The 13 WTCCC GWAS all examine diseases (e.g., Bipolar Disorder, Ischaemic Stroke and Parkinson’s Disease), while the 12 eMERGE GWAS consider a mixture of diseases (e.g., age-related macular degeneration, heart failure and peripheral artery disease) and clinical measurements (e.g., blood pressure, height and lipid levels). After imputation, strict quality control, and excluding SNPs not present in our reference panel, the WTCCC data contain on average 2.3 M SNPs, while the eMERGE data contain 2 972 162 SNPs. When performing single-SNP analysis, we include as covariates sex and ten principal components (five calculated from the data, five derived from the 1000 Genomes Project^14^). As we have access to raw data, we can use REML to estimate how much of the total phenotypic variance explained by SNPs is inflation due to population structure or relatedness.^17, 18^ We estimate that on average 3.6% of the variance explained is inflation (range -0.9% to 6.8%), indicating that confounding due to population structure and relatedness is modest (Supplementary Fig. 3).

**Figure 3:**
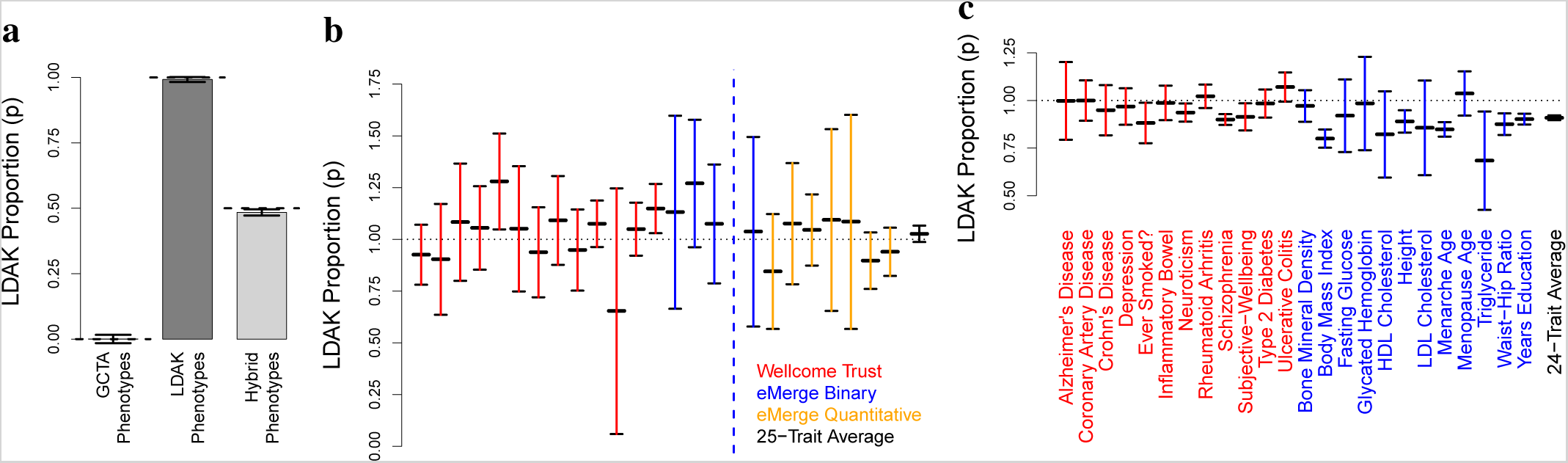
Comparing the GCTA and LDAK Models. These analyses use Hybrid-Zero and Hybrid-GC, versions of SumHer which assign weights 1−*p* and *p* to the GCTA and LDAK heritability models, respectively. **(a)** Estimates of *p* from Hybrid-Zero for GCTA phenotypes (true *p* = 0), LDAK phenotypes (true *p* = 1) and hybrid phenotypes (true *p* = 0.5). **(b)** Estimates of *p* from Hybrid-Zero for the 25 raw GWAS. Colors distinguish between the 13 WTCCC, the 5 binary eMERGE and 7 quantitative eMERGE traits (black denotes the 25-trait average). A precise estimate of *p* was not possible for Shingles (dashed vertical line), due to the trait having very low 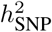. **(c)** Estimates of *p* from Hybrid-GC for the 24 summary GWAS and the 24-trait average. In all plots, vertical line segments mark 95% confidence intervals.

Figure 2a and Supplementary Table 2 show that across the 25 traits, estimates of 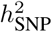 from SumHer-Zero are on average 2.0 times larger (SD 0.05) than those from LDSC-Zero. Figure 2b and Supplementary Table 3 report estimates of enrichment for the 24 functional categories, averaged across the 25 traits. First we estimate enrichment using LDSC-Zero with a 53-part model (the approach taken by Finucane *et al*.^3^), then using SumHer-Zero with a 24-part model (our recommended approach). While there is reasonable agreement between which categories SumHer-Zero and LDSC-Zero declare to be significant, their estimates of enrichment are very different. Notably, LDSC-Zero estimates that conserved regions on average contribute 28% (SD 4) of SNP heritability (corresponding to 11-fold enrichment), whereas SumHer-Zero estimates that they contribute only 5.1% (SD 0.7) of SNP heritability (1.6-fold enrichment). Supplementary Figure 4 compares estimates of enrichment from 2-part, 25-part and 53-part models. When we use LDSC-Zero, we find substantial differences between the results of the three approaches; in particular, not one of the 24 estimates from the 2-part model is consistent (*P >* 0.05*/*24) with either of the corresponding estimates from the 25-way or 53-way models. By contrast, when we use SumHer-Zero, there is much stronger concordance between the three sets of results; for example, all 24 (19 out of 24) of the 2-part model estimates are consistent with those from the 25-way (53-way) model.

**Figure 4:**
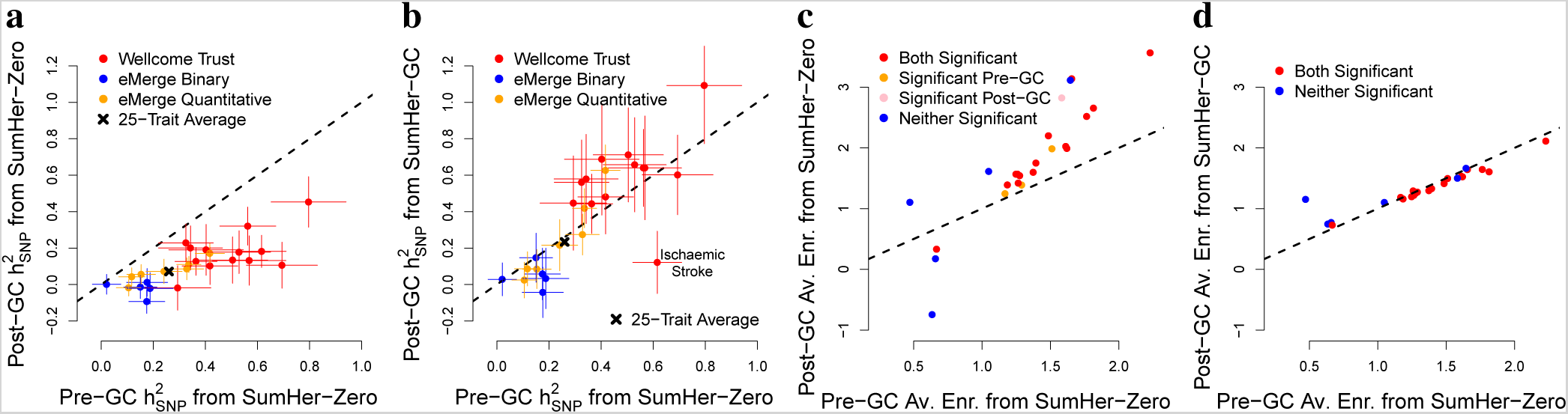
Correcting for genomic control. For the 25 raw GWAS, each plot compares estimates from SumHer using raw test statistics (*x*-axis) with those using test statistics subjected to genomic control (*y*-axis). Horizontal and vertical line segments mark 95% confidence intervals. **(a)** Estimates of 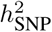 using SumHer-Zero both pre and post genomic control. **(b)** Estimates of 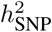 using SumHer-Zero pre genomic control and SumHer-GC post genomic control. **(c)** Estimates of average enrichment for the 24 functional categories using SumHer-Zero both pre and post genomic control. **(d)** Estimates of average enrichment for the 24 functional categories using SumHer-Zero pre genomic control and SumHer-GC post genomic control.

Figure 2c and Supplementary Table 4 report genetic correlations between pairs of traits. In general, it is only possible to get a meaningful estimate when both traits have substantial 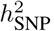, so we restrict to the 18 traits for which both LDSC-Zero and SumHer-Zero find significant 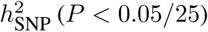. As predicted by our earlier simulations, we find good concordance between the genetic correlations reported by LDSC and SumHer-GC, but that the latter produces more precise estimates: on average the SumHer-GC estimates have SD about half that of the LDSC estimates, and SumHer-GC finds 18 pairs of traits with significant correlation (*P <* 0.05), whereas LDSC finds only 4.

### Comparing heritability models

Previously, we compared different heritability models based on the likelihood from REML analysis; we showed that across 42 human traits (average sample size 7 400), log likelihood was on average 9.8 higher if we assumed the LDAK Model rather than the GCTA Model.^5^ Those 42 traits included the 13 WTCCC traits we study here. If we repeat that analysis using the 12 eMERGE traits (average sample size 13 000), we find that log likelihood is on average 17 higher under the LDAK Model (Supplementary Table 5). In Supplementary Table 6, we show that it remains possible to compare models based on likelihood if only summary statistics are available. However, an alternative, and easier to visualize, method is to fit both the GCTA and LDAK Models simultaneously and allow the data to decide the relative weighting of each. Specifically, we use Hybrid-Zero, with the focus on estimating *p* in Model (1), the “proportion of LDAK” in the heritability model. Figure 3a demonstrates proof of principle: we see that Hybrid-Zero correctly estimates *p* = 0 when applied to GCTA phenotypes, *p* = 1 for LDAK phenotypes, and *p* = 0.5 for “Hybrid Phenotypes” (each created by summing a GCTA and an LDAK phenotype). We then apply Hybrid-Zero to our 25 raw GWAS (Figure 3b and Supplementary Table 6), finding on average *p* = 1.03 (SD 0.02), indicating that the data overwhelmingly support the LDAK Model over the GCTA Model.

### Population structure and relatedness

Until recently, it was standard to estimate confounding bias via the genomic inflation factor^11^ (GIF). However, the GIF tends to over-estimate bias, because it makes the assumption that all observed inflation of test statistics is due to confounding.^19^ LDSC provides a method for estimating confounding bias which appreciates that substantial inflation can instead be due to causal variation.^1^ However, our above simulations indicate that LDSC also tends to over-estimate bias, due to the poor fit of the GCTA heritability model. This is supported by our analysis of the 25 raw GWAS (Supplementary Table 2), for which we performed very careful quality control and verified using REML that confounding due to population structure and relatedness is modest (Supplementary Fig. 3). LDSC finds substantial bias; its average estimate of the intercept (1 + *A*) is 1.031 (SD 0.002), which is similar to the average GIF, 1.035. By contrast, SumHer-GC finds that bias is slight; its average estimate of the scaling factor (*C*) is 1.001 (SD 0.002).

We now construct GWAS when there is substantial confounding. For each of the 13 WTCCC GWAS, we replace 2504 of the controls (on average 67%) with 2504 individuals from POBI^20^ (People of the British Isles); this generates population structure because, although both WTCCC and POBI individuals were recruited from the UK, the latter predominately came from isolated, rural regions (Supplementary Figure 5). Supplementary Table 7 reports estimates of confounding bias and 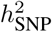 for each GWAS. SumHer-GC now finds substantial bias; its average estimate of the scaling factor is 1.049 (SD 0.003). For comparison, the average GIF is 1.098, while the average LDSC intercept is 1.088 (SD 0.002). SumHer-Zero estimates of 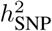 after switching controls are on average 95% (SD 4) higher than SumHer-Zero estimates prior to switching, demonstrating how population structure can lead to substantial inflation when estimating SNP heritability. However, SumHer-GC estimates of 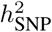 after switching controls are on average only 4% (SD 7) higher, consistent with SumHer-GC being able to reliably estimate SNP heritability despite the population structure.

**Figure 5:**
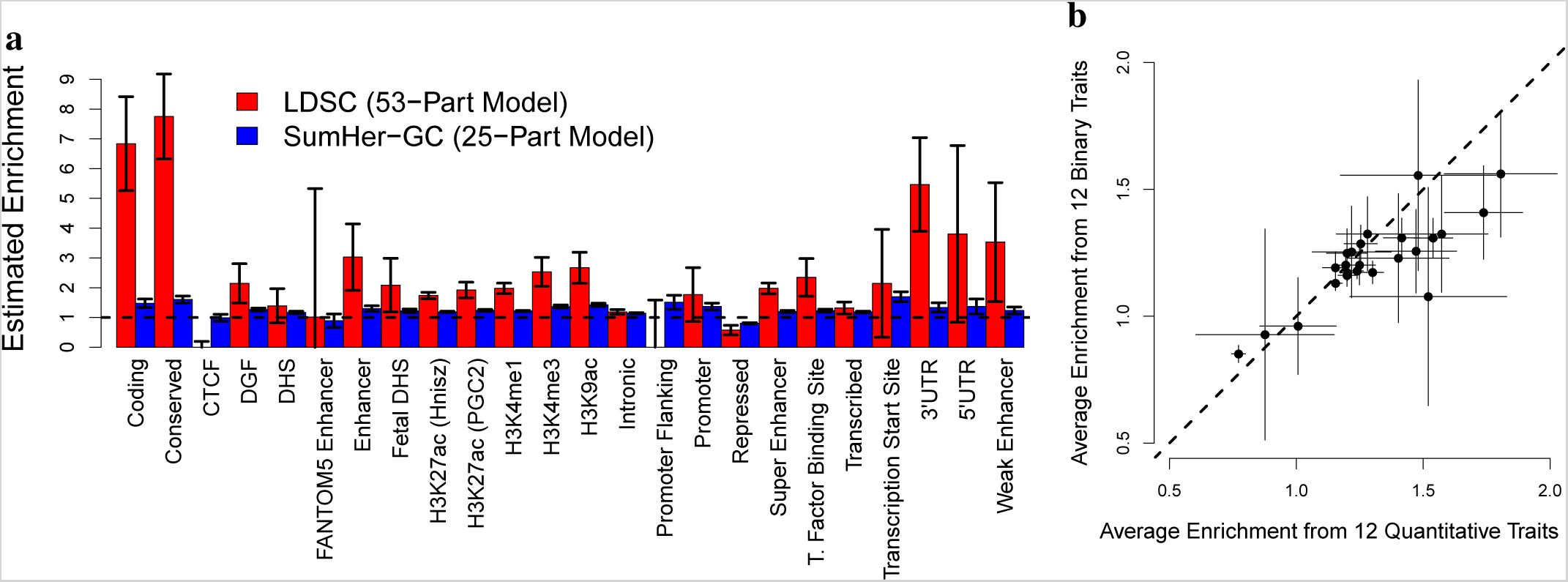
Functional enrichments across the 24 summary GWAS. Both plots show average estimated enrichments for the 24 functional categories. Horizontal and vertical line segments mark 95% confidence intervals. **(a)** Estimates from LDSC using a 53-part model (red bars) and from SumHer-GC using a 25-part model (blue bars). **(b)** Estimates from SumHer-GC using a 25-part model, based either on the 12 quantitative traits (*x*-axis) or on the 12 binary traits (*y*-axis).

### Genomic control and mixed-model association analysis

Although its use is declining, it remains that the majority of published GWAS have performed genomic control (divided test statistics by the GIF) at least once in their analyses.^11^ As the GIF tends to over-estimate confounding,^19^ genomic control tends to produce negatively biased test statistics. For this reason, we recommend using SumHer-GC to estimate 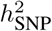 and enrichment, instead of SumHer-Zero, if it is likely that genomic control has been applied. By way of demonstration, we perform genomic control for each of the 25 raw GWAS. Figure 4a shows that if we continue to use SumHer-Zero, estimates of 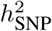 are centered on zero. However, if we instead use SumHer-GC, estimates of 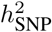 are in general consistent with estimates from SumHer-Zero prior to genomic control (Figure 4b); an exception is stroke, indicative of the LDAK Model being sub-optimal for this trait (Supplementary Figure 6). Similarly, Figures 4c & d show that if genomic control has been performed, it is beneficial to use SumHer-GC instead of SumHer-Zero when estimating enrichments.

**Figure 6:**
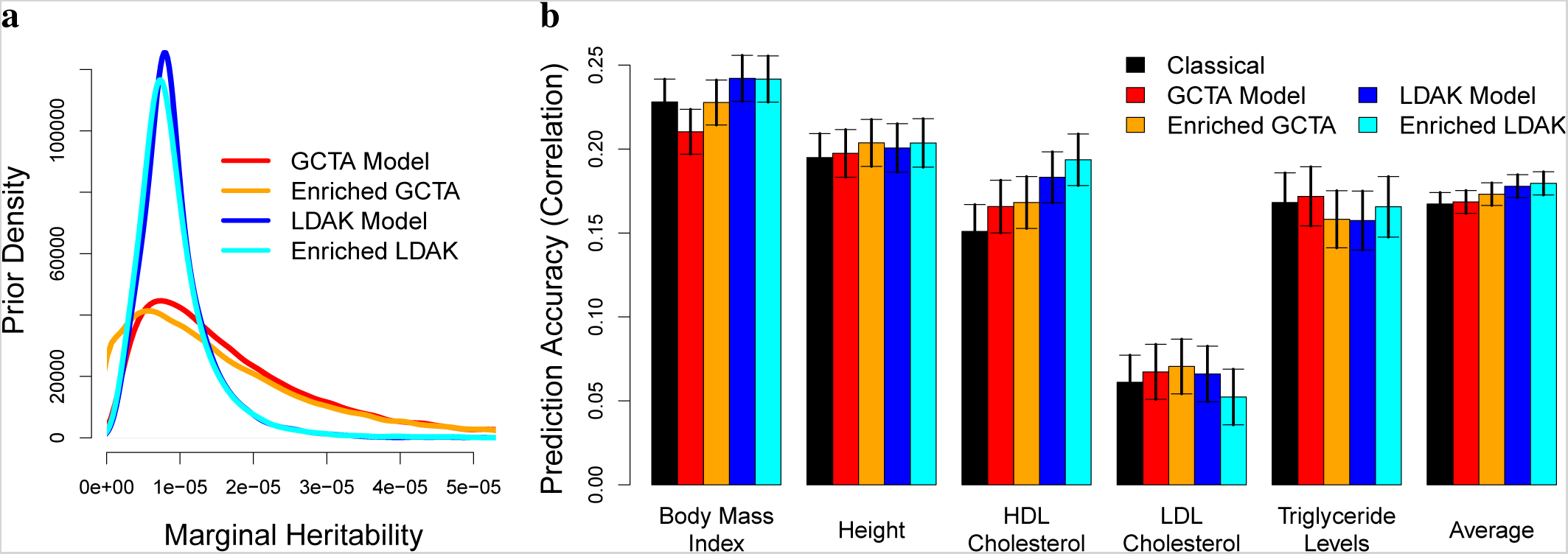
Prediction of five quantitative traits. For each trait, we use data from the 24 summary GWAS to construct Bayesian polygenic risk scores (PRS) corresponding to four heritability models: GCTA, Enriched-GCTA, LDAK and Enriched-LDAK (see main text for details of each model). **(a)** The prior distribution of 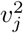, the heritability tagged by SNP *j*, corresponding to each heritability model. **(b)** Prediction accuracy, measured as correlation between observed and predicted phenotypes in the (independent) eMERGE data, for the Classical PRS (effect sizes are frequentist estimates from single-SNP analysis) and for each of the four Bayesian PRS (effect sizes are posterior means). Vertical line segments mark 95% confidence intervals.

SumHer, like LDSC, is designed to be used with test statistics from classical regression (see Online Methods). However, with the development of software such as Fast-LMM, GCTA-LOCO and Bolt-LMM,^21–23^ a popular alternative is for GWAS to use mixed-model association analysis.^24, 25^ Supplementary Figures 6 & 7 show that when estimating 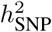 and enrichment, the impact of mixed-model analysis is similar to, albeit less severe than, that of genomic control. For example, SumHer-Zero estimates of 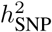 based on mixed-model test statistics are on average about half those based on classical test statistics. However, as with genomic control, reliable estimates can be obtained by using SumHer-GC instead of SumHer-Zero.

## Analysis of published GWAS results

For our main analysis, we use “24 summary GWAS” (12 binary traits, 12 quantitative, average sample size 121 000; see Table 1). These are GWAS for which we do not have individual-level data, but have downloaded summary statistics from previously-published analyses.^7^ After excluding SNPs not present in our reference panel, on average there are summary statistics for 2.2 M SNPs per trait. All of the GWAS either performed genomic control or used mixed-model association analysis, so in general we use SumHer-GC. However, we also repeat all analyses using SumHer-Zero restricted to the 11 traits that did not use mixed-model analysis and for which the impact of genomic control was lowest (Supplementary Table 8).

Figure 3c reports estimates of *p*, the proportion of LDAK in the heritability model, obtained using Hybrid-GC. Across the 24 traits, we estimate *p* = 0.91 (SD 0.01), indicating that again the data strongly support the LDAK Model over the GCTA Model, although not as strongly as for the 25 raw GWAS (see Discussion). Supplementary Table 9 shows that we reach the same conclusion if we instead use Hybrid-Zero restricted to the 11 traits least impacted by genomic control (*p* = 0.90; SD 0.01), or if we compare the GCTA and LDAK Models based on likelihood (the log likelihood is on average 103 higher under the LDAK Model than the GCTA Model).

### Confounding

Table 1 reports the LDSC intercept (1 + *A*) and the SumHer-GC scaling factor (*C*) for each trait. LDSC finds that test statistics are on average inflated by 5.7% (SD 0.2), implying that the GWAS tended to under-correct for confounding. By contrast, SumHer-GC finds that test statistics are on average deflated by 7.4% (SD 0.3), indicating that the studies tended to over-correct. We note that the four GWAS with lowest *C* (0.56 for body mass index, 0.74 for HDL and LDL cholesterol, and 0.73 for triglyceride levels), are all meta-analyses that used genomic control both before and after combining results across cohorts.^35, 38^ Table 1 also reports the number of independent loci with *P <* 5 *×* 10^−8^. Without adjustment (i.e., using the published test statistics), there are on average 73 loci per trait. If we correct using LDSC, this number is reduced to 51, but if we correct using SumHer-GC, it increases to 91. For these counts, we defined two significant SNPs as dependent if they are within 1 cM and have 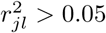, but we find that results are very similar if instead we increase the window size to 3 cM, or use 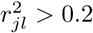 (Supplementary Table 10).

### Functional enrichments

Figure 5a and Supplementary Table 11 report estimates of enrichment for the 24 functional categories, averaged across the 24 traits. We again see striking differences between the estimates from LDSC (using a 53-part model) and those from SumHer-GC (using a 25-part model). For example, LDSC estimates of enrichment range from -1.5 to 7.8, whereas SumHer-GC estimates range from 0.80 to 1.7. Supplementary Table 12 shows that large differences remain if we instead estimate enrichment using LDSC-Zero and SumHer-Zero restricted to the 11 traits least impacted by genomic control. Based on the results from SumHer-GC, we conclude that conserved regions^44, 45^ and transcription start sites^46^ are most enriched for heritability; they are estimated to contribute 5.8% (SD 0.2) and 3.0% (SD 0.2) of 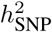, respectively, 1.6 (SD 0.06) and 1.7 (SD 0.08) times higher than their expected contributions under the LDAK Model. Repressed regions^46^ are the only category significantly depleted; although they are estimated to contribute 36% (SD 0.4) of 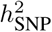, this is 0.80 (SD 0.01) times lower than expected. Further results are provided in Supplementary Figure 8, which confirm that LDSC estimates are similar if we change the SNP sets (LDSC recommends that the reference panel contains as many SNPs as possible, but that only HapMap 3^47^ SNPs with MAF *>* 0.05 are used when performing the regression^1, 3^), or if we use the 75-part model used by Gazal *et al*.^48^

### Genetic correlations

Supplementary Figure 9 and Supplementary Table 13 provide estimates of genetic correlations for the 276 pairs of traits. As expected, there is strong concordance between estimates from LDSC and SumHer-GC, but the SumHer-GC estimates are more precise; for example, across the 41 pairs of traits that both methods find to be significantly correlated (*P <* 0.05*/*276), the SD of SumHer-GC estimates is on average a third lower than the SD of LDSC estimates.

### Improving the efficiency of future analyses

Figure 5b shows that there is strong concordance between the average estimates of enrichment obtained from the 12 binary traits and those from the 12 quantitative traits. This suggests broad similarities between the genetic architectures of different traits, which in turn implies that it should be possible to use information from existing GWAS to improve the efficiency of future analyses. As a demonstration, we consider prediction using polygenic risk scores (PRS). We focus on body mass index, height, HDL & LDL cholesterol and triglyceride levels, as for these five traits we can train prediction models using the 24 summary GWAS, then measure how well these perform on the independent eMERGE data.

To construct a PRS, we need estimates of SNP effect sizes. The current standard is to use estimates from single-SNP analysis (“Classical PRS”). However, in Online Methods, we explain how, given a heritability model, we can obtain a prior distribution for 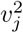, the heritability tagged by SNP *j*, then calculate a “Bayesian PRS” using the posterior mean effect sizes. For each trait, we construct four Bayesian PRS corresponding to four heritability models. First we use the GCTA heritability model (*q*_*j*_ = 1). Next we use the “Enriched GCTA Model”, obtained by scaling the *q*_*j*_ based on the (53-part) estimates of enrichment (e.g., if a category was estimated to have 2-fold enrichment, then the SNPs it contains would have average *q*_*j*_ = 2). We similarly construct PRS based on the LDAK Model, then the “Enriched LDAK Model”, where the *q*_*j*_ are scaled according to the SumHer-GC (25-part) estimates of enrichment.

Figure 6a compares the four prior distributions for 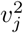. We see the two priors derived from the GCTA Model are more diffuse than the two from the LDAK Model. As explained in the Online Methods, this is because the GCTA Model predicts that 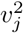 scales with local levels of LD, which vary considerably across the genome, whereas the LDAK Model predicts that 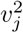 scales with local MAFs, which vary less. Figure 6b and Supplementary Table 14 report the performance of each PRS, measured as correlation between predicted and observed phenotypes for the eMERGE individuals. Averaged across the five traits, the Bayesian PRS constructed from the GCTA and Enriched GCTA Model are, respectively, 1.1% (SD 2.0) worse and 2.3% (SD 2.0) better than the Classical PRS, whereas the Bayesian PRS constructed from the LDAK and Enriched LDAK Model are, respectively, 5.8% (SD 2.0) and 7.5% (SD 2.0) better. The fact that the PRS based on the Enriched GCTA Model outperforms the PRS based on the GCTA Model, is because introducing partitions relaxes the unrealistic assumption that 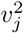 scales with local levels of LD; however, the fact that the PRS based on the LDAK Model outperforms the PRS based on the Enriched GCTA Model, reflects that it is better to start with a more realistic model, than try to fix a sub-optimal model by partitioning.^5^

## Discussion

We have presented SumHer, software for estimating confounding bias, SNP heritability, enrichments of heritability and genetic correlations from GWAS results. While the aims of SumHer are the same as those of LDSC, the key difference is that SumHer allows the user to specify the heritability model. If SumHer is run using the GCTA Model, its estimates will match those from LDSC. However, we instead recommend using the LDAK Model, which we have shown better reflects real data, and therefore produces more accurate estimates. We have analyzed GWAS results for tens of traits, showing that the impact of using an improved heritability model is often substantial, and overall provides a very different description of the genetic architecture of complex traits than has to date been obtained from LDSC analyses.

While the GCTA Model has been used almost exclusively when estimating SNP heritability from summary statistics, alternative models have been used when analyzing individual-level data. A recent submission by Schoech *et al*.,^13^ which has three authors in common with the original LDSC publication,^1^ uses the following model: *q*_*j*_ = [*f*_*j*_(1 − *f*_*j*_)]^0.62^(1 − 0.3*LLD*_*j*_), where *LLD*_*j*_ is a local measure of tagging (obtained by computing LD Scores, binning by MAF, quantile normalizing, then truncating). Like the LDAK Model, the model of Schoech *et al*. assumes that a SNP with high MAF contributes more heritability than one with low MAF, and that a SNP in a region of low LD contributes more than one in a region of high LD. Schoech *et al*. claim that this model is more realistic than both the GCTA Model and the LDAK Model.^13^ However, while they compared their model with the GCTA Model on real data (comparing the two models according to REML likelihood), their only comparison with the LDAK Model was using phenotypes simulated according to their heritability model.^13^ Supplementary Table 5 compares the model of Schoech *et al*. to the GCTA and LDAK Models for the 25 raw GWAS. While we agree with Schoech *et al*. that their model is superior to the GCTA Model (on average, its log likelihood is 9.4 higher), we find it to be inferior to the LDAK Model (on average, its log likelihood is 9.5 lower). Moreover, Figure 1 shows that even if the model of Schoech *et al*. does accurately reflect the genetic architecture of complex traits, it would remain the case that LDSC is producing inaccurate estimates of confounding bias, 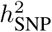 and enrichment.

We note that, despite remaining superior to the GCTA Model, the LDAK Model does not perform as well for the 24 summary GWAS (average LDAK Proportion 0.91) as for the 25 raw GWAS (average 1.02). One possible reason for this is that for the 24 summary GWAS we had to rely on the quality control performed by the original analysts, which was generally much less strict than we would recommend for heritability analysis.^18^ For example, when analyzing the 25 raw GWAS, we restricted to SNPs with imputation information score *>* 0.99, whereas for the 24 summary GWAS, the most common thresholds were 0.5 and 0.8. There will be a correlation between the heritability of a SNP and its genotyping certainty (a SNP genotyped with error will tag less causal variation than were it perfectly typed),^5^ and similarly, there will be a correlation between genotyping certainty and local levels of LD (low-LD regions tend to contain more low-MAF SNPs, which are often hard to genotype reliably, while imputation is easier in high-LD regions). Therefore, including lower-certainty SNPs in a GWAS will generate correlation between the heritability of each SNP and levels of LD, which will result in traits appearing more “GCTA-like”. Alternatively, the results for the 24 summary traits might simply reflect that the LDAK Model is not perfect, and will fit some traits better than others. Therefore, we encourage readers to find ways to improve the LDAK Model, either generally or on a per-trait level, which they can do using the tools provided by SumHer for testing and comparing different heritability models on large-scale GWAS data.

Two practical advantages of SumHer over LDSC are evidenced by the results for height in Table 1 (for these we used summary statistics from the most recent GIANT Consortium meta-analysis,^39^ which has average sample size 245 000). First we focus on estimates of 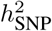. Accurate estimates of 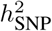 are important not only because they improve our understanding of genetic architecture, but also because they are now being incorporated in software for analyzing complex traits (e.g., the mixed-model association software Bolt-LMM^23^ and the prediction software LDPred^49^). Multiple studies have estimated that common SNPs explain at least 40% of the variation in height;^6, 9, 17, 39, 50^ this indicates that the SumHer-GC estimate of 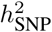 (0.38, SD 0.03) is closer to the truth than the LDSC estimate (0.16, SD 0.01). Complementary to estimates of 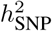 are estimates of confounding bias. These are used both to assess the quality of a GWAS and when correcting test statistics. LDSC estimates the intercept (1 + *A*) to be 1.73 (SD 0.05), implying that there is substantial bias and that strong correction is required; were we to divide test statistics by 1.73, the number of significant loci drops by over half. By contrast, SumHer-GC estimates the scaling factor (*C*) to be 1.02 (SD 0.03), indicating that confounding is slight and that almost no correction of test statistics is needed (dividing the test statistics by 1.02 reduces the number of significant loci by only 3%).

The most striking differences between LDSC and SumHer are observed when estimating heritability enrichments, as highlighted in Figure 5a. Whereas analyses using LDSC have found that heritability is highly focused in specific genomic regions, SumHer instead shows that heritability is spread far more diffusely across the genome, supporting an omnigenic view of genetic architecture.^8^ While the realization that complex traits are even more complicated than previously thought is daunting, as our prediction example demonstrates, it is only by properly understanding their complexity that we can develop more efficient tools for analyzing genetic data.

### URLs

LDAK, http://www.ldak.org/; LDSC, http://www.github.com/bulik/ldsc; 24 functional annotations, https://data.broadinstitute.org/alkesgroup/LDSCORE/baseline_bedfiles.tgz.

### Methods

Methods, including statements of data availability and any associated accession codes and references, are available in the online version of the paper.

## Acknowledgments

We thank Alkes Price, Hilary Finucane, Paul O’Reilly and Maria Speed for helpful discussions. Access to Wellcome Trust Case Control Consortium data was authorized as work related to the project “Genome-wide association study of susceptibility and clinical phenotypes in epilepsy,” access to eMERGE Network data was granted under dbGaP Project 14422, “Comprehensive testing of SNP-based prediction models,” while access to the Health and Retirement Study was granted under dbGaP Project 15139, “Developing summary-statistic tools for analysing genetic association study data.” D.S. is funded by the UK Medical Research Council under grant MR/L012561/1, by the European Unions Horizon 2020 Research and Innovation Programme under the Marie Skłodowska-Curie grant agreement number 754513, and by Aarhus University Research Foundation (AUFF). The eMERGE Network was initiated and funded by NHGRI through the following grants: U01HG006828 (Cincinnati Childrens Hospital Medical Center/Boston Childrens Hospital); U01HG006830 (Childrens Hospital of Philadelphia); U01HG006389 (Essentia Institute of Rural Health, Marshfield Clinic Research Foundation and Pennsylvania State University); U01HG006382 (Geisinger Clinic); U01HG006375 (Group Health Cooperative); U01HG006379 (Mayo Clinic); U01HG006380 (Icahn School of Medicine at Mount Sinai); U01HG006388 (Northwestern University); U01HG006378 (Vanderbilt University Medical Center); and U01HG006385 (Vanderbilt University Medical Center serving as the Coordinating Center). The Health and Retirement Study genetic data is sponsored by the National Institute on Aging (grant numbers U01AG009740, RC2AG036495, and RC4AG039029) and was conducted by the University of Michigan. Analyses were performed with the use of the UCL Computer Science Cluster and the help of the CS Technical Support Group, as well as the use of the UCL Legion High-Performance Computing Facility (Legion@UCL) and associated support services.

## Author contributions

D.S. performed the analysis, D.S. and D.J.B. wrote the manuscript.

## Online Methods

The Supplementary Note provides step-by-step code for using SumHer to estimate SNP heritability, confounding bias, heritability enrichments and genetic correlations from summary statistics.

### Estimating SNP heritability

Suppose that we have summary statistics from a GWAS on *n* individuals and *m* SNPs; let *S*_*j*_ denote the *χ*^2^(1) test statistic from regressing the phenotype on *X*_*j*_, the vector of additively-coded genotypes for SNP *j*, and let *n*_*j*_ ≤ *n* denote the number of individuals used in this regression (note that in the main text, for simplicity, we assumed *n*_*j*_ = *n*). If *S*_*j*_ was obtained using classical (i.e., least-squares) linear regression, then^19^

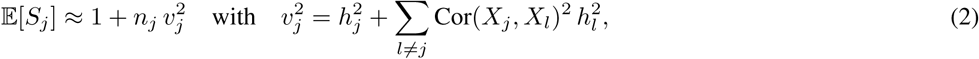

where 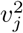 is the total amount of heritability tagged by SNP *j*. In the main text, we referred to 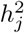 as the heritability “directly contributed” by SNP *j*, to emphasize that while a causal variant can contribute to multiple 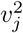 (i.e., be tagged by multiple SNPs), it can only contribute to one 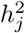. More formally, the 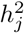 represent a partitioning of 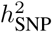, the total heritability tagged by the *m* SNPs genotyped by the GWAS; this formal definition appreciates that a causal variant need not be typed to contribute towards 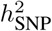, provided it is tagged by one or more SNPs that have been typed (in which case its heritability will be shared across the 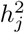 of the tagging SNPs, even though none of these “directly contribute” this heritability). If there is no population structure or cryptic relatedness, then Cor(*X*_*j*_, *X*_*l*_)^2^ will be negligible for distant SNPs, while for local SNPs, an unbiased estimate of Cor(*X*_*j*_, *X*_*l*_)^2^ is ^1^

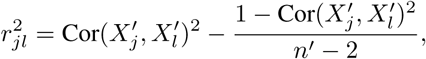

where 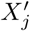 is the vector of SNP *j* genotypes for the *n*′ individuals in a reference panel (for accurate estimates of *r*_*jl*_, these individuals should have similar ancestry to those used in the GWAS). Therefore, in place of Equation (2) we use

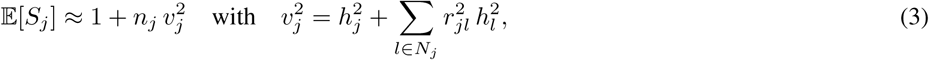

where the set *N*_*j*_ indexes those SNPs “near” SNP *j*; a working definition of near is two SNPs within 1 cM (Supplementary Figure 2). Finally, given a heritability model defined as 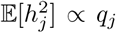, we replace 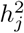 by its expected value 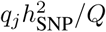, where 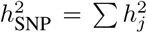 and *Q* = Σ_*j*_ *q*_*j*_, resulting in

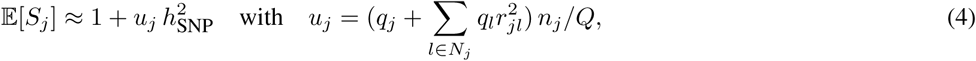

which allows us to estimate 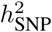 by regressing (*S*_*j*_ − 1) on *u*_*j*_. To account for correlated datapoints and heteroscedasticity we use weighted least squares regression. Specifically, if *D* is a diagonal matrix whose non-zero entries are the regression weights, then the estimate of 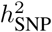 would be (*u*^*T*^ *Du*)^−1^*u*^*T*^ *D*(*S* − 1), where *u* and *S* are vectors containing the *m* values for *u*_*j*_ and *S*_*j*_, respectively. Following LDSC,^1^ we use 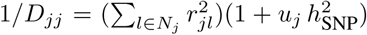. Since 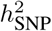 is unknown, we proceed iteratively starting at 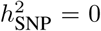 then successively updating *D*_*jj*_ and 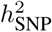 until convergence. Again following LDSC, we estimate standard errors via block jackknifing (by default we use 200 blocks).

### Comparing heritability models

The (weighted) log likelihood is

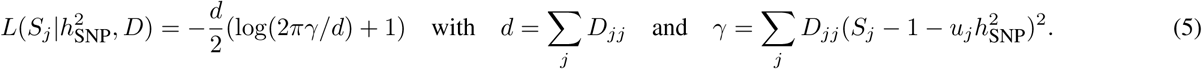

To evaluate 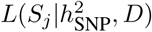 requires values for 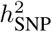 and *D*. While the natural choice is to use the values after the final iteration, in order to compare different heritability models based on 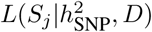, we must use the same *D* for each (else models leading to lower *D*_*jj*_ would have an unfair advantage). Therefore, SumHer reports 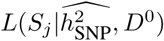, where 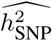 is the final estimate of 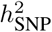, but *D*^*0*^ is the initial weight matrix (obtained by setting 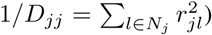); Supplementary Table 6 shows that for the 25 raw GWAS, comparisons of heritability models based on 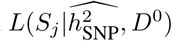 align closely with those based on REML likelihood.

### Estimating confounding bias

We recommend replacing Equation (3) with 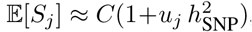, where *C* denotes the multiplicative inflation of test statistics due to confounding; we can then estimate *C* and 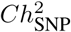 by jointly regressing *S*_*j*_ on 1 and *u*_*j*_. To instead copy LDSC, which considers the additive inflation of test statistics, we replace Equation (4) with 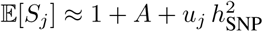, then estimate *A* and 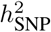 by jointly regressing (*S*_*j*_ − 1) on 1 and *u*_*j*_.

### Estimating enrichments

Suppose we have *K* categories; let *I*_*jk*_ ∈ *{*0, 1*}* indicate whether SNP *j* belongs to Category *k*. We wish to estimate 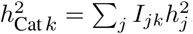, the heritability contributed by SNPs in Category *k*. We now use the heritability model

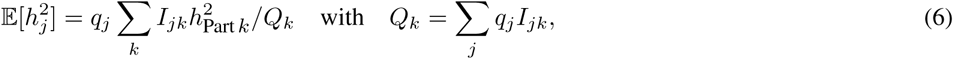

where 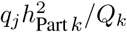 indicates the contribution of Category *k* to the expected heritability of SNP *j*, and 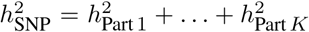. We consider two scenarios: in Scenario 1, the *K* categories partition the genome (_*k*_ *I*_*jk*_ = 1); in Scenario 2, the first *K* − 1 categories correspond to annotations, and the *K*th is the base category containing all SNPs (*I*_*jK*_ = 1). Using Equation (6) ensures that when there is no enrichment, 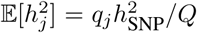. To appreciate why, consider that for Scenario 1, 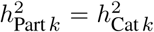 so that when no categories are enriched, 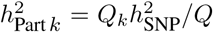; for Scenario 2, 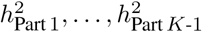 indicate how much each annotation increases the SNP heritability from 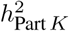, so that when there is no enrichment, 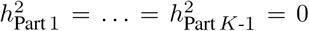 and 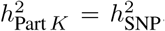 Now when we replace 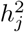 in Equation (3) by its expected value, we obtain

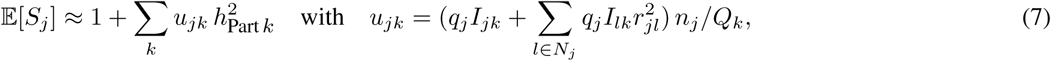

and therefore we can estimate 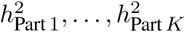 by jointly regressing (*S*_*j*_ − 1) on *u*_*j*,1_, …, *u*_*jk*_. Given these, our estimate of 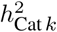 is 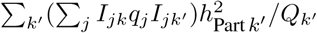, which we then divide by 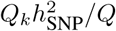 to get an estimate of the enrichment of Category *k*.

### Estimating genetic correlation

Suppose we have summary statistics from two GWAS. Instead of *χ*^2^(1) test statistics, we now use (signed) *Z*-statistics. Let *Z*_*Aj*_ and *Z*_*Bj*_ denote the two *Z*-statistics for SNP *j*, computed using *n*_*Aj*_ and *n*_*Bj*_ individuals, respectively, of which *n*_*Cj*_ were common to both GWAS (if the two GWAS were independent, *n*_*Cj*_ = 0). We assume

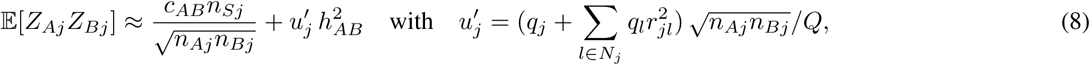

where *c*_*AB*_ is the phenotypic correlation between the two traits and 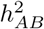 is their genetic covariance. This equation matches that used by LDSC,^2^ except we have replaced 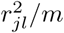 by 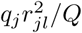 By regressing *Z*_*Aj*_*Z*_*Bj*_ on 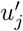, we obtain an estimate of 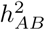, which we then divide by estimates of 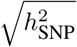 for each trait to produce an estimate of their genetic correlation.

### Regions of extreme LD and large-effect SNPs

Estimates of SNP heritability can be unduly affected by regions of extreme LD and by SNPs with disproportionately large effect size.^4, 5^ Therefore, for all analyses, we exclude SNPs within the major histocompatibility complex (Chromosome 6: 25-34 Mb), as well as SNPs which individually explain *>*1% of phenotypic variation (*S*_*j*_ *> n*_*j*_*/*99), and SNPs in LD with these (within 1 cM and 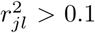); the latter resulted in the exclusion of SNPs for 8 of the 25 raw traits and for 6 of the 24 summary traits (Supplementary Table 15).

### Binary phenotypes

So far, we have implicitly assumed the trait is quantitative. If instead it is binary (i.e., for case-control GWAS), there are two considerations. Firstly, heritability estimates now correspond to the “observed scale;” SumHer will convert them to the “liability scale” if provided with the prevalence and ascertainment.^51, 52^ Secondly, it is likely the *p*-values came from (classical) logistic regression instead of linear regression; however, Supplementary Figure 10 shows that, because linear and logistic *p*-values closely match for SNPs with small or moderate effect, this tends to have limited impact.

### Polygenic risk scoring

To predict phenotypes, we construct PRS of the form 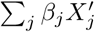, where the vector 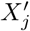 contains standardized genotypes for SNP *j* (obtained by centering *X*_*j*_, then scaling to have variance 1). Without loss of generality, we assume phenotypes have also been standardized, in which case the estimate of *β*_*j*_ from classical linear regression is *ρ*_*j*_, the correlation observed between SNP *j* and the phenotype, and has variance 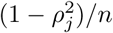. Note that given summary statistics, we can recover *ρ*_*j*_ by appreciating that the Wald test *Z*-statistic is 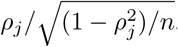. For the Classical PRS, our estimate of *β*_*j*_ is *ρ*_*j*_. For the Bayesian PRS, we must specify a prior distribution for *β*_*j*_. Given a heritability model, we use 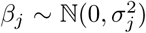, where 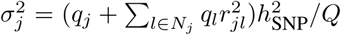; this prior is motivated by recognizing that for the “true PRS”, 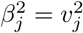, the heritability tagged by SNP *j*. We then estimate *β*_*j*_ by its posterior mean, a shrunken version of the classical estimate. To calculate this, we approximate the likelihood distribution by 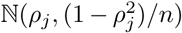,^53^ then the posterior mean equals 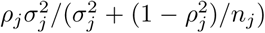. Two technicalities. Firstly, to construct each Bayesian PRS requires a value for 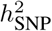, so we used the corresponding estimate from LDSC-Zero or SumHer-Zero (for Enriched GCTA we used the 53-part model, for Enriched LDAK we used the 25-part model). Supplementary Table 14 confirms that the ranking of PRS remains the same if we instead agnostically set 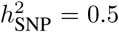. Secondly, for the Bayesian PRS, we performed clumping^54^ (thinned SNPs to ensure no pair within 1 cM had 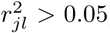), whereas for the Classical PRS we did not; this was because we found clumping benefited all Bayesian PRS, but was detrimental to the Classical PRS. Supplementary Table 14 shows that the ranking of Bayesian PRS remains the same if we do not clump, but then they are inferior to the Classical PRS.

### Choosing a heritability model

Although the heritability model is defined in terms of 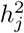, as is clear from Equation (3), its fit depends only on how well it models 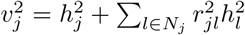. Therefore, when specifying *q*_*j*_, the focus should be on accurately describing how 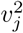 is expected to vary across the genome. LDSC assumes 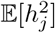 is constant (*q*_*j*_ = 1), and therefore that 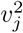 correlates with local levels of LD; we refer to this as the GCTA Model^5^ because the same assumption is made by GCTA, software for estimating SNP heritability via REML.^55^ The GCTA Model is widely used in statistical genetics. For example, it is implicitly assumed by any regression method where SNPs are standardized and the same penalty function / prior distribution is applied to each, and likewise by any simulation study where causal SNPs are picked at random and their standardized effect sizes are sampled from a common distribution. However, as we have shown, the GCTA Model poorly reflects real data.^5^ Even for traits where a correlation is observed between 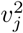 and *S*_*j*_ (those with *p <* 1 in Figures 3b & 3c), the correlation is substantially weaker than predicted by the GCTA Model (hence *p >>* 0), and as we discussed above, might be a consequence of including lower-certainty SNPs in the GWAS. We currently recommend using the LDAK Model. Originally, this took the form *q*_*j*_ = *w*_*j*_, where the weights *w*_*j*_ are calculated based on the assumption that 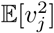 is constant.^6^ We recently updated it to *q*_*j*_ = *w*_*j*_ [*f*_*j*_(1 − *f*_*j*_)]^0.75^ *r*_*j*_, reflecting that 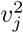 tends to depend on MAF, *f*_*j*_, and genotype certainty, *r*_*j*_ (the latter is not relevant in this study as either *r*_*j*_ ≈ 1 or was unknown), and we recognize that further improvements will be possible.

### Quality control and association analysis

To prepare the 13 WTCCC GWAS, we used our previously described protocol.^5^ In summary, after excluding apparent population outliers, samples with extreme missingness or heterozygosity and SNPs with MAF *<*0.01, call rate *<*0.95 or Hardy-Weinberg *P <* 1 *×* 10^−6^, we phased using SHAPEIT^56^, then imputed using IMPUTE2^57^ with the 1000 Genomes Project Phase 3 (2014) reference panel.^14^ When merging case and control datasets, we converted genotype probabilities to hard calls using a certainty threshold of 0.95, then retained only autosomal SNPs that in all cohorts had MAF ≥ 0.01 and info score ≥ 0.99 (using IMPUTE2 `r2\_type2` for directly genotyped SNPs). Finally, we thinned individuals, so no pair remained with estimated relatedness *>*0.05. We performed the same quality control for the our reference panel, the Health and Retirement Study.^12^ The emerge data were provided post-imputation; this was also performed using SHAPEIT and IMPUTE2, but used the 1000 Genomes Project Phase 2 reference panel.^16^ We converted genotype probabilities to hard calls using a certainty threshold of 0.95, then retained only biallelic SNPs with MAF ≥0.01, call rate ≥0.95, info score ≥0.99 and whose genomic position matched that in the 1000 Genomes Project. Finally, we excluded individuals ancestrally inconsistent (*P <* 0.05) with non-Finnish Europeans from the 1000 Genomes Project (see Supplementary Figure 11) and those whose ethnicity was reported as “Hispanic or Latino”, then filtered until no pair remained with estimated relatedness *>* 0.05 (which left 25 875 individuals).

For the 25 raw GWAS, we performed the association analysis using linear regression (regardless of whether the trait was quantitative or binary), including as covariates sex and ten principal components (five derived from the reference panel, five from the 1000 Genomes Project^14^). For the 24 summary GWAS, we used publicly-available summary statistics (Supplementary Table 8). For each SNP, SumHer requires the two alleles, the *χ*^2^(1) test statistic, *n*_*j*_ and, when estimating genetic correlations, the direction of effect (relative to the first allele). Given summary statistics, we generally used all SNPs in common with our reference panel (and with consistent alleles), however, for the five GWAS which provided info scores, we also excluded SNPs with score *<*0.95. Per-SNP sample sizes were only available for eight of the summary GWAS, so for the remainder, we set *n*_*j*_ = *n*, the total sample size.

### Run times

Given summary statistics and a reference panel, a SumHer analysis has two steps: the first generates a “tagfile”, which contains 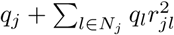 for each SNP; the second performs the regression. The latter is trivial, and typically finishes within minutes. The time to compute the tagfile depends mainly on the size of the reference panel; using our preferred reference panel (8 850 individuals) takes ∼1 day on a single CPU, whereas using the non-Finnish Europeans from the 1000 Genomes Project^14^ (404 individuals) takes a couple of hours. While we recommend users compute tagfiles from scratch, starting with the subset of SNPs common to both the GWAS and reference panel, we alternatively provide a pre-computed tagfile, which should suffice when the GWAS coverage is high.

## References

1. Bulik-Sullivan, B. et al. LD score regression distinguishes confounding from polygenicity in genome-wide association studies. Nat. Genet. 47, 291–295 (2014).

2. Bulik-Sullivan, B. et al. An atlas of genetic correlations across human diseases and traits. Nat. Genet. 47, 1236–1241 (2015).

3. Finucane, H. et al. Partitioning heritability by functional annotation using genome-wide association summary statistics. Nat. Genet. 47, 1228–1235 (2015).

4. Zheng, J. et al. LD Hub: a centralized database and web interface to perform ld score regression that maximizes the potential of summary level GWAS data for SNP heritability and genetic correlation analysis. Bioinformatics 33, 272–279 (2016).

5. Speed, D. et al. Reevaluation of SNP heritability in complex human traits. Nat. Genet. 49, 986–992 (2017).

6. Speed, D., Hemani, G., Johnson, M. & Balding, D. Improved heritability estimation from genome-wide SNP data. Am. J. Hum. Genet. 91, 1011–1021 (2012).

7. Pasaniuc, B. & Price, A. Dissecting the genetics of complex traits using summary association statistics. Nat. Rev. Genet. 18, 117–127 (2017).

8. Boyle, E., Li, Y. & Pritchard, J. An expanded view of complex traits: From polygenic to omnigenic. Cell 169, 1177–1186 (2017).

9. Yang, J. et al. Common SNPs explain a large proportion of the heritability for human height. Nat. Genet. 42, 565–569 (2010).

10. Cross-Disorder Group of the Psychiatric Genomics Consortium. Genetic relationship between five psychiatric disorders estimated from genome-wide SNPs. Nat. Genet. 45, 984–994 (2013).

11. Devlin, B. & Roeder, K. Genomic control for association studies. Biometrics 55, 997–1004 (1999).

12. Juster, F. & Suzman, R. An overview of the Health and Retirement Study. J. Hum. Resources 30, S7–S56 (1995).

13. Schoech, A. et al. Quantification of frequency-dependent genetic architectures and action of negative selection in 25 UK Biobank traits (2017). Preprint available on BioRχiv.

14. The 1000 Genomes Project Consortium. A map of human genome variation from population-scale sequencing. Nature 467, 1061–1073 (2010).

15. The Wellcome Trust Case Control Consortium. Genome-wide association study of 14,000 cases of seven common diseases and 3,000 shared controls. Nature 447, 661–678 (2007).

16. Verma, S. et al. Imputation and quality control steps for combining multiple genome-wide datasets. Front. Genet. 5, 370 (2015).

17. Yang, J. et al. Genomic partitioning of genetic variation for complex traits using common SNPs. Nat. Genet. 43, 519–525 (2011).

18. Speed, D. et al. Describing the genetic architecture of epilepsy through heritability analysis. Brain 137, 26802689 (2014).

19. Yang, J. et al. Genomic inflation factors under polygenic inheritance. Eur. J. Hum. Genet. 19, 807–812 (2011).

20. Leslie, S. et al. The fine-scale genetic structure of the British population. Nature 519, 309–314 (2015).

21. Lippert, C. et al. FaST linear mixed models for genome-wide association studies. Nat. Methods 8, 833–835 (2011).

22. Yang, J., Zaitlen, N., Goddard, M., Visscher, P. & Price, A. Advantages and pitfalls in the application of mixed-model association methods. Nat. Genet. 46, 100–106 (2014).

23. Loh, P. et al. Efficient Bayesian mixed-model analysis increases association power in large cohorts. Nat. Genet. 47, 284–290 (2015).

24. Yu, J. et al. A unified mixed-model method for association mapping that accounts for multiple levels of relatedness. Nat. Genet. 38, 203–208 (2006).

25. The International Multiple Sclerosis Genetics Consortium et al. Genetic risk and a primary role for cell-mediated immune mechanisms in multiple sclerosis. Nature 476, 214–219 (2011).

26. Lambert, J. et al. Meta-analysis of 74,046 individuals identifies 11 new susceptibility loci for Alzheimer’s disease. Nat. Genet. 45, 1452–1458 (2013).

27. Schunkert, H. et al. Large-scale association analysis identifies 13 new susceptibility loci for coronary artery disease. Nat. Genet. 43, 333–338 (2011).

28. Liu, J. et al. Association analyses identify 38 susceptibility loci for inflammatory bowel disease and highlight shared genetic risk across populations. Nat. Genet. 47, 979–986 (2015).

29. Okbay, A. et al. Genetic variants associated with subjective well-being, depressive symptoms, and neuroticism identified through genome-wide analyses. Nat. Genet. 48, 626–633 (2016).

30. The Tobacco and Genetics Consortium. Genome-wide meta-analyses identify multiple loci associated with smoking behavior. Nat. Genet. 42, 441–447 (2010).

31. Okada, Y. et al. Genetics of rheumatoid arthritis contributes to biology and drug discovery. Nature 506, 376–381 (2014).

32. Schizophrenia Working Group of the Psychiatric Genomics Consortium. Biological insights from 108 schizophrenia-associated genetic loci. Nature 511, 421–427 (2014).

33. Scott, R. et al. An expanded genome-wide association study of type 2 diabetes in Europeans. Diabetes 66, 2888–2902 (2017).

34. Zheng, H. et al. Whole-genome sequencing identifies EN1 as a determinant of bone density and fracture. Nature 526, 112–117 (2015).

35. Locke, A. et al. Genetic studies of body mass index yield new insights for obesity biology. Nature 518, 197–206 (2015).

36. Manning, A. et al. A genome-wide approach accounting for body mass index identifies genetic variants influencing fasting glycemic traits and insulin resistance. Nat. Genet. 44, 659–669 (2012).

37. Soranzo, N. et al. Common variants at 10 genomic loci influence hemoglobin a(c) levels via glycemic and nonglycemic pathway. Diabetes 59, 3229–3239 (2010).

38. Global Lipids Genetics Consortium. Discovery and refinement of loci associated with lipid levels. Nat. Genet. 45, 1274–1283 (2013).

39. Wood, A. et al. Defining the role of common variation in the genomic and biological architecture of adult human height. Nat. Genet. 46, 1173–1186 (2014).

40. Perry, J. et al. Parent-of-origin-specific allelic associations among 106 genomic loci for age at menarche. Nature 514, 92–97 (2014).

41. Day, F. et al. Large-scale genomic analyses link reproductive aging to hypothalamic signaling, breast cancer susceptibility and brca1-mediated dna repair. Nat. Genet. 47, 1294–1303 (2015).

42. Shungin, D. et al. New genetic loci link adipose and insulin biology to body fat distribution. Nat. Genet. 518, 187–196 (2015).

43. Okbay, A. et al. Genome-wide association study identifies 74 loci associated with educational attainment. Nature 533, 539–542 (2016).

44. Lindblad-Toh, K. et al. A high-resolution map of human evolutionary constraint using 29 mammals. Nature 478, 476–482 (2011).

45. Ward, L. & Kellis, M. Evidence of abundant purifying selection in humans for recently acquired regulatory functions. Science 337, 1675–1678 (2012).

46. Hoffman, M. et al. Integrative annotation of chromatin elements from encode data. Nucleic Acids Res. 41, 827–841 (2013).

47. The International HapMap 3 Consortium. Integrating common and rare genetic variation in diverse human populations. Nature 467, 52–58 (2010).

48. Gazal, S. et al. Linkage disequilibrium-dependent architecture of human complex traits shows action of negative selection. Nat. Genet..49 (2017).

49. Vilhjálmsson, B. et al. Modeling linkage disequilibrium increases accuracy of polygenic risk scores. Am. J. Hum. Genet 97, 576–592 (2015).

50. Zhou, X., Carbonetto, P. & Stephens, M. Polygeneic modeling with Bayesian sparse linear mixed models. PLoS Genet. 9, e1003264 (2013).

51. Dempster, E. & Lerner, I. Heritability of threshold characters. Genetics 35, 212–236 (1950).

52. Lee, S., Wray, N., Goddard, M. & Visscher, P. Estimating missing heritability for disease from genome-wide association studies. Am. J. Hum. Genet. 88, 294–305 (2011).

53. Wakefield, J. Bayes factors for genome-wide association studies: comparison with p-values. Genet. Epidemiol. 33, 79–86 (2009).

54. Euesden, J., Lewis, C. & O’Reilly, P. PRSice: polygenic risk score software. Bioinformatics 31, 1466–1468 (2015).

55. Yang, J., Lee, S., Goddard, M. & Visscher, P. GCTA: a tool for genome-wide complex trait analysis. Am. J. Hum. Genet. 88, 76–82 (2011).

56. Delaneau, O., Zagury, J. & Marchini, J. Improved whole-chromosome phasing for disease and population genetic studies. Nat. Methods 10, 5–6 (2013).

57. Howie, B., Marchini, J. & Stephens, M. Genotype imputation with thousands of genomes. G3 1, 457–470 (2011).

