## Supplementary Materials for "Better estimation of SNP heritability from summary statistics provides a new understanding of the genetic architecture of complex traits"

**Supplementary Note: Instructions for using SumHer**

**Supplementary Figure 1: Simulations demonstrating the importance of the assumed heritability model**

**Supplementary Figure 2: Choice of reference panel and varying the LD window size**

**Supplementary Figure 3: REML estimates of confounding for the 25 raw GWAS**

**Supplementary Figure 4: Comparing 2-part, 25-part and 53-part models for estimating enrichment for the 25 raw GWAS**

**Supplementary Figure 5: Introducing population structure for the 13 WTCCC GWAS**

**Supplementary Figure 6: Simulations investigating the impact of genomic control and mixed model association analysis**

**Supplementary Figure 7: Mixed model association analysis for the 25 raw GWAS**

**Supplementary Figure 8: Average estimates of enrichment from the 24 summary GWAS**

**Supplementary Figure 9: Estimates of genetic correlation for the the 24 summary GWAS**

**Supplementary Figure 10: Comparing estimates from linear and logistic regression for the 13 WTCCC GWAS**

**Supplementary Figure 11: Sample quality control for eMerge data**

**Supplementary Table 1: Details of the 24 functional annotations**

**Supplementary Table 2: Details of the 25 raw GWAS, and estimates of $h^2_{\text{SNP}}$ and confounding bias**

**Supplementary Table 3: Average estimates of enrichment from the 25 raw GWAS**

**Supplementary Table 4: Estimates of genetic correlation for the 25 raw GWAS**

**Supplementary Table 5: Comparing heritability models for the 25 raw GWAS using REML**

**Supplementary Table 6: Comparing heritability models for the 25 raw GWAS using SumHer**

**Supplementary Table 7: Introducing population structure for the 13 WTCCC GWAS**

**Supplementary Table 8: Details of the 24 summary GWAS**

**Supplementary Table 9: Comparing heritability models for the 24 summary GWAS using SumHer**

**Supplementary Table 10: Number of significant loci for the 24 summary GWAS after correction for confounding bias**

**Supplementary Table 11: Average estimates of enrichment from the 24 summary GWAS**

**Supplementary Table 12: Average estimates of enrichment from the 11 summary GWAS least impacted by genomic control**

**Supplementary Table 13: Estimates of genetic correlation for the 24 summary GWAS**

**Supplementary Table 14: Comparing the predictive performance of Classical and Bayesian PRS**

**Supplementary Table 15: Numbers of large-effect loci for the 24 summary GWAS**

**Supplementary Note: Instructions for using SumHer.** Here we provide step-by-step scripts for using SumHer to estimate confounding bias, SNP heritability, enrichment of heritability and genetic correlations from GWAS results. These analyses require the software LDAK (available at http://ldak.org); we assume that the LDAK executable is saved in current folder, so that LDAK can be run by typing `./ldak5.linux` (or `./ldak5.mac` for the Mac version). They also use PLINK (available at http://cog-genomics.org/plink2) and awk (installed by default in unix); see http://zzz.bwh.harvard.edu/plink/tutorial.shtml and http://ldak.org/awk for tutorials on each. Note that a backslash (\) at the end of a line indicates the command continues on the next line. This code is also available at http://ldak.org/protocol.

**Acquire reference panel.** Suppose our reference panel is stored in binary PLINK format in the files `ref.bed`, `ref.bim` and `ref.fam`. Ideally, Column 3 of `ref.bim` contains genetic distances (otherwise, replace `--window-cm` with `--window-kb` in the scripts below). The individuals in the reference panel should be ancestrally similar to those used in the GWAS. As we analyzed summary statistics from European-centric GWAS, we used 8 850 unrelated Caucasian individuals from the Health and Retirement Study[1] whose genotype data are available upon application from dbGaP (accession code phs000428.v2.p2). An alternative is to instead use the 404 non-Finnish Europeans from the 1000 Genomes Project,[2] whose data can be obtained as followed.

```
#Download sample IDs and extract Europeans
wget ftp://ftp.1000genomes.ebi.ac.uk/vol1/ftp/release/20130502/integrated_call_samples_v3.20130502.ALL.panel
awk < integrated_call_samples_v3.20130502.ALL.panel '($3=="EUR" && $2!="FIN"){print $1, $1}' > eur.keep

#Download data for each autosome, and convert using PLINK, extracting European individuals and SNPs with MAF>0.01
for j in {1..22}; do
wget ftp://ftp.1000genomes.ebi.ac.uk/vol1/ftp/release/20130502/\
ALL.chr$j.phase3_shapeit2_mvncall_integrated_v5a.20130502.genotypes.vcf.gz
./plink --vcf ALL.chr$j.phase3_shapeit2_mvncall_integrated_v5a.20130502.genotypes.vcf.gz \
--make-bed --out chr$j --maf 0.01 --keep eur.keep
done

#Now join these together, excluding multi-allelic SNPs and those with duplicate positions
rm list.txt; for j in {1..22}; do echo chr$j >> list.txt; done
./ldak5.linux --make-bed ref --mbfile list.txt --exclude-odd YES --exclude-dups YES
```

**Format summary statistics.** For the analyses below, we use summary statistics for alzheimers[3] (stored in the file `IGAP_stage_1.txt`, which can be downloaded from http://web.pasteur-lille.fr/en/recherche/u744/igap/igap_download.php) and for years of education[4] (stored in `EduYears_Main.txt.gz`, available at http://ssgac.org). To use with SumHer, the summary statistic files should contain columns labelled `Predictor`, `A1`, `A2`, `Direction`, `Stat` and `n`. The names and alleles should be consistent with those in the reference panel; `Direction` indicates whether the effect of the A1 allele is positive or negative (for quantitative traits, can use the effect size, for binary, the log odds); `Stat` is the $\chi^2(1)$ test statistic (if only $p$-values are available, these can instead be provided in a column labelled `P`), and `n` is the number of individuals used when testing that SNP. The predictor names must be unique, so you should check for (and remove) duplicates. Having processed the summary statistic files, we also want to identify SNPs within the major histocompatibility complex (Chromosome 6: 25-34 Mb), as well as SNPs which individually explain >1% of phenotypic variation, and SNPs in LD with these. If info scores are provided, we recommend excluding SNPs with score <0.95.

```
#Tidy alzheimers summary statistics - the paper tells us there were 17008 cases and 37154 controls
#If possible, compute a chi-squared test statistic (otherwise, could use P)
#For linear regression, can use (beta/SD)^2, for logistic regression, (log.odds/SD)^2
awk < IGAP_stage_1.txt '(NR>1){snp=$3;a1=$4;a2=$5;dir=$6;stat=($6/$7)*($6/$7);n=17008+37154}(NR==1)\
{print "Predictor A1 A2 Direction Stat n"}(NR>1 && (a1=="A"||a1=="C"||a1=="G"||a1=="T") \
&& (a2=="A"||a2=="C"||a2=="G"||a2=="T")){print snp, a1, a2, dir, stat, n}' > alz.txt

#Check for duplicates using the unix functions sort and uniq
#If this command results in no output, it means all predictors are unique
awk < alz.txt '{print $1}' | sort | uniq -d | head
```

```
#If there are duplicates, we can remove then using
mv alz.txt alz2.txt; awk '!seen[$1]++' alz2.txt > alz.txt

#Tidy years education summary statistics - the paper tells us the total sample size was 328917
gunzip -c EduYears_Main.txt.gz | awk '(NR>1){snp=$1;a1=$4;a2=$5;dir=$7;stat=($7/$8)*($7/$8);n=328917}\
(NR==1){print "Predictor A1 A2 Direction Stat n"}(NR>1 && $10!="NA" && (a1=="A"||a1=="C"||a1=="G"||a1=="T") \
&& (a2=="A"||a2=="C"||a2=="G"||a2=="T")){print snp, a1, a2, dir, stat, n}' > years.txt

#Check for duplicates - here there are none
awk < years.txt '{print $1}' | sort | uniq -d | head

#Get list of MHC SNPs (from reference panel)
awk < ref.bim '($1==6 && $4>25000000 && $4<34000000){print $2}' > mhc.snps

#Identify large-effect SNPs (those explaining more than 1% of phenotypic variance)
#This command uses the fact that the variance explained by each SNP is stat/(stat+n)
awk < alz.txt '(NR>1 && $5>$6/99){print $1}' > alz.big
awk < years.txt '(NR>1 && $5>$6/99){print $1}' > years.big
#Note there are no large-effect SNPs for years education

#Find SNPs tagging the alzheimers large-effect loci (within 1cM and correlation squared >.1)
./ldak5.linux --remove-tags alz --bfile ref --top-preds alz.big --window-cm 1 --min-cor .1

#Create exclusion files, containing mhc and (for alzheimers) SNPs tagging large-effect SNPs
cat mhc.snps alz.out > alz.excl
cat mhc.snps > years.excl
```

**Estimate SNP heritability.** To estimate $h^2_{\text{SNP}}$ there are two steps: first we compute a (1-part) tagfile which contains $q_j + \sum_{l \in N_j} q_l r^2_{jl}$ for each SNP; then we perform the regression to estimate $h^2_{\text{SNP}}$. To compute an LDAK tagfile, we must first compute LDAK weights; this can take a few hours, but can be efficiently parallelized.

```
#Calculate LDAK SNP weights; here, we calculate weights separately for each chromosome then merge
#The on-screen instructions explain how to further parallelize (use --calc-weights not --calc-weights-all)
for j in {1..22}; do
./ldak5.linux --cut-weights alz_chr$j --bfile ref --extract alz.txt --chr $j
./ldak5.linux --calc-weights-all alz_chr$j --bfile ref --extract alz.txt --chr $j
done

#Merge weights across chromosomes
cat alz_chr{1..22}/weights.short > alz.weights

#Calculate the (1-part) LDAK tagfile
./ldak5.linux --calc-tagging alz_ldak --bfile ref --extract alz.txt --weights alz.weights --power -.25 --window-cm 1

#Perform the regression (remember to exclude the MHC / large-effect SNPs)
#Pay attention to the screen output (for this example it says I should add --check-sums NO)
./ldak5.linux --sum-hers alz_ldak --tagfile alz_ldak.tagging --summary alz.txt --exclude alz.excl

#Estimates of SNP heritability will be in alz_ldak.hers

#To instead use the GCTA Model, use --ignore-weights YES and --power -1 when computing the tagfile
#This takes longer (as it uses all SNPs), but we can calculate separately for each chromosome then merge
for j in {1..22}; do
./ldak5.linux --calc-tagging alz_gcta$j --bfile ref --extract alz.txt --ignore-weights YES \
--power -1 --window-cm 1 --chr $j
done
```

```
#Join the tagfiles across chromosomes
rm list.txt; for j in {1..22}; do echo "alz_gcta$j.tagging" >> list.txt; done
./ldak5.linux --join-tagging alz_gcta --taglist list.txt

#Then perform the regression (again will have to add --check-sums NO)
./ldak5.linux --sum-hers alz_gcta --tagfile alz_gcta.tagging --summary alz.txt --exclude alz.excl
```

**Estimate confounding bias.** We recommend estimating the scaling factor $C$, but SumHer can also estimate the LDSC intercept $1+A$. There is no need to compute a new tagfile, just add `--genomic-control YES` or `--intercept YES` when performing the regression.

```
#Estimate the scaling factor C
./ldak5.linux --sum-hers alz_ldak.gcon --tagfile alz_ldak.tagging --summary alz.txt --exclude alz.excl \
--genomic-control YES
#The scaling factor will be in alz_ldak.gcon.extra

#Estimate the intercept 1+A
./ldak5.linux --sum-hers alz_gcta.cept --tagfile alz_gcta.cept.tagging --summary alz.txt --exclude alz.excl \
--intercept YES
#The intercept will be in alz_gcta.cept.extra
```

**Estimate functional enrichments.** Now we need to create a (multi-part) tagfile, which contains $q_j I_{jk} + \sum_{l \in N_j} q_j I_{lk} r^2_{jl}$ for each SNP and category. To work out which SNPs are in each of the 24 annotations, we use the genefiles in `annotations.zip`, available at http://ldak.org/annotations (these are modified versions of the bedfiles provided at https://data.broadinstitute.org/alkesgroup/LDSCORE).

```
#First work out which (reference panel) SNPs are in each annotation
#Will save these in ann_snps.1, ..., ann_snps.24 (the SNP lists must have consecutive suffixes)
for j in {1..24}; do
./ldak5.linux --cut-genes ann$j --bfile ref --genefile ann$j.genefile --ignore-weights YES
mv ann$j/genes.predictors.used ann_snps.$j
done

#Now compute a tagfile - as the categories overlap, we use --annotation-number and --annotation-prefix
./ldak5.linux --calc-tagging alz_ldak.ann --bfile ref --extract alz.txt --weights alz.weights \
--power -.25 --window-cm 1 --annotation-number 24 --annotation-prefix ann_snps.

#Perform the regression (adding --exclude, --check-sums NO, --genomic-control YES as required)
./ldak5.linux --sum-hers alz_ldak.ann --tagfile alz_ldak.ann.tagging --summary alz.txt --exclude alz.excl

#The category enrichments and SDs are in Columns 5 and 6 of alz_ldak.ann.enrich
#Suppose we have results for alz, years and height; to compute inverse-variance weighted averages enrichments use
for i in {1..24}; do
grep "Enrich_A$i " {alz,years,height}_ldak.ann.enrich | awk -v i=$i '{a+=$5/$6/$6;b+=1/$6/$6}END\
{print "Annotation:", i, "Av. Enr.:", a/b, "SD:", 1/sqrt(b), "Num. Traits:", NR}'
done

#For the PRS, we must save the average heritability fractions of each annotation and the base (use the .share files)
rm props.ldak
for i in {1..24}; do
grep "Share_A$i " {alz,years,height}_ldak.ann.share | awk -v i=$i '{a+=$2/$3/$3;b+=1/$3/$3}END\
{print a/b, 1/sqrt(b)}' >> props.ldak
done
grep Share_Base {alz,years,height}_ldak.ann.share | awk -v i=$i '{a+=$2/$3/$3;b+=1/$3/$3}END\
{print a/b, 1//sqrt(b)}' >> props.ldak
```

```
#To instead use the GCTA Model, use --ignore-weights YES and --power -1 when computing the tagfile
#Note that to copy Finucane et al, you must use all 52 annotations (not just the first 24)
./ldak5.linux --calc-tagging alz_ldak.ann --bfile ref --extract alz.txt --ignore-weights YES --power -1 --window-cm
    1 --annotation-number 24 --annotation-prefix ann_snps.

#If categories partition the genome (LDAK allows some overlap), should use --partition-number and --partition-prefix
#In this example, we construct lists part.1 and part.2 containing coding (Annotation 1) and non-coding SNPs
cp ann_snps.1 part.1
awk '(NR==FNR){arr[$1];next}!($2 in arr){print $2}' part.1 ref.bim > part.2
./ldak5.linux --calc-tagging alz_ldak.coding --bfile ref --extract alz.txt --weights alz.weights \
--power -.25 --window-cm 1 --partition-number 2 --partition-prefix part.

#Note that we can achieve the same using
./ldak5.linux --calc-tagging alz_ldak.coding --bfile ref --extract alz.txt --weights alz.weights \
--power -.25 --window-cm 1 --partition-number 1 --partition-prefix part. --background YES
```

**Estimate genetic correlations.** We calculate a (1-part) tagfile, using only SNPs common to the two traits (here alzheimers and years education). Now the direction of effects matters, so we recommend using only non-ambiguous SNPs (alleles A & C, A & G or C & T), unless very confident the alleles are consistently aligned (note that if you include ambiguous SNPs, you must add `--allow-ambiguous YES` when performing the regression).

```
#Find the overlap of non-ambiguous SNPs
awk < alz.txt '(($2=="A"&&$3=="C") || ($2=="A"&&$3=="G") || ($2=="C"&&$3=="T") || ($2=="C"&&$3=="A") \
||($2=="G"&&$3=="A") || ($2=="T"&&$3=="C")){print $1}' > alz.nonamb
awk < years.txt '(($2=="A"&&$3=="C") || ($2=="A"&&$3=="G") || ($2=="C"&&$3=="T") || ($2=="C"&&$3=="A") \
||($2=="G"&&$3=="A") || ($2=="T"&&$3=="C")){print $1}' > years.nonamb
awk '(NR==FNR){arr[$1];next}($1 in arr){print $1}' alz.nonamb years.nonamb > overlap.nonamb

#Will exclude SNPs in the exclusion lists for either trait
cat alz.excl years.excl | sort | uniq > alz_years.excl

#Calculate LDAK weights for the overlap
for j in {1..22}; do
./ldak5.linux --cut-weights alz_years_chr$j --bfile ref --extract overlap.nonamb --chr $j
./ldak5.linux --calc-weights-all alz_years_chr$j --bfile ref --extract overlap.nonamb --chr $j
done
cat alz_years_chr{1..22}/weights.short > alz_years.weights

#Calculate LDAK tagging file
./ldak5.linux --calc-tagging alz_years_ldak --bfile ref --extract overlap.nonamb --weights alz_years.weights \
--power -.25 --window-cm 1

#Perform the regression (adding --exclude, --check-sums NO, --genomic-control YES as required)
./ldak5.linux --sum-cors alz_years_ldak --tagfile alz_years_ldak.tagging --summary alz.txt --summary2 years.txt \
--exclude alz_years.excl

#The estimate of genetic correlation is in alz_years_ldak.cors
```

**Comparing heritability models.** The easiest way to compare two heritability models is by likelihood. By default, LDAK excludes SNPs with zero weight when computing tagfiles. However, when comparing heritability models, it is important that each tagfile contain the same SNPs. Therefore, when computing an LDAK tagfile, add `--reduce NO`. The second way to compare two heritability models is by creating a merged tagfile, then LDAK will estimate the share of heritability allocated to each.

```
#Comparing the GCTA and LDAK Models by likelihood

#Compute (standard) GCTA tagfile and the (unreduced) LDAK tagfile (as before, could divide by chromosome)
./ldak5.linux --calc-tagging alz_gcta --bfile ref --extract alz.txt --ignore-weights YES --power -1 --window-cm 1
./ldak5.linux --calc-tagging alz_ldak.non$j --bfile ref --extract alz.txt --weights alz.weights \
--power -.25 --window-cm 1 --reduce NO

#Perform the regression for each (adding --exclude, --check-sums NO, --genomic-control YES as required)
./ldak5.linux --sum-hers alz_gcta --tagfile alz_gcta.tagging --summary alz.txt --exclude alz.excl
./ldak5.linux --sum-hers alz_ldak.non --tagfile alz_ldak.non.tagging --summary alz.txt --exclude alz.excl

#The likelihoods and likelihood ratio test statistics are in alz_gcta.extra and alz_ldak.non.extra

#Comparing the GCTA and LDAK Models by creating a merged tagfile
echo "alz_gcta.tagging
alz_ldak.non.tagging" > list.txt
./ldak5.linux --merge-tagging alz_both --taglist list.txt
./ldak5.linux --sum-hers alz_both --tagfile alz_both.tagging --summary alz.txt --exclude alz.excl

#The share of heritability allocated to the LDAK Model (p) is the value Share_P2 in alz_both.share
```

**Polygenic risk scores.** For each of body mass index, height, HDL & LDL cholesterol and triglyceride, we computed five PRS (one classical and four Bayesian) using the results from the 24 Summary GWAS, then measured how well each of these predicted for the eMERGE data. To avoid strand issue, we reduced to non-ambiguous SNPs. To calculate a Bayesian PRS, we first compute a tagging file (either 1-part or multi-part), then the prior distribution for the heritability tagged by each SNP, and finally the posterior mean effect sizes. We found that the Bayesian PRS benefited from clumping, but not the Classical PRS. Here we demonstrate for height, using results from the most recent GIANT Consortium meta-analysis.[5] Suppose the eMERGE data are stored in `emerge.bed`, `emerge.bim` and `emerge.fam`, and the corresponding phenotypes are in `height.pheno`. We will save results in a folder called `prs`.

```
#Identify the non-ambiguous eMERGE SNPs
awk < emerge.bim '(($5=="A"&&$6=="C") || ($5=="A"&&$6=="G") || ($5=="C"&&$6=="T") || \
($5=="C"&&$6=="A") ||($5=="G"&&$6=="A") || ($5=="T"&&$6=="C")){print $2}' > emerge.nonamb

#Format the summary statistics for height - now per-SNP sample sizes are provided
wget https://portals.broadinstitute.org/collaboration/giant/images/0/01/\
GIANT_HEIGHT_Wood_et_al_2014_publicrelease_HapMapCeuFreq.txt.gz
gunzip -c GIANT_HEIGHT_Wood_et_al_2014_publicrelease_HapMapCeuFreq.txt.gz | awk '(NR>1){snp=$1;a1=$2;a2=$3;\
dir=$5;p=$7;n=$8}(NR==1){print "Predictor A1 A2 Direction P n"}(NR>1 && (a1=="A"||a1=="C"||a1=="G"||a1=="T") \
&& (a2=="A"||a2=="C"||a2=="G"||a2=="T")){print snp, a1, a2, dir, p, n}' - > height.txt

#Check for duplicates (there are none)
awk < height.txt '{print $1}' | sort | uniq -d | head

#Each PRS scorefile should have the columns Predictor, A1, A2, Centre (can set to NA) and Effect

#For the classical PRS, the effect size of each SNP is its correlation, equal to ssign(direction)*(stat/(stat+n)^.5
awk < height.txt '(NR==1){print "Predictor A1 A2 Centre Effect"}(NR>1){r=sqrt($5/($5+$6));\
if($4<0){r=-r};print $1, $2, $3, "NA", r}' >> prs/height_class.score

#To calculate the two LDAK Bayesian PRS, we must first calculate LDAK weights
for j in {1..22}; do
./ldak5.linux --cut-weights height_chr$j --bfile ref --extract height.txt --chr $j
./ldak5.linux --calc-weights-all height_chr$j --bfile ref --extract height.txt --chr $j
done
cat height_chr{1..22}/weights.short > height.weights
```

```
#Now compute 1-part and 25-part tagfiles; use --reduce NO to ensure they contain all SNPs
./ldak5.linux --calc-tagging height_ldak.non --bfile ref --extract height.txt --weights height.weights \
--power -.25 --window-cm 1 --reduce NO
./ldak5.linux --calc-tagging height_ldak.ann.non --bfile ref --extract height.txt --weights height.weights \
--power -.25 --window-cm 1 --annotation-number 24 --annotation-prefix ann_snps. --reduce NO

#Compute the expected heritability tagged by each SNP
#For this we must provide an estimate of SNP heritability (we recommend using the estimates from --sum-her)
#For the enriched LDAK PRS, we must also provide the heritability fractions computed above
her=`grep Her_ALL height_ldak.non.hers | awk '{print $2}'`
./ldak5.linux --calc-exps prs/height_ldak.non --tagfile height_ldak.non.tagging --her $her
her=`grep Her_ALL height_ldak.ann.non.hers | awk '{print $2}'`
./ldak5.linux --calc-exps prs/height_ldak.ann.non --tagfile height_ldak.ann.non.tagging --her $her \
--props props.ldak

#Next we compute the posterior means (again watch screen output; will likely have to add --check-sums NO)
./ldak5.linux --calc-posts prs/height_ldak.non --expectations prs/height_ldak.non.exps --summary height.txt
./ldak5.linux --calc-posts prs/height_ldak.ann.non --expectations prs/height_ldak.ann.non.exps --summary height.txt

#Finally, we clump; we add --extract emerge.nonamb, as we will only be using these when predicting
#Normally to clump we provide pvalues; here we instead provide .psuedos files (which rank SNPs by Bayes factors)
./ldak5.linux --thin prs/height_ldak.non --bfile ref --extract emerge.nonamb --window-cm 1 --window-prune .1 \
--pvalues prs/height_ldak.non.psuedos
./ldak5.linux --thin prs/height_ldak.ann.non --bfile ref --extract emerge.nonamb --window-cm 1 --window-prune .1 \
--pvalues prs/height_ldak.ann.non.psuedos

#To test each PRS, we project onto the eMERGE data; at this step, it is convenient to provide the phenotypes
#All PRS contain standardized effect sizes, so we use --power -1 (for raw effect sizes, use --power 0)
#Having excluded ambiguous, it should be OK to allow flips; likely a few SNPs will have inconsistent alleles
./ldak5.linux --calc-scores prs/height_class --bfile emerge --extract emerge.nonamb \
--scorefile prs/height_class.score --pheno height.pheno --power -1 --allow-flips YES --allow-inconsistent YES
./ldak5.linux --calc-scores prs/height_ldak.non --bfile emerge --extract prs/height_ldak.non.in \
--scorefile prs/height_ldak.non.score --pheno height.pheno --power -1 --allow-flips YES --allow-inconsistent YES
./ldak5.linux --calc-scores prs/height_ldak.ann.non --bfile emerge --extract prs/height_ldak.ann.non.in \
--scorefile prs/height_ldak.ann.non.score --pheno height.pheno --power -1 --allow-flips YES --allow-inconsistent YES

#Predictions are stored in prs/height_class.profile, prs/height_ldak.non.profile and prs/height_ldak.ann.non.profile
#We measured performance by calculating rho, the correlation between predicted and observed phenotypes
#We estimated the SD of rho by block-jackknifing 500 times for each PRS; for each jackknife we computed rho_z
#the correlation across 99.8% of individuals, then the estimate of the SD of rho is SD(rho_z)*499^.5
```

**Additional scripts.** Our previous paper[6] provided scripts for estimating inflation due to confounding via REML and for performing association analysis, including how to construct covariates (when performing linear regression, we included sex and ten principal components; five derived from the reference panel, five from the 1000 Genomes Project[2]). To simulate phenotypes under the GCTA and LDAK Models, we used the following scripts (these assume the Wellcome Trust control individuals[7] are stored in `wt.bed`, `wt.bim` and `wt.fam`).

```
#Simulating 500 phenotypes (each with SNP heritability 0.5 and 2000 causal SNPs)
./ldak5.linux --make-phenos gcta --bfile wt --ignore-weights YES --power -1 --num-causals 2000 --num-phenos 500
./ldak5.linux --make-phenos ldak --bfile wt --weights ref.weights --power -.25 --num-causals 2000 --num-phenos 500

#Simulating 250 pairs of phenotypes (consecutive pairs have genetic correlation 0.5)
./ldak5.linux --make-phenos gcta_bivar --bfile wt --ignore-weights YES --power -1 --num-causals 2000 \
--num-phenos 500 --bivar .5
./ldak5.linux --make-phenos ldak_bivar --bfile wt --weights ref.weights --power -.25 --num-causals 2000 \
--num-phenos 500 --bivar .5
```

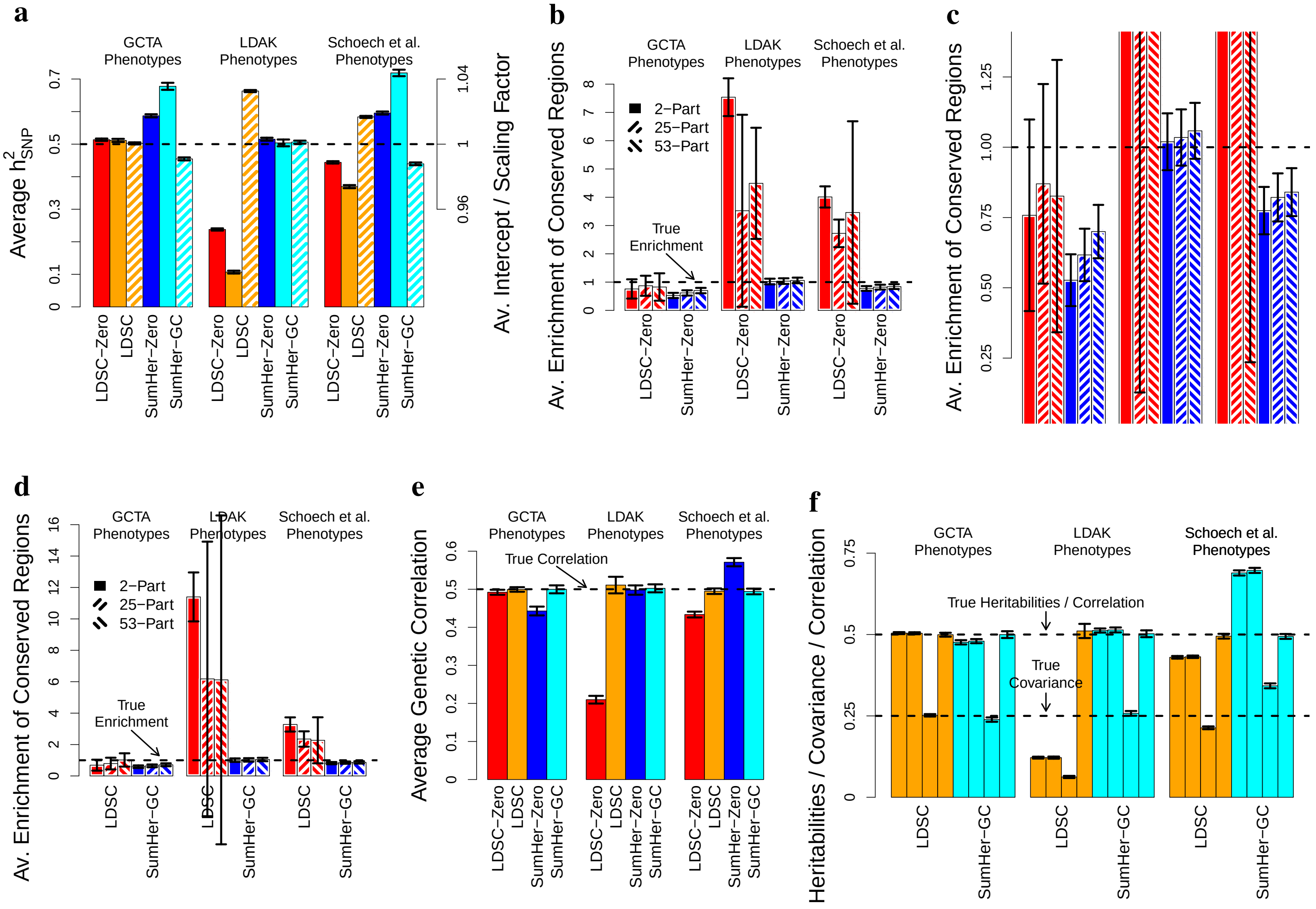


**Supplementary Figure 1: Simulations demonstrating the importance of the assumed heritability model.** We use genotypes from the three population cohorts of the Wellcome Trust Case Control Consortium:[7] 1958 Birth Cohort, National Blood Service and People of the British Isles (after quality control, 7 548 individuals, 3 280 768 SNPs with MAF>0.01). First we generate 1500 phenotypes, each with 2000 causal SNPs (picked uniformly at random) and $h^2_{\text{SNP}} = 0.5$, using the model $Y = \sum_j \beta_j X_j + e$, where $\beta_j$ is the effect size of $X_j$, the $j$th causal SNP, and $e$ is Gaussian-distributed noise. For 500 phenotypes, we sample effect sizes according to the GCTA Model ($\beta_j \sim \mathbb{N}(0, [f_j(1-f_j)]^{-1})$, where $f_j$ is the MAF of $X_j$), the next 500 phenotypes are sampled according to the LDAK Model ($\beta_j \sim \mathbb{N}(0, [f_j(1-f_j)]^{-0.25} w_j)$, where $w_j$ is the LDAK weight for $X_j$), while the final 500 phenotypes are sampled according to a model proposed by Schoech *et al.* ($\beta_j \sim \mathbb{N}(0, [f_j(1-f_j)]^{-0.38}(1-0.3LLD_j))$, where $LLD_j = \sum_{l \in N_j} r^2_{jl}$ is the LD Score[8] of $X_j$). **(a)** Average estimates of $h^2_{\text{SNP}}$ from each of LDSC-Zero, LDSC, SumHer-Zero and SumHer-GC. **(b)** Average estimates of enrichment of SNPs in conserved regions (one of the 24 functional categories) from LDSC-Zero and SumHer-Zero. For each method, we estimate enrichment in three ways: using a 2-part model (conserved and non-conserved SNPs), a 25-part model (one for each of the 24 functional categories, plus one for all other SNPs), and a 53-part model (24 categories, 28 buffer regions and all other SNPs, the approach preferred by Finucane *et al.*[9]). **(c)** Zoomed-in version of (b). **(d)** Same as (b), except now we estimate enrichment using LDSC and SumHer-GC

Next we generate 1500 pairs of phenotypes each with genetic correlation 0.5. Again, each phenotype has 2000 causal SNPs and $h^2_{\text{SNP}} = 0.5$, except now, in order to ensure reasonably precise estimates, we restrict to Chromosomes 1 & 2 (547 948 SNPs). **(e)** Estimates of genetic correlation from LDSC-Zero, LDSC, SumHer-Zero and SumHer-GC. We see that, by allowing for confounding bias (i.e., using LDSC or SumHer-GC), it is possible to obtain accurate estimates of genetic correlation under all three heritability models. **(f)** Average estimates of $h^2_{\text{SNP A}}$ & $h^2_{\text{SNP B}}$ (the SNP heritability of the first and second phenotype in each pair), $h^2_{\text{COV}}$ (their genetic covariance) and $h^2_{\text{COV}}/\sqrt{h^2_{\text{SNP A}} h^2_{\text{SNP B}}}$ (their genetic correlation), from LDSC and SumHer-GC. We see that even though assuming a poor heritability model causes over/under-estimation of the SNP heritabilities, it also causes over/under-estimation of the genetic covariance by on average the same amount, resulting in accurate estimates of genetic correlation.

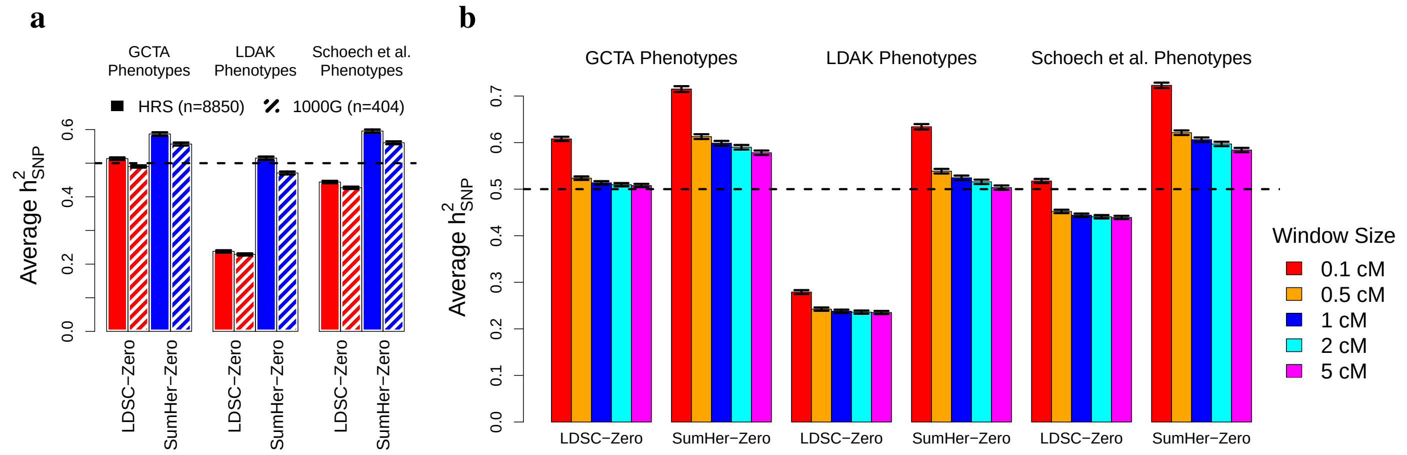


**Supplementary Figure 2: Choice of reference panel and varying the LD window size.** For these analyses, we use the 1500 simulated phenotypes used for Main Text Figures 1a & 1b, and described in Supplementary Figure 1. **(a)** Our preferred reference panel is 8 850 unrelated Caucasian individuals from the Health and Retirement Study (HRS);[1] these data are available upon application from dbGaP (accession code phs000428.v2.p2). An alternative is to use the 404 non-Finnish European individuals from the 1000 Genome Project[2] (see the Supplementary Note for instructions on how to download and prepare these for use with SumHer). Bars report average estimates of $h^2_{\text{SNP}}$ from LDSC-Zero and SumHer-Zero, using either the HRS (solid bars) or 1000 Genome Project (hatched bars) as the reference panel (true $h^2_{\text{SNP}} = 0.5$). While the results suggest it is better to use the HRS, which is to be expected because its larger sample size enables more accurate estimation of $r^2_{jl}$, we see the impact of instead using the 1000 Genome Project individuals is slight, indicating that these also suffice as a reference panel. **(b)** Equation (3) in Online Methods uses the approximation $v_j^2 = h_j^2 + \sum_{l \in N_j} r^2_{jl}\, h^2_l$, where $N_j$ indexes SNPs "near" SNP $j$. This is based on the assumption that $r^2_{jl}$ will be negligible for SNPs sufficiently far apart. Bars report average estimates of $h^2_{\text{SNP}}$ from LDSC-Zero and SumHer-Zero, for each of five definitions of $N_j$ (SNPs within 0.1, 0.5, 1, 2 or 5 cM, indicated by bar color). The window size should be sufficiently large to capture the majority of tagging due to LD, but not too large to make computation prohibitively slow. The fact that estimates change little when we increase the cutoff to 5 cM, indicates that our recommended choice, 1 cM performs well.

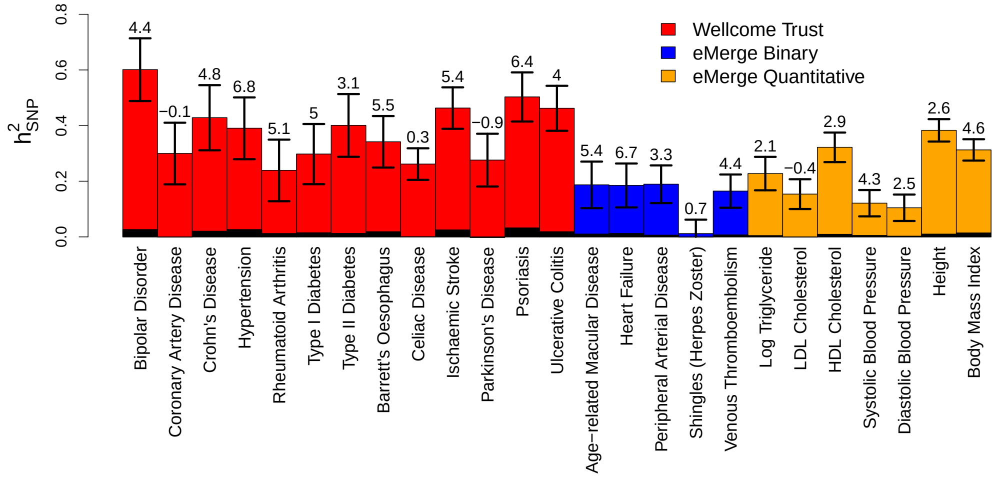


**Supplementary Figure 3: REML estimates of confounding for the 25 raw GWAS.** For each trait, we compute $K_{\text{LEFT}}$, a kinship matrix from Chromosomes 1-8, and $K_{\text{RIGHT}}$, a kinship matrix from Chromosomes 9-22, each time assuming the LDAK Model.[10] Then, using REML, we estimate $h^2_{\text{LEFT}}$ (by regressing the phenotype on $K_{\text{LEFT}}$), $h^2_{\text{RIGHT}}$ (by regressing the phenotype on $K_{\text{RIGHT}}$), and $h^2_{\text{ALL}}$ (by regressing the phenotype on $K_{\text{LEFT}}$ and $K_{\text{RIGHT}}$). Our estimate of inflation due to population structure and cryptic relatedness is then $(h^2_{\text{LEFT}} + h^2_{\text{RIGHT}})/h^2_{\text{ALL}}$. Bars report estimates of $h^2_{\text{ALL}}$ for each trait; the vertical line segments mark 95% confidence intervals. The black rectangle at the base of each bar indicates the estimated inflation, while the number above the bar expresses this as a percentage of $h^2_{\text{ALL}}$.

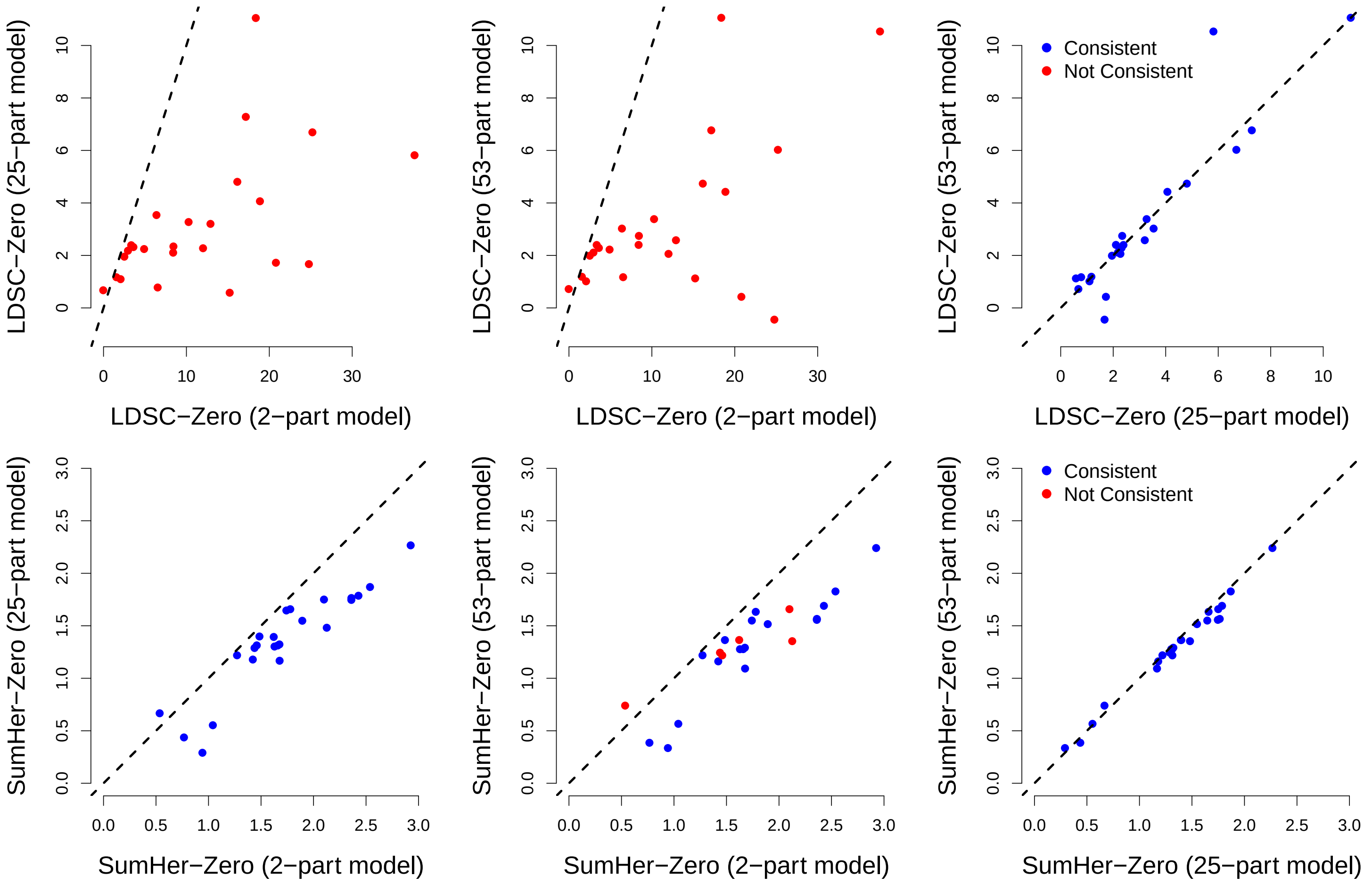


**Supplementary Figure 4: Comparing 2-part, 25-part and 53-part models for estimating enrichment for the 25 raw GWAS.** The figures compare, for LDSC-Zero (top row) and SumHer-Zero (bottom row), average estimates of enrichment for each of the 24 functional categories considered by Finucane *et al.*,[9] obtained using a 2-part, 25-part or 53-part model. For the 2-part model, for each category in turn, we partition SNPs into those inside and those outside the category. For the 25-part model, we divide the genome into a set for each of the 24 categories, plus a set containing all other SNPs. For the 53-part model, we divide the genome into a set for each of the 24 categories, a set for each of 28 buffer regions (see Finucane *et al.*,[9]), plus a set containing all other SNPs. In all panels, red dots indicate a significant discrepancy ($P < 0.05$) between a pair of estimates. It is problematic if estimates of enrichment vary according to how the genome is partitioned, as then it is difficult to know on which to rely. We see that for LDSC-Zero, none of the 2-part estimates are consistent with those from the 25-part and 53-part models (including those which appear very close to the diagonal), whereas for SumHer-Zero, the majority are consistent.

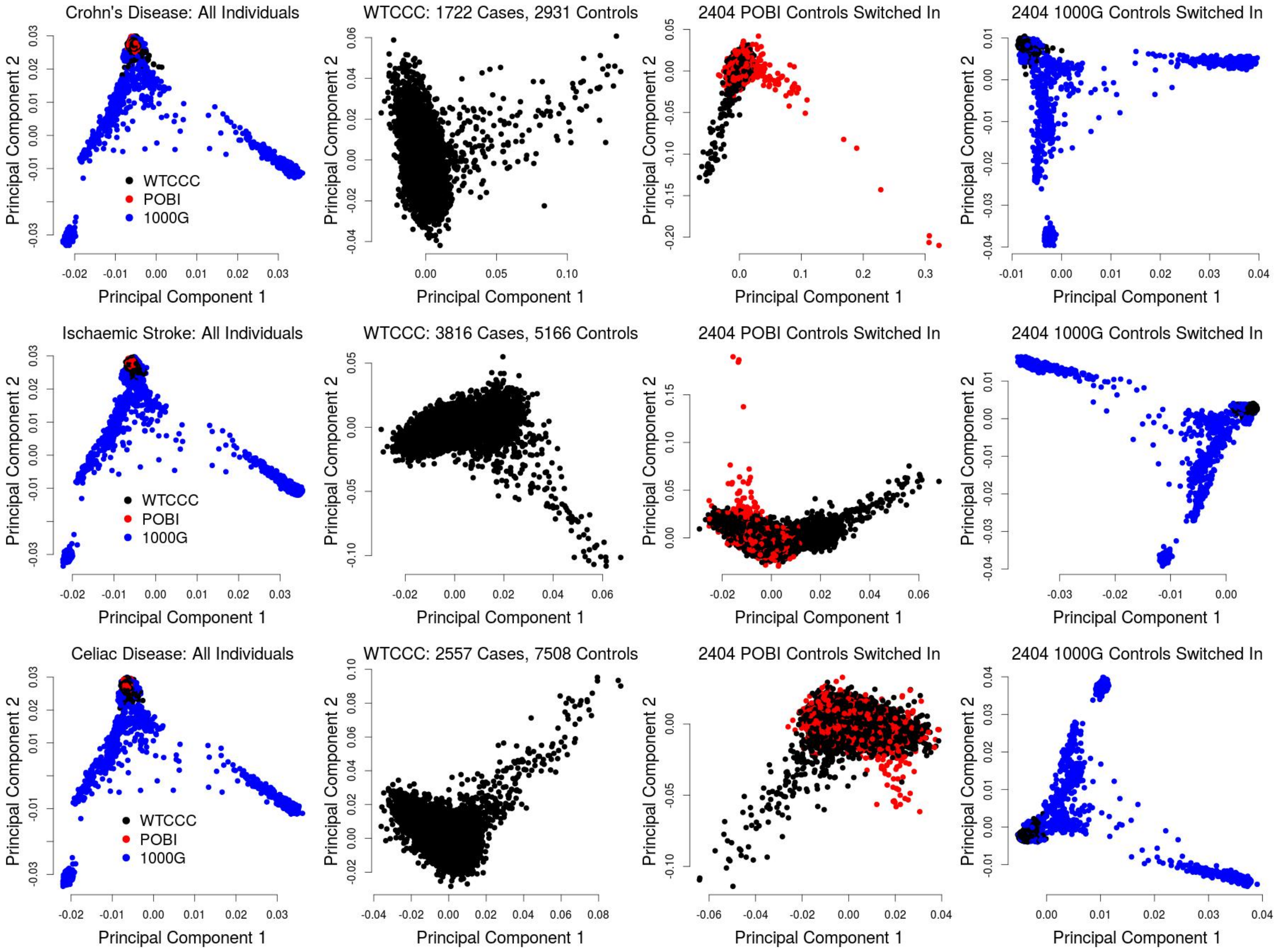


**Supplementary Figure 5: Introducing population structure for the 13 WTCCC GWAS.** For each GWAS, we replace 2 404 randomly picked controls with either 2 404 individuals from People of the British Isles[11] (POBI) or the 2 404 individuals from the 1000 Genome Project[2] (1000G). These principal component plots demonstrate the impact of switching controls for Crohn's Disease (top row), Ischaemic Stroke (middle row) and Celiac Disease (bottom row), the smallest and two largest GWAS, respectively. Points are colored according to collection (black: WTCCC; red: POBI; blue: 1000G). Although both the WTCCC controls and POBI individuals were recruited from the UK, switching in the latter generates modest structure (Column 3), because the POBI individuals came from predominantly isolated, rural regions. Switching in 1000G individuals generates extreme structure (Column 4), because these include Admixed Americans, Africans and Asians, whereas the WTCCC individuals are Caucasian. In practice, we would expect population structure to be modest, rather than extreme, as when performing a GWAS it is standard practice to identify and exclude ancestral outliers.[12]

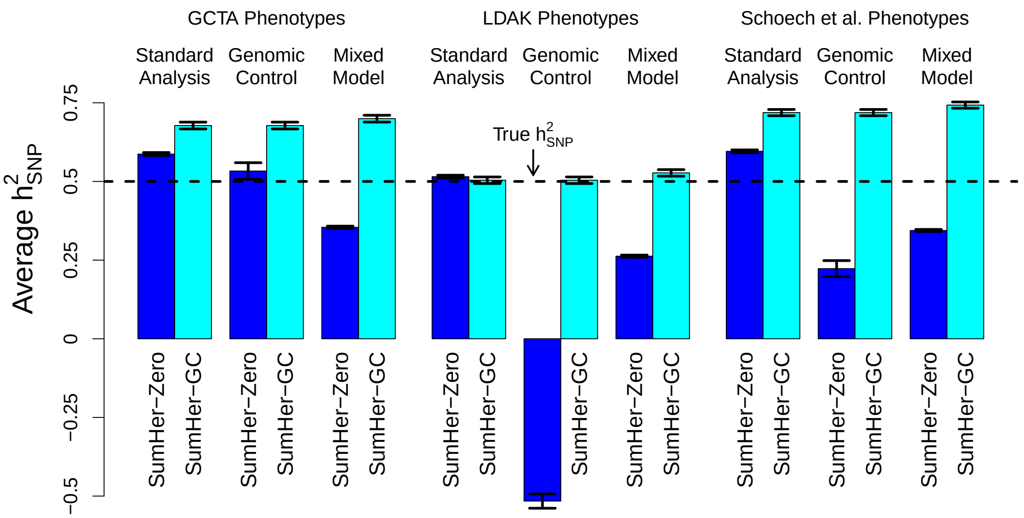


**Supplementary Figure 6: Simulations investigating the impact of genomic control and mixed model association analysis.** For these analyses, we use the 1500 simulated phenotypes used for Main Text Figures 1a & 1b, and described in Supplementary Figure 1. For each phenotype, we perform three types of association analysis. **Standard analysis** tests each SNP using classical (least squares) linear regression. **Genomic control** also performs classical linear regression, but then divides test statistics by the genomic inflation factor. **Mixed Model** tests each SNP using a linear mixed model[13] (to generate the kinship matrix, we thin SNPs then compute allelic correlations[14]). Bars report estimates of $h^2_{\text{SNP}}$ from SumHer-Zero and SumHer-GC; vertical line segments mark 95% confidence intervals. We recommend using SumHer-GC, instead of SumHer-Zero, if test statistics have been subjected to genomic control or were obtained from mixed model association analysis. The results for LDAK Phenotypes demonstrate why. They show that whereas SumHer-Zero will tend to under-estimate $h^2_{\text{SNP}}$ when either genomic control or mixed model association analysis has been used, SumHer-GC will produce accurate estimates. However, the results for GCTA and Schoech *et al.* Phenotypes emphasize that the effectiveness of SumHer-GC depends on the appropriateness of the LDAK Model. Figure 4b in the main text shows that in general, estimates from SumHer-GC post genomic control were consistent with those from SumHer-Zero pre genomic control, reflecting that in general the LDAK Model performs well. However, we recognize that for some traits, the LDAK Model will be sub-optimal, and this likely explains why SumHer-GC performed poorly for Ischaemic Stroke.

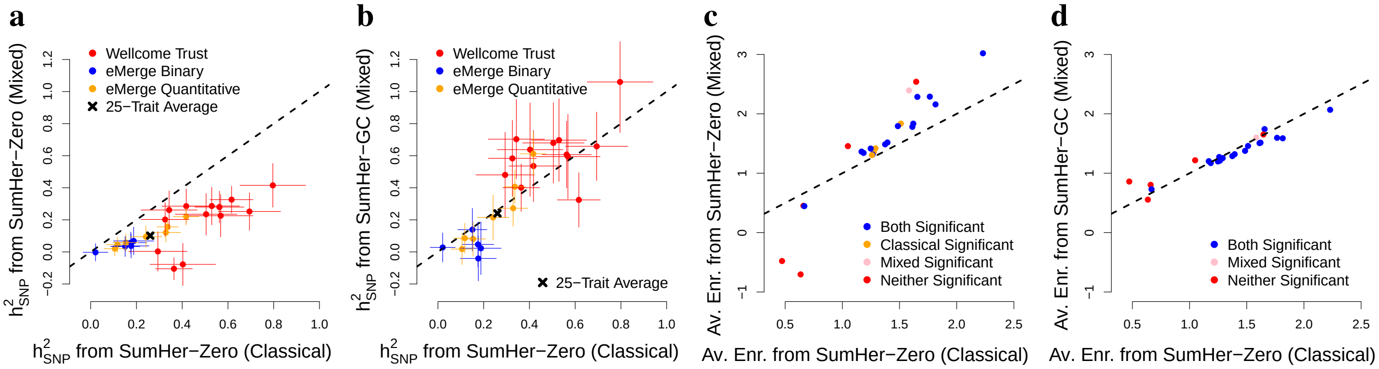


**Supplementary Figure 7: Mixed model association analysis for the 25 raw GWAS.** Each plot compares estimates from SumHer using test statistics from classical linear regression ($x$-axis) with those using test statistics from mixed model association analysis ($y$-axis). Horizontal and vertical line segments mark 95% confidence intervals. **(a)** Estimates of $h^2_{\text{SNP}}$ using SumHer-Zero for both classical and mixed model test statistics. **(b)** Estimates of $h^2_{\text{SNP}}$ using SumHer-Zero for classical test statistic and SumHer-GC for mixed model test statistics. **(c)** Average estimates of enrichment for the 24 functional categories using SumHer-Zero for both classical and mixed model test statistics. **(d)** Average estimates of enrichment for the 24 functional categories using SumHer-Zero for classical test statistics and SumHer-GC for mixed model test statistics.

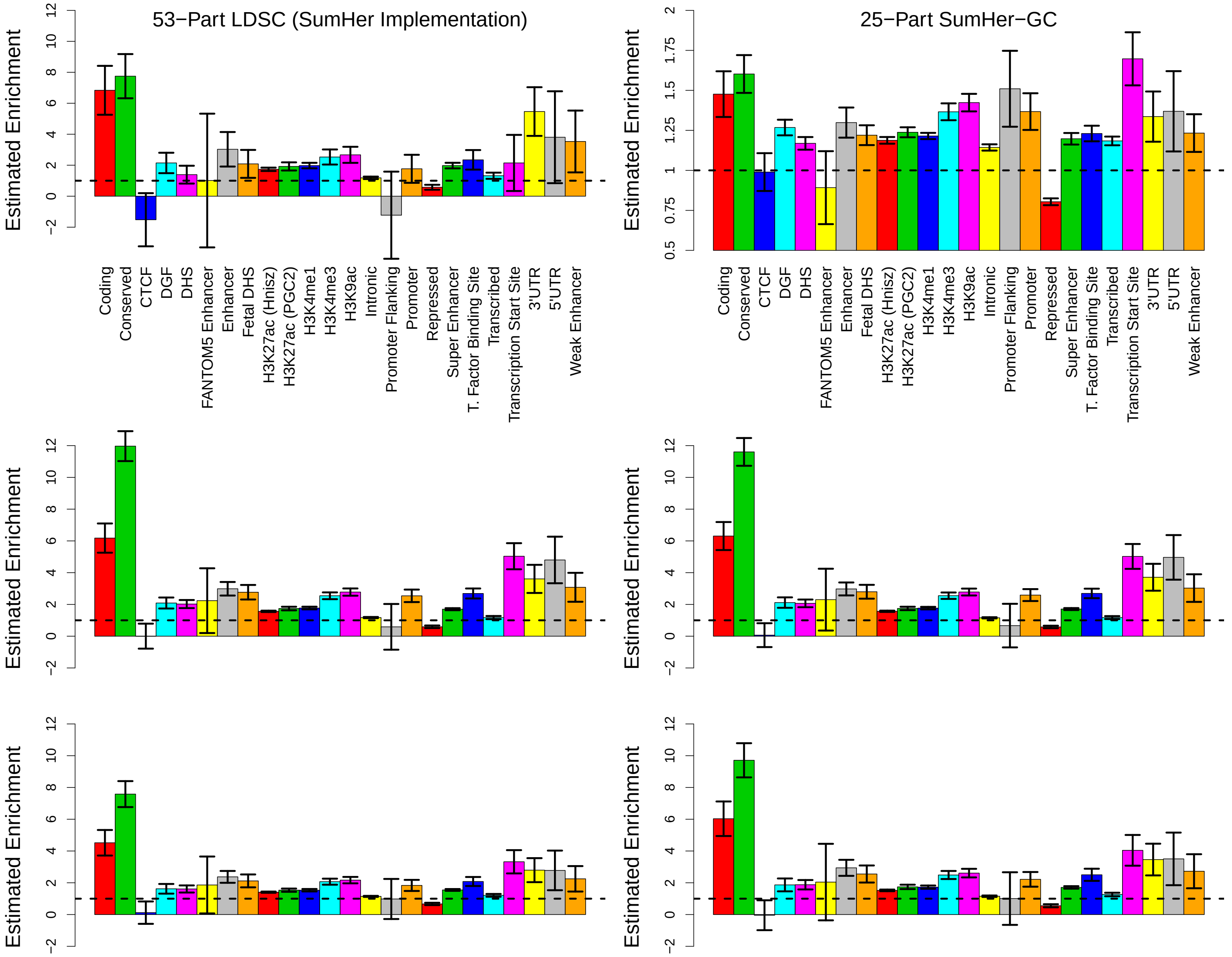


**Supplementary Figure 8: Average estimates of enrichment from the 24 summary GWAS.** Each barplot reports average estimates of enrichments for each of the 24 annotations; vertical line segments mark 95% confidence intervals. The top row presents results from (the SumHer implementation of) LDSC, using a 53-part model, and from SumHer-GC, using a 25-part model. These are the same results presented in Figure 5a in the Main Text. When running SumHer, we ensure that the SNPs in the reference panel match those used for the regression (we achieve this by restricting to SNPs which are both present in the Health and Retirement Study[1] and for which we have summary statistics). By contrast, the authors of LDSC[8] recommend that the reference panel contains as many SNPs as possible, but that the regression should use only HapMap 3[15] SNPs with MAF >0.05. To examine the impact this has, we now estimate enrichments using the LDSC software,[8] following the recommendations on https://github.com/bulik/ldsc (in particular, we use the reference panel they supply, which comprises 489 European individuals from the 1000 Genomes Project,[2] and also use the precomputed LD scores). For the middle row, we use the approach of Finucane *et al.*,[9] (who used the 53-part model); for the bottom row we use the approach of Gazal *et al.*, who instead used a 75-part model (to the 53 categories, they add 3 more functional annotations, 3 extra buffers, 10 MAF tranches and 6 continuous LD-related annotations). By adding the option `--not-M-5-50`, LDSC will perform the regression using HapMap 3 SNPs with MAF>0.01, rather than just those with MAF >0.05. Regardless of settings, we see that estimates from the LDSC software are very similar to those from the SumHer implementation of LDSC, and very different to those from SumHer-GC.

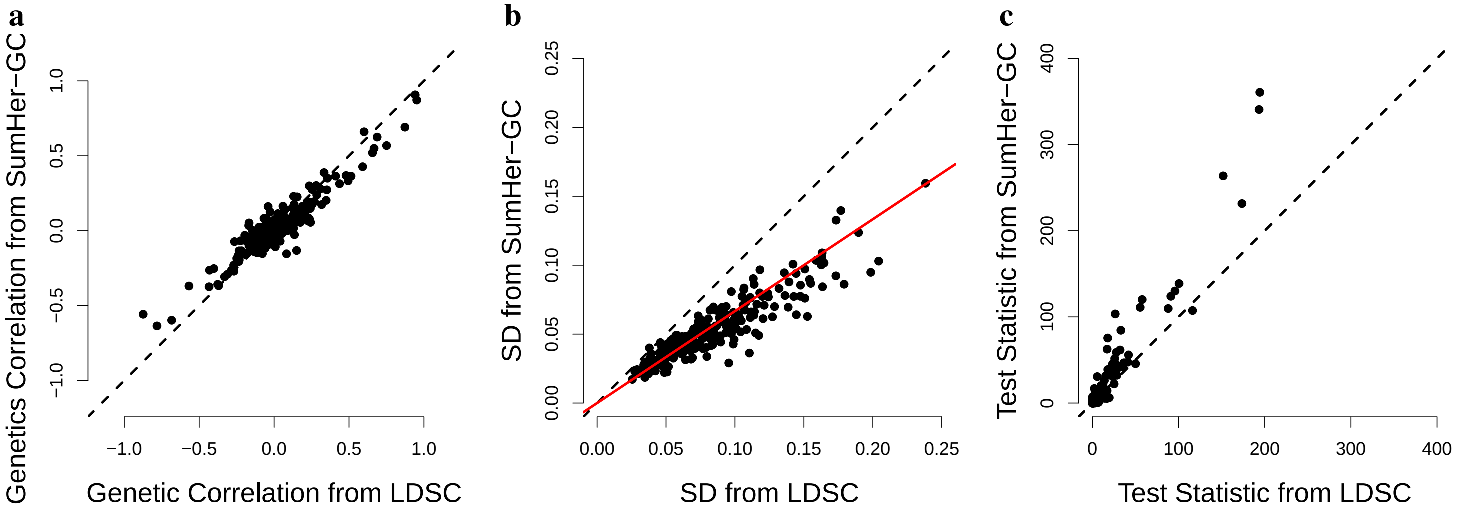


**Supplementary Figure 9: Estimates of genetic correlation for the 24 summary GWAS.** For each of the 276 pairs of traits, we estimate genetic correlation using LDSC ($x$-axis) and ($y$-axis). **(a)** Estimates of genetic correlation. **(b)** Standard deviations of these estimates; the red line marks $y = 2x/3$. **(c)** $\chi^2(1)$ test statistics for significant genetic correlation; two extreme test statistics are not shown, those for Crohns' Disease & Inflammatory Bowel Disease ($x$=1211,$y$=1357) and Ulcerative Colitis & Inflammatory Bowel Disease ($x$=739,$y$=2380). We see that while LDSC and SumHer-GC estimates of genetic correlation are highly concordant (correlation 0.94), the latter are on average a third more precise

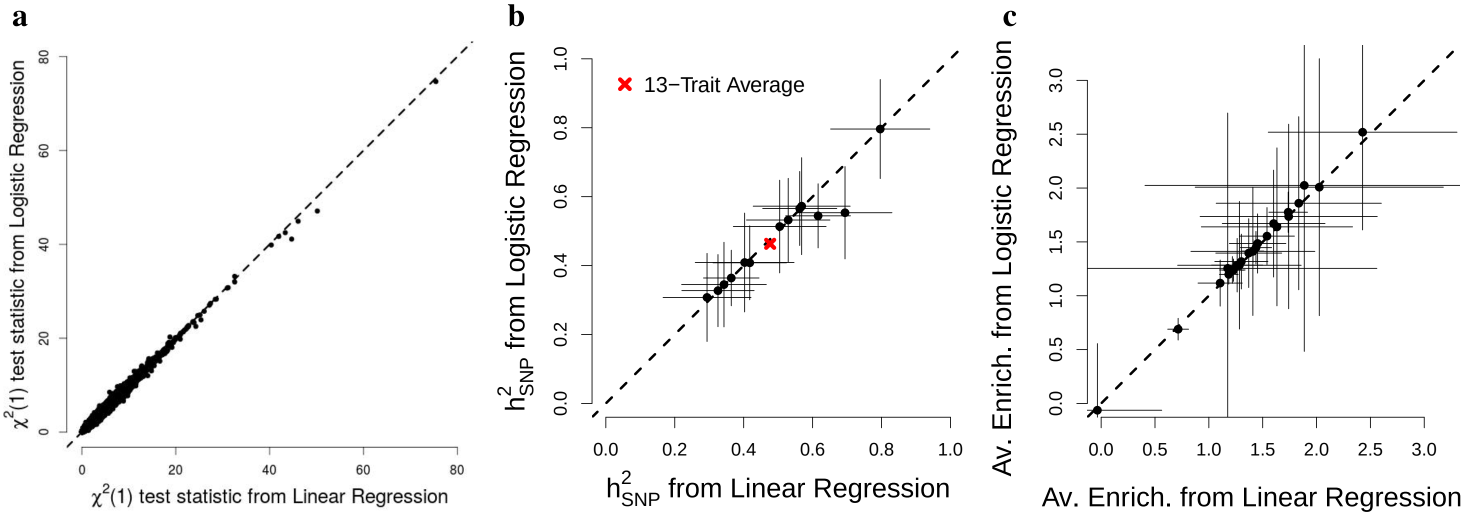


**Supplementary Figure 10: Comparing estimates from linear and logistic regression for the WTCCC GWAS.** SumHer is designed to be used with summary statistics from (classical) linear regression. However, for binary traits (e.g., case-control studies), its estimates will in general be very similar if instead summary statistics from logistic regression are used. This is because for SNPs with small or moderate effect there will be high concordance between test statistics from linear and logistic regression (while the two methods can produce contrasting test statistics for large-effect SNPs, we recommend excluding these when running SumHer). **(a)** Points compare test statistics from logistic regression ($y$-axis) to those from linear regression ($x$-axis); we focus on approximately 60 000 SNPs per trait, obtained by first thinning (within 1 cM and $r^2_{jl} > 0.2$), then excluding SNPs in the major histocompatibility complex (Chromosome 6: 25-34 Mb), as well as SNPs which individually explain >1% of phenotypic variation and SNPs in LD with these (within 1 cM and $r^2_{jl} > 0.1$). **(b)** Points compare estimates of $h^2_{\text{SNP}}$ from SumHer-Zero using test statistics from logistic regression ($y$-axis) to those from linear regression ($x$-axis). **(c)** For the 24 functional categories, points compare the average estimates of enrichment from SumHer-Zero using test statistics from logistic regression ($y$-axis) to those from linear regression ($x$-axis).

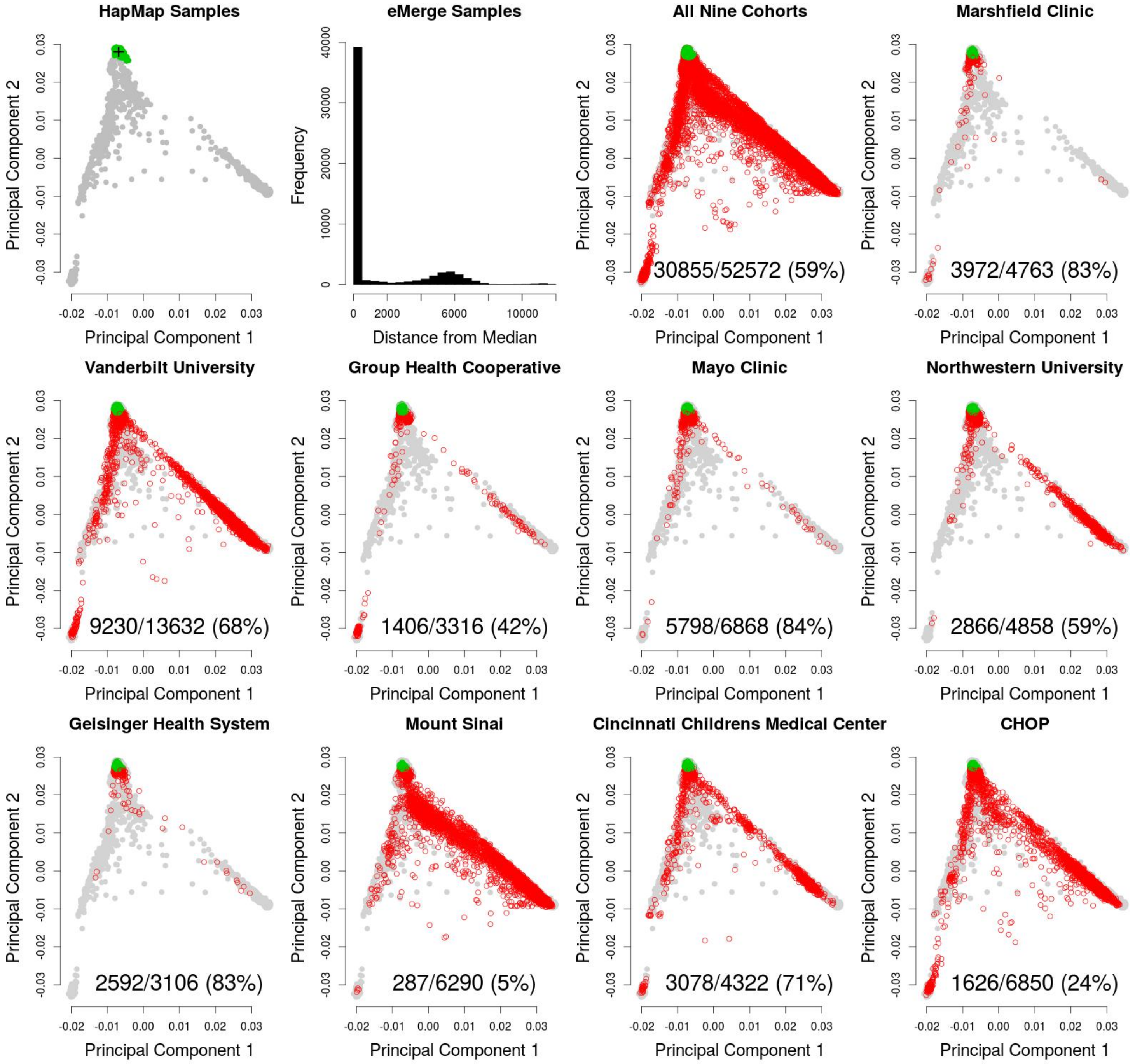


**Supplementary Figure 11: Sample quality control for eMerge data.** The post-imputed eMerge data contains 52 572 individuals. To filter based on ancestry, we perform principal component analysis (PCA) on the 2 404 individuals from the 1000 Genome Project. **Panel 1** plots the first two PCs for the 1000 Genome Project individuals. The 404 non-Finnish Europeans are marked in green; the center of the cross indicates their median $(M_x, M_y)$, while the horizontal and vertical line segments mark 95% confidence intervals (with widths $3.96\,S_x$ and $3.96\,S_y$, respectively). If $(P_{xi}, P_{yo})$ denotes the projection of the $i$th eMerge individual onto these top two PCs, then we compute $D_i^2 = (P_{xi} - M_x)^2/S_x^2 + (P_{yi} - M_y)^2/S_y^2$, the square of the standardized distance between the individual and the median. We will exclude individuals with $D_i^2 > 5.99$, the 95th percentile of the $\chi^2(2)$ distribution. **Panel 2** shows the distribution of $D_i^2$ for the eMerge individuals. **Panel 3** plots $D_i^2$ for all eMerge individuals; the numbers indicate how many of the individuals have $D_i^2 < 5.99$ (these individuals are marked in green). **Panels 4-12** plot the same for each of the nine cohorts separately; we decide to exclude individuals from Mount Sinai and CHOP (Children's Hospital of Philadelphia), as for these two cohorts, fewer than 25% of individuals pass our test. In addition to filtering based on $D_i^2$, we excluded 125 individuals reported as having "Hispanic or Latino" ethnicity, then filtered individuals so that no pair remained with estimated relatedness $> 0.05$, (which left 25 875 individuals).

| Category Annotation | Average LD | Expected share of $h^2_{\text{SNP}}$ GCTA | LDAK |
|---|---|---|---|
| Coding[16,17] | 127.0 | 0.015 | 0.019 |
| Conserved[18,19] | 105.9 | 0.026 | 0.033 |
| CTCF[20] | 113.2 | 0.020 | 0.025 |
| Digital Genomic Footprint[17,21] | 109.8 | 0.124 | 0.152 |
| DNase I Hypersensitive Site[21–23] | 105.6 | 0.157 | 0.193 |
| FANTOM5 Enhancer[24] | 98.6 | 0.003 | 0.005 |
| Enhancer[20] | 104.7 | 0.034 | 0.047 |
| Fetal DHS[21–23] | 101.9 | 0.076 | 0.098 |
| H3K27ac (Hnisz)[22,25] | 102.9 | 0.348 | 0.436 |
| H3K27ac (PGC2)[22,26] | 107.9 | 0.241 | 0.300 |
| H3K4me1[22,23] | 106.5 | 0.399 | 0.481 |
| H3K4me3[22,23] | 141.5 | 0.119 | 0.143 |
| H3K9ac[22,23] | 106.4 | 0.107 | 0.141 |
| Intronic[16,17] | 119.7 | 0.406 | 0.412 |
| Promoter Flanking[20] | 129.1 | 0.008 | 0.009 |
| Promoter[16,17] | 146.2 | 0.029 | 0.035 |
| Repressed[20] | 134.7 | 0.471 | 0.448 |
| Super Enhancer[25] | 94.3 | 0.138 | 0.193 |
| Transcription Factor Binding Site[17,21] | 110.4 | 0.117 | 0.146 |
| Transcribed[20] | 129.8 | 0.368 | 0.351 |
| Transcription Start Site[20] | 125.5 | 0.014 | 0.018 |
| 3' Untranslated Region[16,17] | 117.0 | 0.012 | 0.015 |
| 5' Untranslated Region[16,17] | 129.0 | 0.005 | 0.006 |
| Weak Enhancer[20] | 97.7 | 0.016 | 0.024 |

**Supplementary Table 1: Details of the 24 functional annotations.** When estimating enrichments of SNP categories, we use the same annotations as Finucane *et al.*, which can be downloaded from https://data.broadinstitute.org/alkesgroup/LDSCORE/baseline_bedfiles.tgz. This table summarizes the 24 categories. We measure the LD of SNP $j$ by $\sum_{l \in N_j} r^2_{jl}$, where $N_j$ indexes the SNPs within 1 cM and $r_{jk}$ is (an estimate of) the squared correlation between SNPs $j$ and $l$. Given a heritability model, the expected proportion of $h^2_{\text{SNP}}$ contributed by Category $k$ is $(\sum_j I_{jk} q_j)/(\sum_j q_j)$, where $I_{jk}$ indicates whether SNP $j$ belongs to that category (under the GCTA Model, this fraction represents the proportion of SNPs in the category). The majority of categories contain SNPs whose average LD is lower than the average LD of all SNPs (133.7), which is why most categories are expected to contribute more under the LDAK Model than under the GCTA Model.

| Trait | $n$ | $m$ | GIF | LDSC-Zero $h^2_{SNP}$ (SD) | LDSC $h^2_{SNP}$ (SD) | LDSC $1+A$ (SD) | SumHer-Zero $h^2_{SNP}$ (SD) | SumHer-GC $h^2_{SNP}$ (SD) | SumHer-GC $C$ (SD) |
|---|---|---|---|---|---|---|---|---|---|
| Bipolar Disorder | 4776 | 1937638 | 1.036 | 0.42 (0.05) | 0.23 (0.05) | 1.042 (0.009) | 0.80 (0.07) | 1.09 (0.16) | 0.971 (0.013) |
| Coronary Artery Disease | 4845 | 1943684 | 1.016 | 0.20 (0.04) | 0.13 (0.05) | 1.018 (0.008) | 0.34 (0.06) | 0.58 (0.12) | 0.975 (0.012) |
| Crohn's Disease | 4641 | 1932019 | 1.047 | 0.23 (0.05) | 0.05 (0.07) | 1.046 (0.009) | 0.57 (0.07) | 0.64 (0.15) | 0.993 (0.012) |
| Hypertension | 4853 | 1944541 | 1.042 | 0.21 (0.04) | 0.08 (0.04) | 1.031 (0.009) | 0.50 (0.07) | 0.71 (0.13) | 0.979 (0.012) |
| Rheumatoid Arthritis | 4769 | 1941327 | 1.035 | 0.08 (0.04) | -0.01 (0.04) | 1.022 (0.007) | 0.29 (0.07) | 0.45 (0.13) | 0.984 (0.012) |
| Type I Diabetes | 4878 | 1937840 | 1.024 | 0.16 (0.04) | 0.08 (0.05) | 1.022 (0.009) | 0.40 (0.07) | 0.69 (0.16) | 0.970 (0.014) |
| Type II Diabetes | 4829 | 1940920 | 1.040 | 0.26 (0.04) | 0.13 (0.05) | 1.031 (0.008) | 0.53 (0.06) | 0.66 (0.13) | 0.987 (0.012) |
| Barrett's Oesophagus | 7037 | 2887629 | 1.035 | 0.17 (0.04) | 0.06 (0.04) | 1.029 (0.007) | 0.42 (0.05) | 0.48 (0.10) | 0.993 (0.010) |
| Celiac Disease | 8970 | 2876170 | 1.049 | 0.18 (0.03) | 0.09 (0.03) | 1.043 (0.009) | 0.36 (0.04) | 0.44 (0.08) | 0.985 (0.013) |
| Ischaemic Stroke | 6859 | 2879412 | 1.058 | 0.26 (0.03) | -0.00 (0.03) | 1.078 (0.007) | 0.62 (0.05) | 0.12 (0.09) | 1.067 (0.010) |
| Parkinson's Disease | 7462 | 2877852 | 1.010 | 0.18 (0.04) | 0.16 (0.06) | 1.004 (0.008) | 0.33 (0.05) | 0.56 (0.12) | 0.976 (0.010) |
| Psoriasis | 8020 | 2843806 | 1.068 | 0.32 (0.04) | 0.10 (0.04) | 1.056 (0.008) | 0.69 (0.07) | 0.60 (0.11) | 1.010 (0.010) |
| Ulcerative Colitis | 9934 | 1875844 | 1.031 | 0.25 (0.04) | 0.08 (0.04) | 1.050 (0.007) | 0.56 (0.05) | 0.64 (0.11) | 0.991 (0.011) |
| Age-related Macular Disease | 8475 | 2972162 | 1.027 | 0.06 (0.03) | -0.00 (0.04) | 1.023 (0.007) | 0.19 (0.05) | 0.03 (0.08) | 1.020 (0.010) |
| Heart Failure | 9005 | 2972162 | 1.022 | 0.07 (0.02) | -0.04 (0.02) | 1.034 (0.006) | 0.18 (0.04) | -0.04 (0.07) | 1.030 (0.009) |
| Peripheral Arterial Disease | 10541 | 2972162 | 1.044 | 0.09 (0.02) | 0.01 (0.02) | 1.027 (0.006) | 0.18 (0.04) | 0.06 (0.07) | 1.019 (0.011) |
| Shingles (Herpes Zoster) | 11966 | 2972162 | 1.004 | 0.03 (0.02) | 0.04 (0.02) | 0.994 (0.007) | 0.02 (0.03) | 0.03 (0.05) | 0.998 (0.009) |
| Venous Thromboembolism | 13961 | 2972162 | 1.031 | 0.06 (0.02) | 0.03 (0.03) | 1.017 (0.008) | 0.15 (0.03) | 0.15 (0.07) | 1.001 (0.011) |
| Triglyceride | 12137 | 2972162 | 1.031 | 0.14 (0.02) | 0.06 (0.03) | 1.030 (0.007) | 0.24 (0.04) | 0.22 (0.07) | 1.004 (0.011) |
| LDL Cholesterol | 13420 | 2972162 | 1.020 | 0.07 (0.02) | 0.01 (0.02) | 1.025 (0.007) | 0.15 (0.03) | 0.08 (0.05) | 1.014 (0.010) |
| HDL Cholesterol | 13788 | 2972162 | 1.050 | 0.16 (0.02) | 0.05 (0.03) | 1.047 (0.009) | 0.33 (0.03) | 0.27 (0.06) | 1.011 (0.011) |
| Systolic Blood Pressure | 15058 | 2972162 | 1.017 | 0.04 (0.02) | 0.01 (0.02) | 1.017 (0.007) | 0.12 (0.03) | 0.09 (0.05) | 1.007 (0.010) |
| Diastolic Blood Pressure | 15062 | 2972162 | 1.028 | 0.04 (0.02) | 0.00 (0.02) | 1.020 (0.006) | 0.11 (0.02) | 0.02 (0.05) | 1.018 (0.010) |
| Height | 18152 | 2972162 | 1.066 | 0.25 (0.02) | 0.17 (0.03) | 1.043 (0.009) | 0.42 (0.03) | 0.63 (0.07) | 0.952 (0.013) |
| Body Mass Index | 19309 | 2972162 | 1.064 | 0.20 (0.02) | 0.12 (0.02) | 1.048 (0.008) | 0.34 (0.02) | 0.42 (0.05) | 0.979 (0.011) |
| **Average** | **9710** | **2619385** | **1.034** | **0.12 (0.01)** | **0.04 (0.01)** | **1.031 (0.002)** | **0.26 (0.01)** | **0.23 (0.02)** | **1.001 (0.002)** |
| **Relative** | | | | **1** | **0.39 (0.04)** | | **1.98 (0.05)** | **2.10 (0.10)** | |
| **Relative** | | | | **0.48 (0.02)** | **0.17 (0.02)** | | **1** | **1.00 (0.05)** | |

**Supplementary Table 2: Details of the 25 raw GWAS, and estimates of $h^2_{SNP}$ and confounding bias.** $n$ denotes the number of samples, $m$ the number of SNPs, GIF the genomic inflation factor. For each trait, we report estimates of $h^2_{SNP}$ from LDSC-Zero, LDSC, SumHer-Zero and SumHer-GC, as well as estimates of confounding bias from LDSC and SumHer-GC (LDSC measures bias via the intercept $1+A$, while SumHer-GC estimates the scaling factor $C$). For binary traits, estimates of $h^2_{SNP}$ are on the observed scale. The 18 traits with significant $h^2_{SNP}$ ($P < 0.05/25$) from both LDSC-Zero and SumHer-Zero are marked in red (these are the ones we restricted to when estimating genetic correlations).

| Category | LDSC-Zero (53-part model) Share (SD) | Expected | Av. Enrichment (SD) | SumHer-Zero (25-part model) Share (SD) | Expected | Av. Enrichment (SD) |
|---|---|---|---|---|---|---|
| Coding | 0.086 (0.020) | 0.013 | 6.77 (1.58) | 0.029 (0.005) | 0.017 | 1.75 (0.30) |
| Conserved | 0.277 (0.036) | 0.025 | 11.05 (1.42) | 0.051 (0.007) | 0.031 | 1.65 (0.23) |
| CTCF | 0.021 (0.030) | 0.019 | 1.12 (1.58) | 0.010 (0.006) | 0.023 | 0.44 (0.24) |
| Digital Genomic Footprint | 0.405 (0.077) | 0.120 | 3.38 (0.64) | 0.191 (0.014) | 0.145 | 1.32 (0.09) |
| DNase I Hypersensitive Site | 0.365 (0.078) | 0.152 | 2.40 (0.51) | 0.219 (0.015) | 0.186 | 1.18 (0.08) |
| FANTOM5 Enhancer | 0.034 (0.014) | 0.003 | 10.53 (4.39) | 0.003 (0.002) | 0.004 | 0.55 (0.55) |
| Enhancer | 0.083 (0.030) | 0.032 | 2.58 (0.95) | 0.064 (0.008) | 0.043 | 1.48 (0.19) |
| Fetal DHS | 0.150 (0.059) | 0.073 | 2.06 (0.81) | 0.123 (0.011) | 0.094 | 1.30 (0.12) |
| H3K27ac (Hnisz) | 0.664 (0.031) | 0.334 | 1.99 (0.09) | 0.589 (0.016) | 0.421 | 1.40 (0.04) |
| H3K27ac (PGC2) | 0.526 (0.049) | 0.231 | 2.29 (0.21) | 0.404 (0.017) | 0.290 | 1.39 (0.06) |
| H3K4me1 | 0.817 (0.048) | 0.387 | 2.11 (0.13) | 0.604 (0.018) | 0.469 | 1.29 (0.04) |
| H3K4me3 | 0.248 (0.039) | 0.112 | 2.22 (0.35) | 0.207 (0.014) | 0.133 | 1.55 (0.10) |
| H3K9ac | 0.297 (0.041) | 0.099 | 3.02 (0.42) | 0.227 (0.013) | 0.130 | 1.75 (0.10) |
| Intronic | 0.490 (0.022) | 0.413 | 1.19 (0.05) | 0.504 (0.014) | 0.414 | 1.22 (0.03) |
| Promoter Flanking | -0.003 (0.019) | 0.008 | -0.45 (2.43) | 0.015 (0.004) | 0.008 | 1.79 (0.48) |
| Promoter | 0.031 (0.018) | 0.026 | 1.17 (0.69) | 0.035 (0.007) | 0.030 | 1.17 (0.21) |
| Repressed | 0.341 (0.046) | 0.475 | 0.72 (0.10) | 0.305 (0.018) | 0.457 | 0.67 (0.04) |
| Super Enhancer | 0.310 (0.018) | 0.129 | 2.40 (0.14) | 0.302 (0.012) | 0.181 | 1.66 (0.07) |
| Transcription Factor Binding Site | 0.310 (0.058) | 0.112 | 2.74 (0.52) | 0.182 (0.014) | 0.138 | 1.31 (0.10) |
| Transcribed | 0.381 (0.042) | 0.375 | 1.01 (0.11) | 0.464 (0.017) | 0.353 | 1.31 (0.05) |
| Transcription Start Site | 0.053 (0.021) | 0.012 | 4.42 (1.72) | 0.035 (0.005) | 0.016 | 2.27 (0.32) |
| 3' Untranslated Region | 0.050 (0.015) | 0.010 | 4.74 (1.48) | 0.024 (0.004) | 0.014 | 1.76 (0.32) |
| 5' Untranslated Region | 0.003 (0.012) | 0.005 | 0.42 (2.64) | 0.001 (0.003) | 0.005 | 0.29 (0.53) |
| Weak Enhancer | 0.093 (0.028) | 0.015 | 6.02 (1.88) | 0.041 (0.006) | 0.022 | 1.87 (0.28) |

**Supplementary Table 3: Average estimates of enrichment from the 25 raw GWAS.** We estimate enrichments using either LDSC-Zero with a 53-part model or SumHer-Zero with a 25-part model. For the 25-part model, we divide the genome into a set for each of the 24 categories, plus a set containing all other SNPs, while for the 53-part model, we divide the genome into a set for each of the 24 categories, a set for each of 28 buffer regions (see Finucane *et al.*,[9]), plus a set containing all other SNPs. For each trait, we calculate the estimated enrichment of each category, obtained by dividing the estimated share of $h^2_{\text{SNP}}$ contributed by the category by its expected share; for each category, we then report the (inverse-weighted) average estimated enrichment across the 25 traits. Average estimated enrichments significantly different from 1 ($p < 0.05$) are marked in red.

| Trait 1 | Trait 2 | LDSC-Zero Correlation (SD) | LDSC Correlation (SD) | SumHer-Zero Correlation (SD) | SumHer-GC Correlation (SD) |
|---|---|---|---|---|---|
| Bipolar Disorder | Crohn's Disease | 0.19 (0.12) | 0.48 (1.11) | 0.41 (0.17) | 0.37 (0.12) |
| Bipolar Disorder | Hypertension | 0.18 (0.13) | 0.38 (0.25) | 0.46 (0.15) | **0.37 (0.10)** |
| Bipolar Disorder | Type I Diabetes | 0.15 (0.11) | 0.30 (0.19) | 0.27 (0.14) | 0.23 (0.10) |
| Bipolar Disorder | Psoriasis | 0.07 (0.14) | 0.16 (0.31) | 0.44 (0.16) | 0.40 (0.14) |
| Coronary Artery Disease | Rheumatoid Arthritis | 0.42 (0.18) | 0.75 (0.33) | 0.22 (0.23) | 0.16 (0.14) |
| Coronary Artery Disease | Type I Diabetes | 0.20 (0.15) | 0.38 (0.22) | 0.37 (0.20) | 0.29 (0.13) |
| Coronary Artery Disease | Parkinson's Disease | 0.07 (0.13) | 0.14 (0.28) | 0.46 (0.18) | 0.42 (0.14) |
| Crohn's Disease | Psoriasis | 0.25 (0.18) | 0.94 (0.85) | **0.89 (0.19)** | **0.82 (0.19)** |
| Crohn's Disease | Shingles (Herpes Zoster) | -0.18 (0.07) | -1.05 (NaN) | -0.09 (0.10) | -0.09 (0.09) |
| Hypertension | Type I Diabetes | 0.11 (0.15) | 0.23 (0.30) | 0.31 (0.16) | 0.27 (0.12) |
| Hypertension | Psoriasis | 0.11 (0.18) | 0.27 (0.52) | 0.53 (0.19) | 0.61 (0.21) |
| Rheumatoid Arthritis | Barrett's Oesophagus | 0.52 (0.17) | 1.31 (0.84) | 0.33 (0.19) | 0.29 (0.16) |
| Rheumatoid Arthritis | Ischaemic Stroke | 0.42 (0.23) | 0.80 (0.42) | 0.71 (0.27) | 0.48 (0.17) |
| Rheumatoid Arthritis | Parkinson's Disease | 0.09 (0.16) | 0.24 (0.42) | 0.39 (0.17) | 0.38 (0.16) |
| Rheumatoid Arthritis | Venous Thromboembolism | 0.24 (0.10) | 0.67 (0.48) | 0.08 (0.11) | 0.06 (0.09) |
| Type II Diabetes | Barrett's Oesophagus | -0.33 (0.15) | -0.92 (4.83) | 0.04 (0.16) | 0.03 (0.14) |
| Type II Diabetes | Celiac Disease | -0.14 (0.14) | -2.51 (NaN) | -0.29 (0.15) | -0.70 (0.58) |
| Type II Diabetes | Ischaemic Stroke | 0.42 (0.18) | 0.82 (0.40) | -0.31 (0.22) | -0.24 (0.18) |
| Type II Diabetes | Psoriasis | -0.11 (0.25) | NaN (NaN) | 0.40 (0.20) | 0.39 (0.19) |
| Barrett's Oesophagus | Ischaemic Stroke | -0.02 (0.15) | 0.02 (0.28) | 0.47 (0.22) | 0.31 (0.14) |
| Barrett's Oesophagus | Parkinson's Disease | 0.01 (0.10) | 0.07 (0.22) | 0.33 (0.12) | 0.32 (0.12) |
| Celiac Disease | Shingles (Herpes Zoster) | -0.13 (0.06) | NaN (NaN) | -0.04 (0.08) | -0.10 (0.18) |
| Ischaemic Stroke | Psoriasis | 0.26 (0.20) | 0.54 (0.38) | 0.55 (0.23) | 0.44 (0.18) |
| Parkinson's Disease | Psoriasis | 0.17 (0.13) | 0.45 (0.35) | 0.46 (0.16) | 0.45 (0.15) |
| Parkinson's Disease | Venous Thromboembolism | -0.01 (0.06) | 0.00 (0.16) | -0.17 (0.08) | -0.19 (0.10) |
| Psoriasis | Age-related Macular Disease | -0.24 (0.12) | -0.51 (0.32) | -0.12 (0.15) | -0.11 (0.13) |
| Age-related Macular Disease | Peripheral Arterial Disease | -0.09 (0.12) | -0.18 (0.28) | **0.75 (0.17)** | 0.86 (0.30) |
| Age-related Macular Disease | Venous Thromboembolism | 0.22 (0.10) | 0.38 (0.19) | 0.08 (0.13) | 0.08 (0.12) |
| Heart Failure | Peripheral Arterial Disease | 0.16 (0.15) | 0.57 (0.62) | 0.37 (0.18) | 0.60 (0.37) |
| Heart Failure | Venous Thromboembolism | -0.15 (0.13) | -0.40 (0.42) | -0.41 (0.16) | -0.55 (0.32) |
| Peripheral Arterial Disease | Venous Thromboembolism | -0.26 (0.09) | -0.56 (0.20) | **-0.51 (0.11)** | **-0.51 (0.10)** |

**Supplementary Table 4: Estimates of genetic correlation for the 25 raw GWAS.** In general, it is only possible to get a meaningful estimate of genetic correlation when both traits have substantial $h^2_{\text{SNP}}$, so for this analysis we only use the 18 traits for which both LDSC-Zero and SumHer-Zero find significant $h^2_{\text{SNP}}$ ($P < 0.05/25$); this filtering excludes rheumatoid arthritis, age-related macular disease, heart failure, shingles, venous thromboembolism and both systolic and diastolic 5ressure (Supplementary Table 2). Of the ${}^{18}C_2 = 153$ pairs of traits we considered, this table reports genetic correlations for the 31 pairs with significant correlation $P < 0.05$) from either LDSC-Zero, LDSC, SumHer-Zero or SumHer-GC. Nominally significant estimates ($P < 0.05$) are marked in red, while Bonferonni significant estimates ($P < 0.05/153$) are also in **bold**. Some LDSC estimates are missing (NaN). When a genetic correlation is missing, this indicates that one of the two traits is estimated to have negative $h^2_{\text{SNP}}$ (as then $\sqrt{h^2_{\text{SNP}}}$ can not be computed); when a SD is missing, this indicates that one or more negative $h^2_{\text{SNP}}$ occurred during the jackknifing.

| Trait | GCTA Model | | LDAK Model | | Schoech *et al.* Model | |
|---|---|---|---|---|---|---|
| | $h^2_{\text{SNP}}$ (SD) | Log Likelihood | $h^2_{\text{SNP}}$ (SD) | Log Likelihood | $h^2_{\text{SNP}}$ (SD) | Log Likelihood |
| Bipolar Disorder | 0.29 (0.04) | -3253.4 | 0.59 (0.06) | -3229.7 | 0.44 (0.04) | -3234.6 |
| Coronary Artery Disease | 0.14 (0.03) | -3148.1 | 0.30 (0.06) | -3143.9 | 0.24 (0.04) | -3141.1 |
| Crohn's Disease | 0.16 (0.03) | -3147.1 | 0.43 (0.06) | -3128.0 | 0.30 (0.04) | -3133.2 |
| Hypertension | 0.14 (0.03) | -3367.9 | 0.40 (0.06) | -3350.1 | 0.26 (0.04) | -3357.7 |
| Rheumatoid Arthritis | 0.05 (0.03) | -3185.7 | 0.24 (0.06) | -3177.3 | 0.13 (0.04) | -3181.3 |
| Type I Diabetes | 0.12 (0.03) | -3419.4 | 0.30 (0.05) | -3409.8 | 0.21 (0.04) | -3413.1 |
| Type II Diabetes | 0.17 (0.03) | -3340.5 | 0.40 (0.06) | -3328.7 | 0.28 (0.04) | -3331.6 |
| Barrett's Oesophagus | 0.12 (0.03) | -3876.8 | 0.34 (0.05) | -3858.6 | 0.21 (0.04) | -3869.0 |
| Celiac Disease | 0.08 (0.01) | -5514.0 | 0.26 (0.03) | -5489.7 | 0.17 (0.02) | -5497.6 |
| Ischaemic Stroke | 0.18 (0.02) | -5615.9 | 0.46 (0.04) | -5565.2 | 0.26 (0.03) | -5605.1 |
| Parkinson's Disease | 0.13 (0.03) | -3884.4 | 0.28 (0.05) | -3882.2 | 0.18 (0.04) | -3885.9 |
| Psoriasis | 0.21 (0.03) | -4439.2 | 0.50 (0.05) | -4398.9 | 0.32 (0.03) | -4422.1 |
| Ulcerative Colitis | 0.19 (0.03) | -5230.5 | 0.46 (0.04) | -5188.9 | 0.30 (0.03) | -5209.9 |
| Age-related Macular Disease | 0.06 (0.03) | -2989.7 | 0.19 (0.04) | -2982.3 | 0.08 (0.03) | -2989.2 |
| Heart Failure | 0.07 (0.03) | -4770.3 | 0.18 (0.04) | -4762.5 | 0.10 (0.03) | -4767.4 |
| Peripheral Arterial Disease | 0.09 (0.02) | -4597.6 | 0.19 (0.03) | -4588.4 | 0.11 (0.03) | -4595.3 |
| Shingles (Herpes Zoster) | 0.03 (0.02) | -2394.3 | 0.01 (0.03) | -2395.7 | 0.03 (0.02) | -2394.6 |
| Venous Thromboembolism | 0.06 (0.02) | -5638.5 | 0.16 (0.03) | -5628.5 | 0.07 (0.02) | -5638.3 |
| Triglyceride | 0.14 (0.02) | -71357.2 | 0.23 (0.03) | -71350.4 | 0.16 (0.02) | -71353.6 |
| LDL Cholesterol | 0.07 (0.02) | -64288.3 | 0.15 (0.03) | -64278.1 | 0.08 (0.02) | -64287.2 |
| HDL Cholesterol | 0.16 (0.02) | -56038.1 | 0.32 (0.03) | -56001.6 | 0.19 (0.02) | -56029.0 |
| Systolic Blood Pressure | 0.04 (0.02) | -60425.9 | 0.12 (0.02) | -60415.4 | 0.05 (0.02) | -60425.3 |
| Diastolic Blood Pressure | 0.04 (0.02) | -52671.9 | 0.10 (0.02) | -52665.4 | 0.05 (0.02) | -52671.2 |
| Height | 0.23 (0.02) | -60517.5 | 0.38 (0.02) | -60462.2 | 0.28 (0.02) | -60482.1 |
| Body Mass Index | 0.19 (0.01) | -64108.2 | 0.31 (0.02) | -64066.4 | 0.23 (0.02) | -64070.9 |
| **Average** | **0.12 (0.00)** | **-20048.8** | **0.25 (0.01)** | **-20029.9** | **0.16 (0.01)** | **-20039.4** |
| **Difference** | | **0** | | **18.9** | | **9.4** |

**Supplementary Table 5: Comparing heritability models for the 25 raw GWAS using REML.** For each trait, we compare the performance of the GCTA, LDAK and Schoech *et al.* Models based on likelihood from REML;[6] for consistency with our SumHer analyses, when calculating kinship matrices we excluded SNPs in the major histocompatibility complex (Chromosome 6: 25-34 Mb), as well as SNPs which individually explain >1% of phenotypic variation and SNPs in LD with these (within 1 cM and $r^2_{jl} > 0.1$). For each trait, the model resulting in the highest likelihood is shown in red.

| Trait | $L(S_j \mid \widehat{h^2_{\text{SNP}}}, D^0)$ GCTA | LDAK | $L(S_j \mid \widehat{h^2_{\text{SNP}}}, D^{\widehat{h^2_{\text{SNP}}}}) - L(S_j \mid \widehat{h^2_{\text{SNP}}}, D^0)$ GCTA | LDAK | Proportion of LDAK, $p$ (SD) |
|---|---|---|---|---|---|
| Bipolar Disorder | -77350.6 | -77321.6 | 34.5 | 57.9 | 0.93 (0.07) |
| Coronary Artery Disease | -75691.8 | -75685.6 | 10.7 | 13.6 | 0.90 (0.14) |
| Crohn's Disease | -76535.7 | -76515.3 | 12.1 | 30.4 | 1.08 (0.14) |
| Hypertension | -75942.1 | -75927.4 | 11.5 | 27.7 | 1.06 (0.10) |
| Rheumatoid Arthritis | -75316.4 | -75310.2 | 1.9 | 9.8 | 1.28 (0.12) |
| Type I Diabetes | -76441.0 | -76431.2 | 7.4 | 18.1 | 1.05 (0.15) |
| Type II Diabetes | -76350.4 | -76339.0 | 15.9 | 29.1 | 0.94 (0.11) |
| Barrett's Oesophagus | -108847.8 | -108833.4 | 11.6 | 27.4 | 1.09 (0.11) |
| Celiac Disease | -85466.4 | -85444.3 | 25.6 | 46.3 | 0.95 (0.10) |
| Ischaemic Stroke | -110924.2 | -110883.2 | 34.1 | 80.9 | 1.08 (0.06) |
| Parkinson's Disease | -107955.1 | -107959.9 | 11.7 | 16.6 | 0.65 (0.30) |
| Psoriasis | -112302.1 | -112264.8 | 34.6 | 70.6 | 1.05 (0.07) |
| Ulcerative Colitis | -107001.2 | -106960.5 | 27.6 | 59.9 | 1.15 (0.06) |
| Age-related Macular Disease | -112084.4 | -112081.2 | 2.5 | 8.2 | 1.13 (0.24) |
| Heart Failure | -111871.7 | -111866.5 | 3.0 | 8.3 | 1.27 (0.16) |
| Peripheral Arterial Disease | -111627.4 | -111621.9 | 6.5 | 11.1 | 1.08 (0.15) |
| Shingles (Herpes Zoster) | -110061.7 | -110063.0 | 1.4 | 0.3 | -11.20 (1449.09) |
| Venous Thromboembolism | -111792.2 | -111788.5 | 4.2 | 10.2 | 1.04 (0.23) |
| Triglyceride | -116336.4 | -116330.3 | 17.1 | 24.1 | 0.85 (0.14) |
| LDL Cholesterol | -112292.2 | -112286.9 | 6.0 | 13.3 | 1.08 (0.15) |
| HDL Cholesterol | -117197.3 | -117172.7 | 27.4 | 51.6 | 1.05 (0.09) |
| Systolic Blood Pressure | -111024.6 | -111021.5 | 2.8 | 10.3 | 1.09 (0.22) |
| Diastolic Blood Pressure | -111336.7 | -111333.8 | 3.1 | 8.3 | 1.09 (0.26) |
| Height | -118819.3 | -118768.4 | 89.2 | 128.2 | 0.90 (0.07) |
| Body Mass Index | -117139.7 | -117087.9 | 74.0 | 101.3 | 0.94 (0.06) |
| **Average** | **-101108.3** | **-101092.0** | **19.0** | **34.5** | **1.03 (0.02)** |
| **Difference** | **0** | **16.4** | | | |

**Supplementary Table 6: Comparing heritability models for the 25 raw GWAS using SumHer.** $L(S_j|h^2_{\text{SNP}}, D)$ is the (weighted) log likelihood, defined in Online Methods. SumHer provides two likelihood-based metrics for comparing heritability models. Our preferred metric is $L(S_j|\widehat{h^2_{\text{SNP}}}, D^0)$, where $\widehat{h^2_{\text{SNP}}}$ is the final estimate of $h^2_{\text{SNP}}$, and $D^0$ is the initial weight matrix (obtained by setting $1/D_{jj} = \sum_{l \in N_j} r^2_{jl}$). The second metric is $L(S_j|\widehat{h^2_{\text{SNP}}}, D^{\widehat{h^2_{\text{SNP}}}}) - L(S_j|\widehat{h^2_{\text{SNP}}}, D^0)$, where $D^{\widehat{h^2_{\text{SNP}}}}$ is the weight matrix after the final iteration. While it might seem natural to compare heritability models based on $L(S_j|\widehat{h^2_{\text{SNP}}}, D^{\widehat{h^2_{\text{SNP}}}})$, this is not valid because $D^{\widehat{h^2_{\text{SNP}}}}$ depends on the heritability model, hence why we normalize by subtracting $L(S_j|0, D^{\widehat{h^2_{\text{SNP}}}})$, the null likelihood calculated using the same weights (the result is equal to half the likelihood ratio test statistic). We see that regardless of which metric is used, the relative performances of the GCTA and LDAK Models are very similar to when we compare models using the log likelihood from REML (Columns 3 & 5 of Supplementary Table 5). An alternative way to compare the GCTA and LDAK Models is using Hybrid-Zero, which assumes the heritability model $q_j = (1-p) \times 1/m + p \times [f_j(1-f_j)]^{0.75}\, w_j/Q'$, where $Q' = \sum_j [f_j(1-f_j)]^{0.75}\, w_j$. The final column reports estimates of $p$, the LDAK proportion. The very imprecise estimate for Shingles reflects that this trait has very low $h^2_{\text{SNP}}$.

Note that when comparing heritability models based on $L(S_j|\widehat{h^2_{\text{SNP}}}, D^0)$, we must ensure the same SNPs are used for each model (so that $D^0$ is constant). When calculating the tagfile (which contains $q_j + \sum_{l \in N_j} q_l r^2_{jl}$ for each SNP), by default SumHer excludes SNPs with $q_0 = 0$. This is not relevant for the GCTA Model ($q_j = 0$), but is for the LDAK Model ($q_j = w_j\ [f_j(1-f_j)]^{0.75}$), as many SNPs will have $w_j = 0$ (these are SNPs whose variation is perfectly tagged by their neighbors). To prevent SumHer ignoring SNPs with $q_j = 0$, use the option `--reduce NO` (see the Supplementary Note for more details).

| Trait | Original Controls — Confounding (SD) | | | 2 404 POBI Controls — Confounding (SD) | | | 2 404 1000G Controls — Confounding (SD) | | |
|---|---|---|---|---|---|---|---|---|---|
| | **GIF** | **LDSC (SD)** | **SumHer-GC (SD)** | **GIF** | **LDSC (SD)** | **SumHer-GC (SD)** | **GIF** | **LDSC (SD)** | **SumHer-GC (SD)** |
| Bipolar Disorder | 1.032 | 1.033 (0.009) | 0.972 (0.014) | 1.129 | 1.116 (0.010) | 1.082 (0.015) | 65.58 | 79.70 (0.70) | 83.31 (1.00) |
| Coronary Artery Disease | 1.010 | 1.017 (0.009) | 0.978 (0.012) | 1.090 | 1.092 (0.010) | 1.066 (0.013) | 69.09 | 83.49 (0.73) | 87.32 (1.03) |
| Crohn's Disease | 1.046 | 1.044 (0.009) | 1.000 (0.014) | 1.097 | 1.121 (0.010) | 1.065 (0.016) | 61.81 | 74.90 (0.67) | 77.49 (0.94) |
| Hypertension | 1.038 | 1.031 (0.009) | 0.981 (0.012) | 1.100 | 1.090 (0.008) | 1.048 (0.014) | 66.88 | 81.62 (0.72) | 85.38 (1.03) |
| Rheumatoid Arthritis | 1.013 | 1.021 (0.007) | 0.993 (0.013) | 1.109 | 1.102 (0.008) | 1.076 (0.014) | 67.29 | 80.82 (0.71) | 84.23 (1.00) |
| Type I Diabetes | 1.015 | 1.024 (0.009) | 0.978 (0.015) | 1.144 | 1.122 (0.009) | 1.055 (0.015) | 67.63 | 82.29 (0.70) | 86.21 (1.05) |
| Type II Diabetes | 1.033 | 1.028 (0.008) | 0.993 (0.013) | 1.096 | 1.077 (0.009) | 1.032 (0.014) | 67.23 | 81.26 (0.71) | 84.82 (1.04) |
| Barrett's Oesophagus | 1.034 | 1.036 (0.007) | 1.011 (0.010) | 1.065 | 1.062 (0.007) | 1.037 (0.011) | 27.83 | 36.15 (0.28) | 41.81 (0.48) |
| Celiac Disease | 1.023 | 1.043 (0.009) | 0.978 (0.014) | 1.051 | 1.055 (0.009) | 0.994 (0.015) | 18.92 | 23.87 (0.21) | 26.92 (0.33) |
| Ischaemic Stroke | 1.075 | 1.071 (0.007) | 1.061 (0.011) | 1.165 | 1.159 (0.009) | 1.118 (0.012) | 44.70 | 61.08 (0.52) | 75.64 (1.03) |
| Parkinson's Disease | 0.999 | 1.007 (0.007) | 0.983 (0.011) | 1.041 | 1.020 (0.007) | 0.990 (0.011) | 23.66 | 30.80 (0.24) | 35.32 (0.37) |
| Psoriasis | 1.059 | 1.052 (0.008) | 1.010 (0.011) | 1.104 | 1.110 (0.008) | 1.056 (0.011) | 31.81 | 41.56 (0.33) | 48.88 (0.55) |
| Ulcerative Colitis | 1.045 | 1.056 (0.008) | 1.003 (0.012) | 1.082 | 1.079 (0.009) | 1.023 (0.012) | 30.69 | 41.23 (0.33) | 49.12 (0.61) |
| **Average** | **1.033** | **1.036 (0.002)** | **0.999 (0.003)** | **1.098** | **1.088 (0.002)** | **1.048 (0.004)** | **49.47** | **41.33 (0.11)** | **47.12 (0.17)** |

| Trait | Original Controls — $h^2_{\text{SNP}}$ (SD) | | | | 2 404 POBI Controls — $h^2_{\text{SNP}}$ (SD) | | | | 2 404 1000G Controls — $h^2_{\text{SNP}}$ (SD) | | | |
|---|---|---|---|---|---|---|---|---|---|---|---|---|
| | **LDSC-Zero** | **LDSC** | **SH-Zero** | **SH-GC** | **LDSC-Zero** | **LDSC** | **SH-Zero** | **SH-GC** | **LDSC-Zero** | **LDSC** | **SH-Zero** | **SH-GC** |
| Bipolar Disorder | 0.38 (0.05) | 0.24 (0.06) | 0.69 (0.07) | 0.97 (0.16) | 0.74 (0.05) | 0.21 (0.06) | 1.38 (0.08) | 0.57 (0.15) | 2324 (89) | 13.3 (4.4) | 1242 (13) | -0.25 (0.11) |
| Coronary Artery Disease | 0.21 (0.04) | 0.15 (0.05) | 0.33 (0.06) | 0.53 (0.13) | 0.54 (0.05) | 0.15 (0.05) | 0.99 (0.07) | 0.37 (0.13) | 2412 (93) | 15.7 (4.5) | 1289 (13) | -0.24 (0.11) |
| Crohn's Disease | 0.23 (0.05) | 0.06 (0.06) | 0.52 (0.07) | 0.53 (0.16) | 0.62 (0.06) | 0.06 (0.06) | 1.29 (0.09) | 0.64 (0.17) | 2228 (85) | 14.8 (4.2) | 1197 (12) | -0.11 (0.12) |
| Hypertension | 0.22 (0.04) | 0.09 (0.05) | 0.49 (0.07) | 0.67 (0.12) | 0.43 (0.05) | 0.05 (0.05) | 0.92 (0.07) | 0.47 (0.13) | 2343 (89) | 15.5 (4.5) | 1258 (13) | -0.23 (0.11) |
| Rheumatoid Arthritis | 0.08 (0.04) | -0.00 (0.04) | 0.26 (0.07) | 0.33 (0.14) | 0.46 (0.04) | 0.02 (0.04) | 1.04 (0.07) | 0.31 (0.14) | 2367 (89) | 16.1 (4.5) | 1268 (13) | -0.18 (0.11) |
| Type I Diabetes | 0.18 (0.04) | 0.09 (0.05) | 0.40 (0.07) | 0.61 (0.16) | 0.60 (0.05) | 0.05 (0.06) | 1.26 (0.08) | 0.72 (0.17) | 2366 (88) | 14.9 (4.5) | 1266 (13) | -0.24 (0.11) |
| Type II Diabetes | 0.24 (0.04) | 0.13 (0.05) | 0.48 (0.06) | 0.54 (0.14) | 0.53 (0.05) | 0.20 (0.05) | 1.05 (0.07) | 0.73 (0.15) | 2363 (90) | 14.9 (4.3) | 1258 (13) | -0.20 (0.11) |
| Barrett's Oesophagus | 0.17 (0.04) | 0.03 (0.04) | 0.42 (0.06) | 0.32 (0.11) | 0.28 (0.04) | 0.03 (0.05) | 0.64 (0.06) | 0.30 (0.11) | 838 (31) | 4.9 (1.7) | 561 (5) | -1.36 (0.10) |
| Celiac Disease | 0.18 (0.03) | 0.09 (0.03) | 0.36 (0.04) | 0.48 (0.09) | 0.19 (0.03) | 0.07 (0.03) | 0.40 (0.04) | 0.43 (0.09) | 320 (12) | 1.2 (0.7) | 184 (2) | -0.55 (0.06) |
| Ischaemic Stroke | 0.24 (0.03) | 0.01 (0.03) | 0.59 (0.04) | 0.14 (0.08) | 0.70 (0.03) | 0.10 (0.04) | 1.40 (0.05) | 0.46 (0.09) | 1172 (44) | 3.4 (2.3) | 778 (8) | -1.61 (0.09) |
| Parkinson's Disease | 0.18 (0.04) | 0.16 (0.05) | 0.34 (0.06) | 0.51 (0.13) | 0.24 (0.04) | 0.16 (0.05) | 0.48 (0.05) | 0.59 (0.12) | 718 (26) | 4.1 (1.5) | 483 (5) | -1.31 (0.10) |
| Psoriasis | 0.32 (0.04) | 0.11 (0.04) | 0.66 (0.07) | 0.56 (0.12) | 0.60 (0.05) | 0.13 (0.05) | 1.22 (0.07) | 0.68 (0.11) | 922 (34) | 4.0 (1.9) | 613 (6) | -1.46 (0.09) |
| Ulcerative Colitis | 0.26 (0.04) | 0.06 (0.04) | 0.57 (0.06) | 0.54 (0.11) | 0.39 (0.04) | 0.10 (0.04) | 0.81 (0.06) | 0.63 (0.11) | 820 (31) | 3.3 (1.6) | 550 (6) | -1.37 (0.09) |
| **Average** | **0.21 (0.01)** | **0.08 (0.01)** | **0.47 (0.02)** | **0.46 (0.03)** | **0.44 (0.01)** | **0.10 (0.01)** | **0.90 (0.02)** | **0.51 (0.03)** | **665 (9)** | **3.7 (0.5)** | **408 (1)** | **-0.76 (0.03)** |
| **Relative** | **1** | | | | **2.02 (0.05)** | | | | **3395 (43)** | | | |
| **Relative** | | **1** | | | | **0.99 (0.12)** | | | | **35.8 (5.1)** | | |
| **Relative** | | | **1** | | | | **1.95 (0.04)** | | | | **1019 (3)** | |
| **Relative** | | | | **1** | | | | **0.97 (0.07)** | | | | **-1.19 (0.05)** |

**Supplementary Table 7: Introducing population structure for the 13 WTCCC GWAS.** For each GWAS, we replace 2 404 randomly picked controls with either 2 404 individuals from People of the British Isles[11] (POBI) or the 2 404 individuals from the 1000 Genome Project[2] (1000G). The top table compares estimates of confounding bias, measured using the genomic inflation factor, the LDSC intercept ($1 + A$) or the SumHer-GC scaling factor ($C$); the bottom table compares estimates of $h^2_{\text{SNP}}$ from LDSC-Zero, LDSC, SumHer-Zero and SumHer-GC. Note that the values in the first block (the analyses prior to switching out controls) are slightly different to those in Supplementary Table 2, because here we restricted to SNPs common to the POBI and 1000G datasets. These results indicate that while SumHer-GC copes well with country-level population structure (that introduced by switching in POBI individuals), it fares poorly with continental-level structure (switching in 1000G individuals); however, as it is standard to identify and exclude ancestral outliers prior to performing association analysis,[12] in practice, it such severe confounding is unlikely to be encountered.

| Trait | Type | $\bar{n}_j$ | $m$ | MM | Correction of test statistics | GIF | GIF' | $1+A$ (SD) | $C$ (SD) |
|---|---|---|---|---|---|---|---|---|---|
| Alzheimer's Diseases[3] | Meta | 54162 | 3280621 | | GC within cohorts (average GIF 1.03) | 1.07 | 1.09 | 1.07 (0.01) | 1.03 (0.01) |
| Coronary Artery[27] | Meta | 79514 | 1218378 | | GC within cohorts (average GIF 1.06) | 1.07 | 1.11 | 1.07 (0.01) | 0.99 (0.01) |
| Crohn's Disease[28] | Mega | 20883 | 3018655 | | None | 1.08 | 1.14 | 1.10 (0.02) | 0.97 (0.02) |
| Depression[29] | Meta | 161460 | 3011263 | | Corrected using LDSC | 1.12 | 1.24 | 1.05 (0.01) | 0.97 (0.01) |
| Ever Smoked?[30] | Meta | 74053 | 1226914 | | GC within cohorts (average GIF 1.03) | 1.05 | 1.11 | 1.03 (0.01) | 0.96 (0.01) |
| Inflammatory Bowel[28] | Mega | 34652 | 3099673 | | None | 1.13 | 1.19 | 1.14 (0.02) | 0.98 (0.02) |
| Neuroticism[29] | Meta | 170911 | 3011181 | | Corrected using LDSC | 1.24 | 1.24 | 1.09 (0.01) | 0.88 (0.02) |
| Rheumatoid Arthritis[31] | Meta | 58284 | 3531676 | | GC within cohorts and after meta-analysis | 1.00 | 1.05 | 1.00 (0.01) | 0.89 (0.01) |
| Schizophrenia[26] | Meta | 81080 | 3262530 | | None | 1.35 | 1.61 | 1.24 (0.01) | 0.90 (0.01) |
| Subjective-Wellbeing[29] | Meta | 298420 | 1139680 | | Corrected using LDSC | 1.12 | 1.24 | 1.09 (0.01) | 0.93 (0.02) |
| Type 2 Diabetes[32] | Meta | 156633 | 3541873 | | GC within cohorts (average GIF unknown) | 1.08 | 1.15 | 1.08 (0.01) | 0.95 (0.01) |
| Ulcerative Colitis[28] | Mega | 27432 | 3142657 | | None | 1.09 | 1.13 | 1.11 (0.01) | 1.00 (0.01) |
| Bone Mineral Density[33] | Meta | 32965 | 3129002 | ✓ | None | 1.08 | 1.11 | 1.07 (0.01) | 1.00 (0.01) |
| Body Mass Index[34] | Meta | 228644 | 1255312 | | GC within cohorts and after meta-analysis | 0.92 | 1.13 | 0.82 (0.01) | 0.56 (0.02) |
| Fasting Glucose[35] | Meta | 58074 | 1277111 | | GC within cohorts (average GIF 1.06) | 1.04 | 1.09 | 1.05 (0.01) | 0.99 (0.01) |
| Glycated Hemoglobin[36] | Meta | 46368 | 1251236 | | GC within cohorts (average GIF 1.03) | 1.02 | 1.05 | 1.03 (0.01) | 1.00 (0.01) |
| HDL Cholesterol[37] | Meta | 95169 | 1228883 | ✓ | GC within cohorts and after meta-analysis | 0.95 | 1.03 | 1.00 (0.04) | 0.74 (0.02) |
| Height[5] | Meta | 244528 | 1251999 | | GC within cohorts (average GIF 1.03) | 1.69 | 2.09 | 1.73 (0.05) | 1.02 (0.03) |
| LDL Cholesterol[37] | Meta | 90473 | 1226496 | ✓ | GC within cohorts and after meta-analysis | 0.96 | 1.04 | 1.00 (0.04) | 0.74 (0.04) |
| Menarche Age[38] | Meta | 252514 | 3539601 | | GC within cohorts (average GIF 1.03) | 1.36 | 1.65 | 1.25 (0.02) | 0.91 (0.02) |
| Menopause Age[39] | Meta | 69360 | 1218477 | ✓ | Unclear whether GC was performed | 1.05 | 1.10 | 1.07 (0.02) | 0.93 (0.02) |
| Triglyceride Levels[37] | Meta | 91630 | 1226949 | ✓ | GC within cohorts and after meta-analysis | 0.96 | 1.03 | 0.94 (0.03) | 0.73 (0.03) |
| Waist-Hip Ratio[40] | Meta | 141432 | 1250319 | | GC within cohorts and after meta-analysis | 0.95 | 1.05 | 0.93 (0.01) | 0.77 (0.01) |
| Years Education[4] | Meta | 328917 | 3447867 | | GC within cohorts (average GIF 1.02) | 1.18 | 1.41 | 1.14 (0.01) | 0.84 (0.01) |
| **Average** | | **120732** | **2241181** | | | **1.11** | **1.21** | **1.06 (0.00)** | **0.93 (0.00)** |

**Supplementary Table 8: Details of the 24 summary GWAS.** Type reports whether the GWAS performed a mega-analysis (all individuals analyzed together) or meta-analysis (cohorts analyzed separately, then their results combined). $\bar{n}_j$ denotes average sample size; numbers in red mark the eight traits for which per-SNP sample sizes were available (else we set $n_j = n$, the total sample size). $m$ denotes the number of SNPs after filtering; we excluded SNPs not in our reference panel and, if possible, SNPs with info score <.95 (red numbers mark the five traits for which info scores were available). MM indicates traits where mixed-model association analysis was used for one or more analyses (this list is not exhaustive, as for some of the meta-analysis GWAS, it was unclear what analyses were used within cohorts). Column 6 summarizes any correction of test statistics, either using genomic control (GC) or dividing by the LDSC intercept. The 11 traits marked in red are those where correction was lowest; specifically, those where classical linear regression was performed, correction was only within cohorts, and the average correction was 1.03 or less (we decided to include bone mineral density even though mixed-model association analysis was performed, because classical linear regression was used for the majority of cohorts). GIF and GIF' are the genomic inflation factors computed from a thinned subset of SNPs (within 1 cM and $r^2_{jl} > 0.1$) and from all SNPs, respectively. $1 + A$ and $C$ are the estimates of the intercept from LDSC and scaling factor from SumHer-GC, respectively.

| Trait | log likelihood $L(S_j \| \widehat{h^2_{\text{SNP}}}, D^0)$ LDSC | SumHer-GC | LDSC-Zero | SumHer-GC | Proportion of LDAK $p$ (SD) Hybrid-GC | Hybrid-Zero |
|---|---|---|---|---|---|---|
| Alzheimer's Disease | -138159 | -138150 | -138195 | -138152 | 1.00 (0.10) | 1.01 (0.08) |
| Coronary Artery Disease | -114080 | -114058 | | | 1.00 (0.05) | |
| Crohn's Disease | -128896 | -128849 | -128939 | -128852 | 0.95 (0.07) | 0.94 (0.08) |
| Depression | -110431 | -110413 | | | 0.97 (0.05) | |
| Ever Smoked? | -110106 | -110090 | -110113 | -110094 | 0.88 (0.05) | 0.80 (0.08) |
| Inflammatory Bowel | -137434 | -137371 | -137517 | -137372 | 0.99 (0.05) | 0.99 (0.05) |
| Neuroticism | -120130 | -120008 | | | 0.94 (0.02) | |
| Rheumatoid Arthritis | -147348 | -147281 | | | 1.02 (0.03) | |
| Schizophrenia | -149884 | -149385 | -150376 | -149422 | 0.90 (0.01) | 0.89 (0.02) |
| Subjective-Wellbeing | -104209 | -104171 | | | 0.91 (0.04) | |
| Type 2 Diabetes | -159541 | -159460 | | | 0.98 (0.04) | |
| Ulcerative Colitis | -125174 | -125127 | -125249 | -125127 | 1.07 (0.04) | 1.08 (0.04) |
| Bone Mineral Density | -136558 | -136524 | -136605 | -136524 | 0.97 (0.04) | 0.97 (0.05) |
| Body Mass Index | -158031 | -157890 | | | 0.80 (0.02) | |
| Fasting Glucose | -137044 | -137038 | | | 0.92 (0.10) | |
| Glycated Hemoglobin | -111622 | -111618 | -111628 | -111618 | 0.98 (0.13) | 0.99 (0.12) |
| HDL Cholesterol | -164182 | -164140 | | | 0.82 (0.12) | |
| Height | -202174 | -201967 | -202725 | -201968 | 0.89 (0.03) | 0.89 (0.03) |
| LDL Cholesterol | -173559 | -173537 | | | 0.86 (0.13) | |
| Menarche Age | -205826 | -205487 | -206163 | -205511 | 0.85 (0.02) | 0.83 (0.02) |
| Menopause Age | -134492 | -134452 | | | 1.04 (0.06) | |
| Triglyceride | -173385 | -173379 | | | 0.68 (0.13) | |
| Waist-Hip Ratio | -117724 | -117607 | | | 0.88 (0.03) | |
| Years Education | -160262 | -159783 | -160542 | -159861 | 0.90 (0.01) | 0.87 (0.02) |
| **Average** | **-142510** | **-142408** | **-146186** | **-145864** | **0.91 (0.01)** | **0.90 (0.01)** |

**Supplementary Table 9: Comparing heritability models for the 24 summary GWAS using SumHer.** Columns 2 to 5 report the log likelihood $L(S_j|\widehat{h^2_{\text{SNP}}}, D^0)$ (see Supplementary Table 6), first calculated from LDSC and SumHer-GC, then, for the 11 traits least impacted by genomic control, calculated from LDSC-Zero and SumHer-Zero (the higher value in each pair is marked in red). The final two columns report estimates of $p$, the LDAK proportion, for each trait from Hybrid-GC and, for the 11 traits least impacted by genomic control, from Hybrid-Zero.

| Trait | SNPs $>$ 1 Mb OR $r^2 < 0.2$ Raw | GC | LDSC | SumHer-GC | SNPs $>$ 1 Mb OR $r^2 < 0.05$ Raw | GC | LDSC | SumHer-GC | SNPs $>$ 3 Mb OR $r^2 < 0.05$ Raw | GC | LDSC | SumHer-GC |
|---|---|---|---|---|---|---|---|---|---|---|---|---|
| Alzheimer's Disease | 22 | 20 | 20 | 21 | 19 | 17 | 17 | 18 | 19 | 17 | 17 | 18 |
| Coronary Artery Disease | 11 | 7 | 7 | 11 | 11 | 7 | 7 | 11 | 11 | 7 | 7 | 11 |
| Crohn's Disease | 67 | 61 | 60 | 68 | 55 | 51 | 50 | 56 | 55 | 51 | 50 | 56 |
| Depression | 1 | 0 | 1 | 1 | 1 | 0 | 1 | 1 | 1 | 0 | 1 | 1 |
| Ever Smoked? | 0 | 0 | 0 | 0 | 0 | 0 | 0 | 0 | 0 | 0 | 0 | 0 |
| Inflammatory Bowel | 72 | 61 | 60 | 76 | 60 | 51 | 51 | 62 | 60 | 51 | 51 | 62 |
| Neuroticism | 16 | 5 | 12 | 30 | 14 | 5 | 12 | 24 | 13 | 4 | 11 | 23 |
| Rheumatoid Arthritis | 196 | 196 | 196 | 216 | 71 | 71 | 71 | 81 | 70 | 70 | 70 | 80 |
| Schizophrenia | 96 | 38 | 49 | 133 | 82 | 35 | 44 | 108 | 82 | 34 | 43 | 108 |
| Subjective-Wellbeing | 1 | 0 | 0 | 2 | 1 | 0 | 0 | 2 | 1 | 0 | 0 | 2 |
| Type 2 Diabetes | 37 | 32 | 31 | 39 | 31 | 27 | 26 | 33 | 31 | 27 | 26 | 33 |
| Ulcerative Colitis | 38 | 34 | 32 | 38 | 30 | 28 | 27 | 30 | 30 | 28 | 27 | 30 |
| Bone Mineral Density | 22 | 21 | 22 | 22 | 16 | 16 | 16 | 16 | 16 | 16 | 16 | 16 |
| Body Mass Index | 76 | 88 | 114 | 286 | 60 | 70 | 92 | 239 | 60 | 70 | 92 | 239 |
| Fasting Glucose | 27 | 24 | 24 | 28 | 20 | 18 | 18 | 21 | 20 | 18 | 18 | 21 |
| Glycated Hemoglobin | 11 | 11 | 11 | 11 | 8 | 8 | 8 | 8 | 8 | 8 | 8 | 8 |
| HDL Cholesterol | 131 | 142 | 131 | 198 | 99 | 106 | 99 | 142 | 99 | 106 | 99 | 141 |
| Height | 814 | 322 | 310 | 787 | 580 | 242 | 234 | 563 | 580 | 242 | 234 | 563 |
| LDL Cholesterol | 128 | 135 | 128 | 183 | 83 | 89 | 83 | 117 | 83 | 89 | 83 | 117 |
| Menarche Age | 306 | 159 | 197 | 376 | 244 | 134 | 160 | 293 | 244 | 134 | 160 | 293 |
| Menopause Age | 45 | 39 | 38 | 53 | 36 | 31 | 30 | 42 | 36 | 31 | 30 | 42 |
| Triglyceride | 110 | 115 | 116 | 172 | 67 | 71 | 72 | 109 | 68 | 72 | 73 | 109 |
| Waist-Hip Ratio | 22 | 27 | 30 | 53 | 21 | 25 | 28 | 48 | 21 | 25 | 28 | 48 |
| Years Education | 170 | 99 | 99 | 191 | 151 | 88 | 88 | 166 | 151 | 88 | 88 | 166 |
| **Total** | **2419** | **1636** | **1688** | **2995** | **1760** | **1190** | **1234** | **2190** | **1759** | **1188** | **1232** | **2187** |
| **Average** | **100.8** | **68.2** | **70.3** | **124.8** | **73.3** | **49.6** | **51.4** | **91.2** | **73.3** | **49.5** | **51.3** | **91.1** |
| **Relative** | **1** | **0.68** | **0.70** | **1.24** | | | | | | | | |
| **Relative** | | | | | **1** | **0.68** | **0.70** | **1.24** | | | | |
| **Relative** | | | | | | | | | **1** | **0.68** | **0.70** | **1.24** |

**Supplementary Table 10: Number of significant loci for the 24 summary GWAS after correction for confounding bias.** Values report the number of independent loci with $P < 5 \times 10^{-8}$ (equivalently, $\chi^2(1)$ test statistic $> 29.72$), either based on the reported test statistics, or after correction using genomic control (dividing test statistics by the genomic inflation factor), LDSC (dividing them by the intercept $1 + A$) or SumHer-GC (dividing them by the scaling factor $C$). We consider increasingly strict definitions of independent: first we define two SNPs as independent if they are either >1 cM apart or have correlation-squared $> 0.2$; then as independent if either >1 cM apart or have correlation-squared $> 0.05$, and finally as independent if either >3 cM apart or have correlation-squared $> 0.05$.

| Category | LDSC (53-part model) Share (SD) | Expected | Enrichment (SD) | SumHer-GC (25-part model) Share (SD) | Expected | Enrichment (SD) |
|---|---|---|---|---|---|---|
| Coding | 0.119 (0.014) | 0.017 | 6.84 (0.81) | 0.030 (0.002) | 0.021 | 1.48 (0.07) |
| Conserved | 0.257 (0.024) | 0.033 | 7.75 (0.73) | 0.058 (0.002) | 0.036 | 1.60 (0.06) |
| CTCF | -0.033 (0.018) | 0.021 | -1.52 (0.88) | 0.025 (0.002) | 0.025 | 0.99 (0.06) |
| Digital Genomic Footprint | 0.295 (0.047) | 0.139 | 2.15 (0.34) | 0.200 (0.004) | 0.158 | 1.27 (0.02) |
| DNase I Hypersensitive Site | 0.251 (0.053) | 0.179 | 1.39 (0.29) | 0.241 (0.004) | 0.206 | 1.17 (0.02) |
| FANTOM5 Enhancer | 0.003 (0.008) | 0.004 | 1.01 (2.20) | 0.004 (0.001) | 0.005 | 0.89 (0.12) |
| Enhancer | 0.109 (0.021) | 0.036 | 3.03 (0.57) | 0.062 (0.002) | 0.047 | 1.30 (0.05) |
| Fetal DHS | 0.187 (0.040) | 0.087 | 2.09 (0.46) | 0.128 (0.003) | 0.105 | 1.22 (0.03) |
| H3K27ac (Hnisz) | 0.623 (0.020) | 0.358 | 1.74 (0.06) | 0.519 (0.005) | 0.437 | 1.19 (0.01) |
| H3K27ac (PGC2) | 0.480 (0.034) | 0.250 | 1.92 (0.13) | 0.375 (0.005) | 0.303 | 1.24 (0.02) |
| H3K4me1 | 0.852 (0.039) | 0.432 | 1.98 (0.09) | 0.602 (0.005) | 0.496 | 1.21 (0.01) |
| H3K4me3 | 0.307 (0.030) | 0.121 | 2.53 (0.25) | 0.198 (0.004) | 0.145 | 1.37 (0.03) |
| H3K9ac | 0.297 (0.030) | 0.111 | 2.67 (0.26) | 0.203 (0.004) | 0.143 | 1.42 (0.03) |
| Intronic | 0.485 (0.016) | 0.409 | 1.19 (0.04) | 0.470 (0.004) | 0.411 | 1.14 (0.01) |
| Promoter Flanking | -0.010 (0.011) | 0.008 | -1.23 (1.44) | 0.013 (0.001) | 0.009 | 1.51 (0.12) |
| Promoter | 0.053 (0.013) | 0.029 | 1.77 (0.46) | 0.047 (0.002) | 0.034 | 1.37 (0.06) |
| Repressed | 0.272 (0.039) | 0.473 | 0.58 (0.08) | 0.362 (0.005) | 0.451 | 0.80 (0.01) |
| Super Enhancer | 0.277 (0.013) | 0.140 | 1.98 (0.09) | 0.229 (0.004) | 0.190 | 1.20 (0.02) |
| Transcription Factor Binding Site | 0.299 (0.041) | 0.126 | 2.35 (0.32) | 0.185 (0.004) | 0.151 | 1.23 (0.02) |
| Transcribed | 0.482 (0.037) | 0.365 | 1.32 (0.10) | 0.413 (0.005) | 0.349 | 1.18 (0.01) |
| Transcription Start Site | 0.029 (0.012) | 0.013 | 2.15 (0.92) | 0.030 (0.002) | 0.018 | 1.70 (0.08) |
| 3’ Untranslated Region | 0.069 (0.010) | 0.013 | 5.47 (0.80) | 0.022 (0.001) | 0.016 | 1.34 (0.08) |
| 5’ Untranslated Region | 0.022 (0.008) | 0.006 | 3.81 (1.51) | 0.008 (0.001) | 0.006 | 1.37 (0.13) |
| Weak Enhancer | 0.063 (0.018) | 0.018 | 3.53 (1.02) | 0.030 (0.001) | 0.025 | 1.23 (0.06) |

**Supplementary Table 11: Average estimates of enrichment from the 24 summary GWAS.** We estimate enrichments using either LDSC with a 53-part model or SumHer-GC with a 25-part model. For the 25-part model, we divide the genome into a set for each of the 24 categories, plus a set containing all other SNPs, while for the 53-part model, we divide the genome into a set for each of the 24 categories, a set for each of 28 buffer regions (see Finucane *et al.*,[9]), plus a set containing all other SNPs. For each trait, we calculate the estimated enrichment of each category, obtained by dividing the estimated share of $h^2_{\text{SNP}}$ contributed by the category by its expected share; for each category, we then report the (inverse-weighted) average estimated enrichment across the 24 traits. Average estimated enrichments significantly different from 1 ($p < 0.05$) are marked in red.

| Category | LDSC-Zero (53-part model) Share (SD) | Expected | Enrichment (SD) | SumHer-Zero (25-part model) Share (SD) | Expected | Enrichment (SD) |
|---|---|---|---|---|---|---|
| Coding | 0.130 (0.021) | 0.02 | 7.89 (1.265) | 0.030 (0.002) | 0.020 | 1.53 (0.12) |
| Conserved | 0.135 (0.028) | 0.03 | 4.77 (0.965) | 0.060 (0.003) | 0.035 | 1.74 (0.09) |
| CTCF | -0.000 (0.022) | 0.02 | 0.06 (1.108) | 0.027 (0.002) | 0.025 | 1.08 (0.09) |
| Digital Genomic Footprint | 0.390 (0.059) | 0.13 | 2.93 (0.446) | 0.202 (0.006) | 0.157 | 1.30 (0.04) |
| DNase I Hypersensitive Site | 0.248 (0.061) | 0.17 | 1.45 (0.362) | 0.245 (0.006) | 0.203 | 1.22 (0.03) |
| FANTOM5 Enhancer | 0.026 (0.011) | 0.00 | 7.41 (3.138) | 0.006 (0.001) | 0.005 | 1.15 (0.18) |
| Enhancer | 0.130 (0.026) | 0.03 | 3.74 (0.736) | 0.068 (0.003) | 0.047 | 1.44 (0.07) |
| Fetal DHS | 0.235 (0.046) | 0.08 | 2.85 (0.564) | 0.130 (0.005) | 0.104 | 1.26 (0.04) |
| H3K27ac (Hnisz) | 0.684 (0.024) | 0.35 | 1.94 (0.068) | 0.556 (0.006) | 0.436 | 1.28 (0.01) |
| H3K27ac (PGC2) | 0.534 (0.039) | 0.25 | 2.15 (0.160) | 0.394 (0.007) | 0.302 | 1.31 (0.02) |
| H3K4me1 | 0.762 (0.047) | 0.42 | 1.82 (0.112) | 0.619 (0.007) | 0.494 | 1.26 (0.01) |
| H3K4me3 | 0.329 (0.035) | 0.12 | 2.73 (0.294) | 0.215 (0.006) | 0.144 | 1.50 (0.04) |
| H3K9ac | 0.272 (0.036) | 0.11 | 2.45 (0.332) | 0.219 (0.006) | 0.142 | 1.55 (0.04) |
| Intronic | 0.405 (0.022) | 0.41 | 0.99 (0.054) | 0.491 (0.006) | 0.411 | 1.19 (0.01) |
| Promoter Flanking | -0.017 (0.013) | 0.01 | -2.16 (1.677) | 0.014 (0.002) | 0.009 | 1.58 (0.19) |
| Promoter | 0.061 (0.017) | 0.03 | 2.06 (0.579) | 0.051 (0.003) | 0.034 | 1.52 (0.09) |
| Repressed | 0.265 (0.044) | 0.47 | 0.56 (0.094) | 0.336 (0.007) | 0.451 | 0.74 (0.02) |
| Super Enhancer | 0.316 (0.015) | 0.14 | 2.28 (0.110) | 0.243 (0.005) | 0.190 | 1.27 (0.03) |
| Transcription Factor Binding Site | 0.432 (0.055) | 0.12 | 3.42 (0.447) | 0.190 (0.006) | 0.149 | 1.29 (0.04) |
| Transcribed | 0.379 (0.042) | 0.37 | 1.03 (0.113) | 0.442 (0.007) | 0.350 | 1.26 (0.02) |
| Transcription Start Site | 0.061 (0.017) | 0.01 | 4.39 (1.225) | 0.039 (0.002) | 0.018 | 2.19 (0.14) |
| 3’ Untranslated Region | 0.053 (0.013) | 0.01 | 4.36 (1.094) | 0.023 (0.002) | 0.016 | 1.48 (0.13) |
| 5’ Untranslated Region | 0.043 (0.012) | 0.01 | 8.06 (2.330) | 0.010 (0.001) | 0.006 | 1.63 (0.20) |
| Weak Enhancer | 0.066 (0.021) | 0.02 | 3.73 (1.248) | 0.027 (0.002) | 0.024 | 1.12 (0.09) |

**Supplementary Table 12: Average estimates of enrichment from the 11 summary GWAS least impacted by genomic control.** Details are the same as Supplementary Table 11, except the averages are across the 11 traits least impacted by genomic control (see Supplementary Table 8), and instead of LDSC and SumHer-GC, estimates of enrichment are obtained using LDSC-Zero and SumHer-Zero, respectively.

| **Trait 1** | **Trait 2** | **LDSC Correlation (SD)** | **SumHer-GC Correlation (SD)** |
|---|---|---|---|
| Alzheimer's Disease | Years Education | -0.27 (0.08) | **-0.23 (0.05)** |
| Body Mass Index | Coronary Artery Disease | **0.22 (0.06)** | 0.14 (0.04) |
| Body Mass Index | Depression | **0.21 (0.05)** | 0.09 (0.04) |
| Body Mass Index | Ever Smoked? | **0.20 (0.05)** | **0.15 (0.04)** |
| Body Mass Index | Fasting Glucose | **0.35 (0.07)** | **0.27 (0.04)** |
| Body Mass Index | HDL Cholesterol | **-0.40 (0.10)** | **-0.25 (0.03)** |
| Body Mass Index | Menarche Age | **-0.38 (0.03)** | **-0.36 (0.02)** |
| Body Mass Index | Schizophrenia | **-0.10 (0.03)** | **-0.08 (0.02)** |
| Body Mass Index | Triglyceride | 0.26 (0.07) | **0.19 (0.04)** |
| Body Mass Index | Type 2 Diabetes | **0.59 (0.05)** | **0.43 (0.04)** |
| Body Mass Index | Waist-Hip Ratio | **0.67 (0.05)** | **0.55 (0.03)** |
| Body Mass Index | Years Education | **-0.27 (0.03)** | **-0.25 (0.02)** |
| Bone Mineral Density | Menarche Age | -0.08 (0.05) | **-0.15 (0.04)** |
| Coronary Artery Disease | HDL Cholesterol | **-0.43 (0.10)** | **-0.26 (0.04)** |
| Coronary Artery Disease | Height | -0.15 (0.06) | **-0.12 (0.03)** |
| Coronary Artery Disease | LDL Cholesterol | 0.23 (0.10) | **0.30 (0.05)** |
| Coronary Artery Disease | Triglyceride | **0.44 (0.08)** | **0.31 (0.05)** |
| Coronary Artery Disease | Type 2 Diabetes | **0.51 (0.09)** | **0.36 (0.06)** |
| Coronary Artery Disease | Waist-Hip Ratio | **0.28 (0.08)** | **0.24 (0.05)** |
| Coronary Artery Disease | Years Education | **-0.27 (0.06)** | **-0.25 (0.04)** |
| Crohn's Disease | Inflammatory Bowel | **0.95 (0.03)** | **0.87 (0.02)** |
| Crohn's Disease | Ulcerative Colitis | **0.75 (0.10)** | **0.57 (0.05)** |
| Depression | Neuroticism | **0.87 (0.06)** | **0.69 (0.04)** |
| Depression | Schizophrenia | **0.31 (0.06)** | **0.28 (0.05)** |
| Depression | Subjective-Wellbeing | **-0.78 (0.08)** | **-0.64 (0.06)** |
| Depression | Years Education | **-0.37 (0.06)** | **-0.37 (0.05)** |
| Ever Smoked? | Triglyceride | 0.13 (0.07) | **0.15 (0.04)** |
| Ever Smoked? | Years Education | **-0.31 (0.05)** | **-0.29 (0.04)** |
| Fasting Glucose | Type 2 Diabetes | **0.60 (0.11)** | **0.66 (0.09)** |
| Glycated Hemoglobin | Type 2 Diabetes | **0.69 (0.16)** | **0.63 (0.10)** |
| Glycated Hemoglobin | Waist-Hip Ratio | 0.49 (0.15) | **0.33 (0.08)** |
| HDL Cholesterol | Menarche Age | **0.15 (0.04)** | **0.13 (0.02)** |
| HDL Cholesterol | Triglyceride | **-0.87 (0.11)** | **-0.56 (0.05)** |
| HDL Cholesterol | Type 2 Diabetes | **-0.43 (0.08)** | **-0.37 (0.04)** |
| HDL Cholesterol | Waist-Hip Ratio | **-0.57 (0.11)** | **-0.37 (0.04)** |
| HDL Cholesterol | Years Education | **0.22 (0.04)** | **0.17 (0.02)** |
| Height | Menarche Age | **0.14 (0.03)** | **0.14 (0.02)** |
| Height | Triglyceride | **-0.15 (0.04)** | -0.07 (0.02) |
| Height | Years Education | **0.13 (0.03)** | **0.11 (0.02)** |
| Inflammatory Bowel | Schizophrenia | **0.17 (0.05)** | 0.11 (0.03) |
| Inflammatory Bowel | Ulcerative Colitis | **0.94 (0.03)** | **0.91 (0.02)** |
| LDL Cholesterol | Triglyceride | **0.41 (0.10)** | **0.36 (0.05)** |
| Menarche Age | Triglyceride | -0.12 (0.04) | **-0.11 (0.03)** |
| Menarche Age | Type 2 Diabetes | **-0.24 (0.04)** | **-0.19 (0.03)** |
| Menarche Age | Waist-Hip Ratio | **-0.25 (0.04)** | **-0.18 (0.03)** |
| Menopause Age | Years Education | 0.16 (0.06) | **0.12 (0.03)** |
| Neuroticism | Schizophrenia | **0.20 (0.05)** | 0.11 (0.04) |
| Neuroticism | Subjective-Wellbeing | **-0.68 (0.05)** | **-0.60 (0.04)** |
| Neuroticism | Years Education | **-0.27 (0.04)** | **-0.27 (0.04)** |
| Rheumatoid Arthritis | Years Education | **-0.25 (0.05)** | **-0.21 (0.03)** |
| Schizophrenia | Subjective-Wellbeing | **-0.29 (0.05)** | **-0.26 (0.04)** |
| Schizophrenia | Ulcerative Colitis | **0.23 (0.06)** | 0.09 (0.04) |
| Triglyceride | Type 2 Diabetes | **0.36 (0.07)** | **0.35 (0.06)** |
| Triglyceride | Waist-Hip Ratio | **0.48 (0.08)** | **0.37 (0.05)** |
| Triglyceride | Years Education | **-0.19 (0.04)** | **-0.16 (0.03)** |
| Type 2 Diabetes | Waist-Hip Ratio | **0.66 (0.07)** | **0.52 (0.04)** |
| Type 2 Diabetes | Years Education | **-0.24 (0.05)** | **-0.21 (0.03)** |
| Waist-Hip Ratio | Years Education | **-0.33 (0.03)** | **-0.31 (0.03)** |

**Supplementary Table 13: Estimates of genetic correlation for the 24 summary GWAS.** Of the ${}^{24}C_2 = 276$ pairs of traits we considered, this table reports genetic correlations for the 58 pairs with significant correlation ($P < 0.05/276$) from either LDSC or SumHer-GC. Nominally significant estimates ($P < 0.05$) are marked in red, while Bonferonni significant estimates ($P < 0.05/276$) are also in **bold**.

| Polygenic Risk Score | $h^2_{\text{SNP}}$ | Clump | BMI | Height | HDL | LDL | TG | Mean (SD) | Relative |
|---|---|---|---|---|---|---|---|---|---|
| **Classical** | | YES | 0.221 | 0.189 | 0.156 | 0.051 | 0.118 | **0.156 (0.003)** | **0.936 (0.020)** |
| **Classical** | | NO | 0.228 | 0.195 | 0.151 | 0.061 | 0.168 | **0.167 (0.004)** | 1 |
| **Bayesian: GCTA Model** | $\widehat{h^2_{\text{SNP}}}$ | YES | 0.210 | 0.197 | 0.166 | 0.067 | 0.172 | **0.169 (0.003)** | **0.989 (0.020)** |
| **Bayesian: GCTA Model** | 0.5 | YES | 0.226 | 0.196 | 0.159 | 0.053 | 0.156 | **0.166 (0.003)** | **0.992 (0.019)** |
| **Bayesian: GCTA Model** | $\widehat{h^2_{\text{SNP}}}$ | NO | 0.174 | 0.178 | 0.101 | 0.061 | 0.149 | **0.137 (0.004)** | **0.816 (0.020)** |
| **Bayesian: GCTA Model** | 0.5 | NO | 0.213 | 0.182 | 0.131 | 0.061 | 0.162 | **0.156 (0.004)** | **0.931 (0.020)** |
| **Bayesian: Enriched GCTA** | $\widehat{h^2_{\text{SNP}}}$ | YES | 0.228 | 0.204 | 0.168 | 0.070 | 0.158 | **0.173 (0.003)** | **1.023 (0.019)** |
| **Bayesian: Enriched GCTA** | 0.5 | YES | 0.234 | 0.199 | 0.158 | 0.065 | 0.157 | **0.171 (0.003)** | **1.016 (0.019)** |
| **Bayesian: Enriched GCTA** | $\widehat{h^2_{\text{SNP}}}$ | NO | 0.194 | 0.172 | 0.108 | 0.062 | 0.167 | **0.147 (0.004)** | **0.867 (0.020)** |
| **Bayesian: Enriched GCTA** | 0.5 | NO | 0.215 | 0.183 | 0.134 | 0.062 | 0.167 | **0.159 (0.004)** | **0.945 (0.020)** |
| **Bayesian: LDAK Model** | $\widehat{h^2_{\text{SNP}}}$ | YES | 0.242 | 0.201 | 0.183 | 0.066 | 0.157 | **0.178 (0.003)** | **1.058 (0.020)** |
| **Bayesian: LDAK Model** | 0.5 | YES | 0.237 | 0.198 | 0.176 | 0.062 | 0.145 | **0.171 (0.004)** | **1.025 (0.020)** |
| **Bayesian: LDAK Model** | $\widehat{h^2_{\text{SNP}}}$ | NO | 0.192 | 0.182 | 0.096 | 0.064 | 0.149 | **0.144 (0.004)** | **0.852 (0.020)** |
| **Bayesian: LDAK Model** | 0.5 | NO | 0.221 | 0.184 | 0.131 | 0.064 | 0.162 | **0.160 (0.004)** | **0.950 (0.020)** |
| **Bayesian: Enriched LDAK** | $\widehat{h^2_{\text{SNP}}}$ | YES | 0.242 | 0.204 | 0.194 | 0.052 | 0.166 | **0.180 (0.004)** | **1.075 (0.020)** |
| **Bayesian: Enriched LDAK** | 0.5 | YES | 0.238 | 0.201 | 0.181 | 0.065 | 0.153 | **0.175 (0.004)** | **1.043 (0.020)** |
| **Bayesian: Enriched LDAK** | $\widehat{h^2_{\text{SNP}}}$ | NO | 0.192 | 0.184 | 0.100 | 0.063 | 0.155 | **0.145 (0.004)** | **0.863 (0.020)** |
| **Bayesian: Enriched LDAK** | 0.5 | NO | 0.221 | 0.186 | 0.134 | 0.067 | 0.165 | **0.162 (0.004)** | **0.960 (0.020)** |

**Supplementary Table 14: Comparing the predictive performance of Classical and Bayesian PRS.** Each PRS uses a model of the form $\sum_j \beta_j X_j$, where the vector $X_j$ contains genotypes for SNP $j$, and the $\beta_j$ are trained using the relevant set of summary statistics from the 24 summary GWAS (either body mass index, HDL cholesterol, height, LDL cholesterol or triglycerides). Values report correlation between predicted and observed phenotypes for the eMerge data (which are independent of the 24 summary GWAS). For the Classical PRS, the $\beta_j$ are estimates from classical linear regression. For the Bayesian PRS, the $\beta_j$ are posterior means, obtained using one of four prior distributions: GCTA Model, Enriched GCTA, LDAK and Enriched LDAK (the enriched versions incorporate average estimates of enrichments for the 24 functional categories). For the Bayesian PRS, it is necessary to provide a value for $h^2_{\text{SNP}}$; we either used the corresponding estimate from LDSC-Zero or SumHer-Zero (for Enriched GCTA we used a 53-part model, for Enriched LDAK we used a 25-part model), or agnostically set $h^2_{\text{SNP}} = 0.5$. When constructing PRS, it can be beneficial to clump[41] (identify pairs of SNPs within 1 cM with $r^2_{jl} > 0.05$, then discard the one with highest $p$-value / lowest Bayes Factor); we found that clumping was detrimental for the Classical PRS, but always benefited the Bayesian PRS. The PRS marked in red are those reported in the main text.

| Raw GWAS | Number of loci (MHC) | Summary GWAS | Number of loci (MHC) |
|---|---|---|---|
| Crohn's Disease | 5 (0) | Alzheimer's Disease | 5 (0) |
| Rheumatoid Arthritis | 5 (5) | Crohn's Disease | 2 (0) |
| Type 1 Diabetes | 8 (5) | Rheumatoid Arthritis | 4 (4) |
| Celiac Disease | 7 (5) | Bone Mineral Density | 1 (0) |
| Psoriasis | 5 (5) | HDL Cholesterol | 1 (0) |
| Ulcerative Colitis | 1 (1) | Triglyceride Levels | 1 (0) |
| Age-related Macular Disease | 1 (0) | | |
| HDL Cholesterol | 1 (0) | | |

**Supplementary Table 15: Numbers of large-effect loci for the 25 raw and 24 summary GWAS.** For all analyses, we exclude SNPs within the major histocompatibility complex (Chromosome 6: 25-34 Mb), as well as SNPs which individually explain >1% of phenotypic variation, and SNPs in LD with these. This table reports for each GWAS the number of large-effect SNPs after thinning (within 1 cM and $r^2_{jl} > 0.05$), and how many of these are in the MHC.

1. Juster, F. & Suzman, R. An overview of the Health and Retirement Study. *J. Hum. Resources* **30**, S7–S56 (1995).
2. The 1000 Genomes Project Consortium. A map of human genome variation from population-scale sequencing. *Nature* **467**, 1061–1073 (2010).
3. Lambert, J. *et al.* Meta-analysis of 74,046 individuals identifies 11 new susceptibility loci for Alzheimer's disease. *Nat. Genet.* **45**, 1452–1458 (2013).
4. Okbay, A. *et al.* Genome-wide association study identifies 74 loci associated with educational attainment. *Nature* **533**, 539–542 (2016).
5. Wood, A. *et al.* Defining the role of common variation in the genomic and biological architecture of adult human height. *Nat. Genet.* **46**, 1173–1186 (2014).
6. Speed, D. *et al.* Reevaluation of SNP heritability in complex human traits. *Nat. Genet.* **49**, 986–992 (2017).
7. The Wellcome Trust Case Control Consortium. Genome-wide association study of 14,000 cases of seven common diseases and 3,000 shared controls. *Nature* **447**, 661–678 (2007).
8. Bulik-Sullivan, B. *et al.* LD score regression distinguishes confounding from polygenicity in genome-wide association studies. *Nat. Genet.* **47**, 291–295 (2014).
9. Finucane, H. *et al.* Partitioning heritability by functional annotation using genome-wide association summary statistics. *Nat. Genet.* **47**, 1228–1235 (2015).
10. Speed, D. *et al.* Describing the genetic architecture of epilepsy through heritability analysis. *Brain* **137**, 26802689 (2014).
11. Leslie, S. *et al.* The fine-scale genetic structure of the British population. *Nature* **519**, 309–314 (2015).
12. Anderson, C. *et al.* Data quality control in genetic case-control association studies. *Nat. Protoc.* **5**, 1564–1573 (2010).
13. Yu, J. *et al.* A unified mixed-model method for association mapping that accounts for multiple levels of relatedness. *Nat. Genet.* **38**, 203–208 (2006).
14. Astle, W. & Balding, D. Population structure and cryptic relatedness in genetic association studies. *Statist. Sci.* **24**, 451–471 (2009).
15. The International HapMap 3 Consortium. Integrating common and rare genetic variation in diverse human populations. *Nature* **467**, 52–58 (2010).
16. Kent, W. *et al.* The human genome browser at ucsc. *Genome Res.* **12**, 996–1006 (2002).
17. Gusev, A. *et al.* Partitioning Heritability of Regulatory and Cell-Type-Specific Variants across 11 Common Diseases. *Am. J. Hum. Genet.* **95**, 535–552 (2014).
18. Lindblad-Toh, K. *et al.* A high-resolution map of human evolutionary constraint using 29 mammals. *Nature* **478**, 476–482 (2011).
19. Ward, L. & Kellis, M. Evidence of abundant purifying selection in humans for recently acquired regulatory functions. *Science* **337**, 1675–1678 (2012).
20. Hoffman, M. *et al.* Integrative annotation of chromatin elements from encode data. *Nucleic Acids Res.* **41**, 827–841 (2013).
21. The ENCODE Project Consortium. An integrated encyclopedia of dna elements in the human genome. *Nature* **489**, 57–74 (2012).
22. Roadmap Epigenomics Consortium *et al.* Integrative analysis of 111 reference human epigenomes. *Nature* **518**, 317–330 (2015).
23. Trynka, G. *et al.* Chromatin marks identify critical cell types for fine mapping complex trait variants. *Nat. Genet.* **45**, 124–130 (2013).
24. Andersson, R. *et al.* An atlas of active enhancers across human cell types and tissues. *Nature* **507**, 455–461 (2014).
25. Hnisz, D. *et al.* Super-enhancers in the control of cell identity and disease. *Cell* **155**, 934–947 (2013).

26. Schizophrenia Working Group of the Psychiatric Genomics Consortium. Biological insights from 108 schizophrenia-associated genetic loci. *Nature* **511**, 421–427 (2014).

27. Schunkert, H. *et al.* Large-scale association analysis identifies 13 new susceptibility loci for coronary artery disease. *Nat. Genet.* **43**, 333–338 (2011).

28. Liu, J. *et al.* Association analyses identify 38 susceptibility loci for inflammatory bowel disease and highlight shared genetic risk across populations. *Nat. Genet.* **47**, 979–986 (2015).

29. Okbay, A. *et al.* Genetic variants associated with subjective well-being, depressive symptoms, and neuroticism identified through genome-wide analyses. *Nat. Genet.* **48**, 626–633 (2016).

30. The Tobacco and Genetics Consortium. Genome-wide meta-analyses identify multiple loci associated with smoking behavior. *Nat. Genet.* **42**, 441–447 (2010).

31. Okada, Y. *et al.* Genetics of rheumatoid arthritis contributes to biology and drug discovery. *Nature* **506**, 376–381 (2014).

32. Scott, R. *et al.* An expanded genome-wide association study of type 2 diabetes in Europeans. *Diabetes* **66**, 2888–2902 (2017).

33. Zheng, H. *et al.* Whole-genome sequencing identifies EN1 as a determinant of bone density and fracture. *Nature* **526**, 112–117 (2015).

34. Locke, A. *et al.* Genetic studies of body mass index yield new insights for obesity biology. *Nature* **518**, 197–206 (2015).

35. Manning, A. *et al.* A genome-wide approach accounting for body mass index identifies genetic variants influencing fasting glycemic traits and insulin resistance. *Nat. Genet.* **44**, 659–669 (2012).

36. Soranzo, N. *et al.* Common variants at 10 genomic loci influence hemoglobin a(c) levels via glycemic and nonglycemic pathway. *Diabetes* **59**, 3229–3239 (2010).

37. Global Lipids Genetics Consortium. Discovery and refinement of loci associated with lipid levels. *Nat. Genet.* **45**, 1274–1283 (2013).

38. Perry, J. *et al.* Parent-of-origin-specific allelic associations among 106 genomic loci for age at menarche. *Nature* **514**, 92–97 (2014).

39. Day, F. *et al.* Large-scale genomic analyses link reproductive aging to hypothalamic signaling, breast cancer susceptibility and brca1-mediated dna repair. *Nat. Genet.* **47**, 1294–1303 (2015).

40. Shungin, D. *et al.* New genetic loci link adipose and insulin biology to body fat distribution. *Nat. Genet.* **518**, 187–196 (2015).

41. Euesden, J., Lewis, C. & O'Reilly, P. PRSice: polygenic risk score software. *Bioinformatics* **31**, 1466–1468 (2015).
